# CRISPR-TSKO facilitates efficient cell type-, tissue-, or organ-specific mutagenesis in Arabidopsis

**DOI:** 10.1101/474981

**Authors:** Ward Decaestecker, Rafael Andrade Buono, Marie L. Pfeiffer, Nick Vangheluwe, Joris Jourquin, Mansour Karimi, Gert Van Isterdael, Tom Beeckman, Moritz K. Nowack, Thomas B. Jacobs

## Abstract

Detailed functional analyses of many fundamentally-important plant genes via conventional loss-of-function approaches are impeded by severe pleiotropic phenotypes. In particular, mutations in genes that are required for basic cellular functions and/or reproduction often interfere with the generation of homozygous mutant plants, precluding further functional studies. To overcome this limitation, we devised a CRISPR-based tissue-specific knockout system, CRISPR-TSKO, enabling the generation of somatic mutations in particular plant cell types, tissues, and organs. In Arabidopsis, CRISPR-TSKO mutations in essential genes caused well-defined, localized phenotypes in the root cap, stomatal lineage, or entire lateral roots. The underlying modular cloning system allows for efficient selection, identification, and functional analysis of mutant lines directly in the first transgenic generation. The efficacy of CRISPR-TSKO opens new avenues to discover and analyze gene functions in spatial and temporal contexts of plant life while avoiding pleiotropic effects of system-wide loss of gene function.

## Introduction

The generation of stable, inheritable loss-of-function mutant alleles has been indispensable for functional genomic studies in plants. Such knockout or knockdown lines have been generated with various techniques such as ionizing radiation, ethyl methanesulfonate, T-DNA or transposon insertions, RNA interference (RNAi), or artificial microRNAs. In recent years there has been an explosion in the generation of knockout plant lines by clustered regularly interspaced short palindromic repeat (CRISPR) technology.

CRISPR technology contains two components, the CRISPR-associated (Cas) nuclease and CRISPR RNAs (crRNA) that direct the nuclease to the target nucleic acid. The most commonly used CRISPR system in plants is based on the CRISPR-associated 9 (Cas9) DNA endonuclease and its artificial crRNA, the guide RNA (gRNA) (Jinek et al., 2012). In plants, Cas9 is very efficient at inducing double-strand DNA breaks. DNA breaks repaired by the error-prone non-homologous end joining pathway ultimately result in the formation of short insertions and/or deletions (indels) at the break site (Bortesi and Fischer, 2015). These indels most often lead to frame shifts and/or early stop codons, effectively generating knockout mutations in the targeted gene(s).

Most CRISPR efforts in plants to date have focused on generating stable and inheritable mutant alleles for reverse genetics approaches. Yet this approach is limited as the knockout of many fundamentally-important genes convey severe pleiotropic phenotypes up to lethality. Of the approximately 25,000 protein-coding genes in the *Arabidopsis thaliana* genome, 10% are estimated to be indispensable (Lloyd et al., 2015). This presents a considerable challenge for functional analyses of genes with essential functions.

An approach to overcome these problems is the use of tissue-specific gene silencing (Alvarez et al., 2006; Schwab et al., 2006). However, gene silencing is often incomplete, interfering with the interpretation of the observed phenotypes. Furthermore, it has been well established that small RNAs can be mobile (Melnyk et al., 2011), limiting the tissue specificity in gene-silencing experiments. Therefore, the results obtained using gene silencing are often not comparable with the use of stably transmitted DNA-based mutants.

Transgenic vectors generating dominant-negative protein versions have been developed for certain genes. Expressing these mutant versions in a tissue-specific context can locally interfere with endogenous gene functions (Fukaki et al., 2005; Mitsuda et al., 2011). Other methods include the conditional knockout of genes in specific cell types or tissues by using a CRE-recombinase (Sieburth et al., 1998). These approaches, however, can be cumbersome and difficult to scale (Munoz-Nortes et al., 2017).

Outside of the plant field, researchers have recently overcome such limitations with the development of conditional knockouts using CRISPR technology. In zebrafish, the *gata1* promoter driving Cas9 expression was used to knockout genes specifically in the erythrocytic lineage (Ablain et al., 2015). In Drosophila, targeted knockout mutations in two essential genes *Wingless* and *Wntless* only in germ cells permitted the generation of adult flies, whereas ubiquitous knockout individuals did not survive past the pupal stage (Port et al., 2014). Additionally, cardiomyocyte-specific expression of Cas9 led to organ-specific knockout in a mouse model (Carroll et al., 2016). The use of tissue-specific promoters to drive Cas9 expression have been reported in plants (Hyun et al., 2015; Yan et al., 2015; Mao et al., 2016). However, these efforts have been done to increase the recovery of stably-transmitted mutant alleles. Recently, the fiber-specific *NST3/SND1* promoter was used to drive Cas9 expression and target the essential gene *hydroxycinnamoyltransferase* in Arabidopsis (Liang et al., 2019). This allowed the authors to specifically decrease lignin in xylem cells while avoiding the strong pleiotropic growth defects in full knockout mutants.

Herein, we describe the development of a CRISPR **t**issue-**s**pecific **k**nock**o**ut (CRISPR-TSKO) vector system in Arabidopsis that allows for the specific generation of somatic DNA mutations in plants. The CRISPR-TSKO toolset is simple to use, highly efficient and allows for multiplexing and large-scale screening approaches. We show the potential of CRISPR-TSKO for somatic gene knockouts of essential genes in diverse plant cell types, tissues, and organs. We also detail important considerations and limitations on the use of CRISPR-TSKO and provide best practices for researchers. Our approach opens new opportunities to study the function of fundamentally-important genes in specific contexts of plant development and creates new possibilities to investigate post-embryonic developmental processes.

## Results

### Proof-of-concept: tissue-specific GFP knockout in the lateral root cap

We reasoned that by using tissue-specific, somatic promoters to drive Cas9 expression, CRISPR could be used to generate cell type-, tissue-, and organ-specific DNA mutations in plants. To test this hypothesis, T-DNA vectors were constructed with Cas9 expression controlled by the promoter region of *SOMBRERO*/*ANAC033* (*SMB*; AT1G79580). The *SMB* promoter (*pSMB*) is highly root cap-specific and activated directly after the formative division of root cap stem cells (Willemsen et al., 2008; Fendrych et al., 2014). A *pSMB*:Cas9 expression cassette was combined with one of two gRNAs targeting the *GFP* coding sequence, GFP-1 and GFP-2, and transformed into a homozygous Arabidopsis line with ubiquitous expression of a nuclear-localized GFP and β-glucuronidase (GUS) fusion protein (*pHTR5:NLS-GFP-GUS* (Ingouff et al., 2017), henceforth, NLS-GFP). Primary transgenic plants (T1 seedlings) were selected via resistance to the herbicide glufosinate and investigated for loss of GFP signal in the root tips of five-day-old seedlings. Six out of eleven *pSMB*:Cas9;GFP-1 events and three out of ten *pSMB*:Cas9;GFP-2 events showed an almost complete loss of GFP specifically in the root cap, suggesting CRISPR-mediated knockout soon after the formative division of the root cap stem cells (**Supplementary File 1A**). All other root tissues maintained GFP expression, indicating that Cas9 activity specifically in the root cap cells led to cell-autonomous *GFP* knockout. The tissue-specific knockout phenotype (*de novo* generation of mutations) was heritable, as T2 progeny from three lines with *pSMB*:Cas9;GFP-1 and three lines with *pSMB*:Cas9;GFP-2 had no GFP fluorescence in root cap cells while having normal NLS-GFP expression in all other tissues examined (**Supplementary File 1B**). These results clearly indicated that the use of a tissue-specific promoter driving Cas9 can efficiently induce somatic, tissue-specific knockout phenotypes.

### Design of the CRISPR-TSKO gene knockout toolset

To facilitate a wide range of future gene-modification approaches in an easy-to-use cloning system, we devised CRISPR-TSKO, a modular and versatile vector toolset based on Golden Gate technology and modified GreenGate vectors (Engler et al., 2008; Lampropoulos et al., 2013). CRISPR-TSKO is inexpensive and immediately compatible with GreenGate modules already in use by other laboratories. The modularity allows for the combination of Cas9, or any nuclease, with virtually any promoter sequence of choice. Furthermore, Cas9 fusion proteins are possible on the N- and C-termini, allowing for the wide range of CRISPR technologies such as base editors (Marzec and Hensel, 2018) or transcriptional regulators (Lowder et al., 2015). The promoter, Cas9, N- and C-tags, and terminator modules can be combined with an “unarmed” gRNA cassette to generate an unarmed destination vector (Figure. 1). One or two gRNAs can be directly cloned into this destination vector with a single Golden Gate reaction (see methods). Alternatively, when an AarI linker is used instead of the unarmed gRNA cassette, a second round of Golden Gate assembly can be performed for the cloning of up to 12 gRNAs in a single destination vector (**Supplementary File 2**).

**Figure 1:**
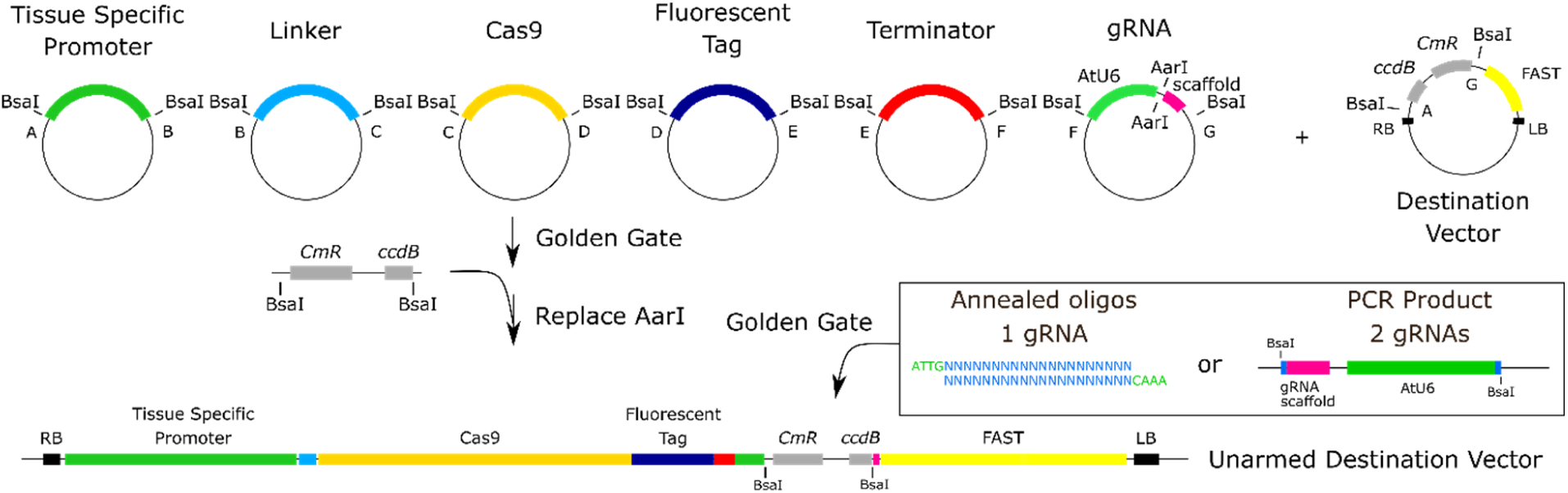
Cloning workflow for CRISPR-TSKO vectors. Six entry modules are combined in a binary destination vector, containing a FAST screenable marker cassette, with Golden Gate assembly. The six entry modules contain a tissue specific promoter, a cloning linker, the Cas9 nuclease, a fluorescent tag, a terminator and a module containing an *AtU6-26* promoter driving the expression of an unarmed gRNA scaffold. These modules replace the *ccd*B and *Cm*R selectable markers allowing for the negative selection of the destination vector in *ccd*B-sensitive *E. coli* cells. The resulting vector can be directly ‘armed’ with one or two gRNAs, upon pre-digestion with AarI. Alternatively, the AarI restriction sites can be replaced by a PCR product containing two BsaI sites flanking *ccd*B and *Cm*R expression cassettes. In a single Golden Gate reaction, a pair of annealed oligonucleotides are cloned, resulting in an expression vector containing one gRNA. Alternatively, Golden Gate cloning of a PCR product containing a first gRNA attached to an *AtU6-26* promoter and the protospacer sequence of the second gRNA, results in an expression vector containing two gRNAs.

A collection of binary destination vectors containing different selectable markers and/or non-destructive fluorescent markers based on the fluorescence-accumulating seed technology (FAST) system (Shimada et al., 2010) were generated to take advantage of this general cloning strategy (**Supplementary File 3**). The FAST system allows for the antibiotic- or herbicide-free selection of transformed T1 seeds and permits screening for phenotypes directly in T1 seedlings. To facilitate the evaluation of tissue specificity and expression levels of Cas9, a nuclear-localized fluorescent mCherry tag was fused to the *Cas9* coding sequence via a P2A ribosomal skipping peptide (Cermak et al., 2017). Using this Cas9-P2A-mCherry expression cassette (henceforth, Cas9-mCherry) we targeted different tissue types, cell lineages, and organs in Arabidopsis to explore the potential of CRISPR-TSKO for plant research.

### Root-cap specific gene knockout

To confirm the functionality of our new vector system, the expression of Cas9-mCherry was controlled by *pSMB* and combined with the gRNA GFP-1. Ten of the 21 T1 seedlings showed a loss of GFP fluorescence specifically in the root cap, while six were chimeric (partial loss of GFP) and five maintained normal GFP expression (Figure 2A, Table 1). We observed a delay in the onset of the knockout phenotype as cells of the youngest root cap layers had overlapping signals of GFP and mCherry (Figure 2B). This suggests that a certain time for mRNA and/or protein turnover of GFP is required after the onset of Cas9 expression for the knockout phenotype to become apparent. We observed a clear correlation between the intensity of mCherry signal and the penetrance of the knockout phenotype; all ten highly-expressing mCherry lines were entirely devoid of GFP signal in the root-cap (except for the youngest cells), the medium mCherry-expressing lines had chimeric knockout phenotypes and the low-to-no mCherry lines had chimeric or full expression of GFP (Table 1). By comparing the intensity of both fluorescent proteins in individual root cap nuclei, we confirmed that highly expressing Cas9 lines have a significantly higher probability of gene knockout (Figure 2C).

**Figure 2:**
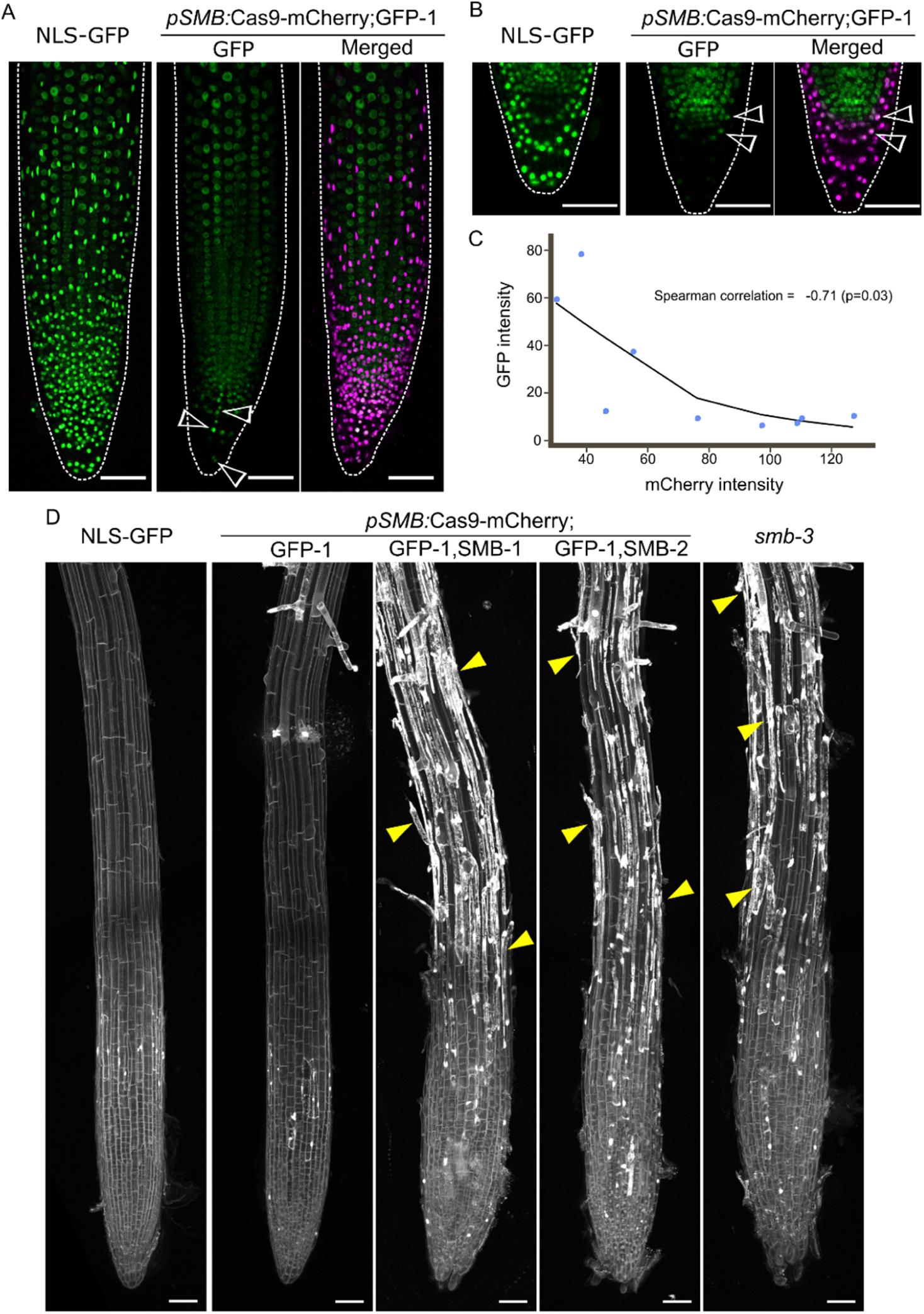
Root-cap specific knockout with CRISPR-TSKO. **A**, Maximum intensity projection of representative seedling of NLS-GFP and T1 of *pSMB:*Cas9-mCherry;GFP-1 with absence of GFP and presence of Cas9-mCherry signal specific to root cap cells. GFP is shown in green and Cas9-mCherry in magenta. Arrowheads indicate a patch of root cap cells in which GFP knockout was not achieved (chimera). **B**, Mid-section of root tip of both NLS-GFP and T1 seedling of *pSMB:*Cas9-mCherry;GFP-1. Arrowheads show young root cap cells in which GFP signal can still be observed. **C**, Plot of median intensity of root cap nuclei for both GFP and Cas9-mCherry in T1 seedlings. Line shows a Loess regression curve. **D**, Overview of root tips of 6 DAG T2 seedlings for both gRNAs for *SMB* displaying the characteristic cell corpse accumulation at the root surface (yellow arrowheads) with propidium iodide staining. All scale bars represent 50 µm.

**Table 1.** Phenotypes of T1 seedlings transformed with *pSMB*:Cas9-mCherry;GFP-1

| GFP Signal |  |  |  |
| --- | --- | --- | --- |
| mCherry | No | Chimeric | Normal |
| High | 10 | 0 | 0 |
| Medium | 0 | 3 | 0 |
| Low/No | 0 | 3 | 5 |
| Total seedlings | 21 |  |  |

To test if a root-cap expressed gene, the NAC transcription factor *SMB* itself, could be successfully targeted by CRISPR-TSKO, the gRNA GFP-1 was combined with one of two different gRNAs targeting *SMB* (SMB-1 and −2) with Cas9 expression driven by *pSMB*. Loss of *SMB* delays root cap maturation and preparation of programmed cell death in root cap cells, causing larger root caps and a delayed and aberrant root cap cell death with a lack of cell corpse clearance (Bennett et al., 2010; Fendrych et al., 2014). In T1 seedlings, we found *smb* mutant phenotypes for both SMB-1 and −2 coupled with the disappearance of root cap GFP signal (**Supplementary File 4A**). Both gRNAs appear equally effective as 13 out of 21 and 9 out of 12 T1 events gave clear simultaneous *smb* and *GFP* knockout phenotypes, respectively (Table 2, **Supplementary File 4A**). Knockout phenotypes were scored in four segregating lines in the T2 generation. The *smb* and *GFP* knockout phenotypes were observed in all FAST-positive T2 seedlings, whereas all FAST-negative seedlings (null segregants) showed no knockout phenotypes (Figure 2D**, Supplementary File 4B,** Table 3). These data demonstrate that CRISPR-TSKO-induced mutations are strictly somatic when using *pSMB* and that the mutagenic effect is heritable.

**Table 2.** Phenotypes of T1 seedlings transformed with *pSMB*:Cas9-mCherry;GFP-1,SMB-1 and *pSMB*:Cas9-mCherry;GFP-1,SMB-2

|  |  | GFP Signal |  |  | *smb* Phenotype |  |  |
| --- | --- | --- | --- | --- | --- | --- | --- |
| Vector | mCherry | No | Chimeric | Normal | Yes | Weak | No |
| GFP-1, *SMB*-1 | High | 12 (57%) | 0 | 0 | 11 (52%) | 1 (5%) | 0 |
|  | Medium | 3 (14%) | 0 | 0 | 1 (5%) | 2 (10%) | 0 |
|  | Low | 0 | 0 | 2 (10%) | 0 | 0 | 2 (10%) |
|  | No/Very low | 1 (5%) | 1 (5%) | 2 (10%) | 1 (5%) | 0 | 3 (14%) |
| GFP-1, *SMB*-2 | High | 8 (67%) | 0 | 0 | 7 (58%) | 0 | 1 (8%) |
|  | Medium | 0 | 0 | 0 | 0 | 0 | 0 |
|  | Low | 0 | 3 (25%) | 1 (8%) | 2 (17%) | 0 | 2 (17%) |
|  | No/Very low | 0 | 0 | 0 | 0 | 0 | 0 |

**Table 3.** Segregating phenotypes in T2 *pSMB*:Cas9-mCherry;GFP-1,SMB-1 and *pSMB*:Cas9-mCherry;GFP-1,SMB-2

| T1 Line | FAST | n | mCherry |  | GFP Signal |  | smb Phenotype |  |  |
| --- | --- | --- | --- | --- | --- | --- | --- | --- | --- |
|  |  |  | + | - | No | Normal | Yes | Weak | No |
| GFP-1,SMB-1<br>Line 2 | + | 34 | 34 | 0 | 34 | 0 | 33 | 1 | 0 |
|  | - | 30 | 0 | 30 | 0 | 30 | 0 | 0 | 30 |
| GFP-1,SMB-1<br>Line 16 | + | 21 | 21 | 0 | 21 | 0 | 19 | 2 | 0 |
|  | - | 30 | 0 | 30 | 0 | 30 | 0 | 0 | 30 |
| GFP-1,SMB-2<br>Line 12 | + | 30 | 30 | 0 | 30 | 0 | 29 | 1 | 0 |
|  | - | 25 | 0 | 25 | 0 | 25 | 0 | 0 | 25 |
| GFP-1,SMB-2<br>Line 5 | + | 26 | 26 | 0 | 26 | 0 | 25 | 1 | 0 |
|  | - | 33 | 0 | 33 | 0 | 33 | 0 | 0 | 33 |

To determine if the observed phenotypes were due to root cap-specific DNA mutations, protoplasts were prepared from root tips of four independent T2 lines (two for each *SMB* target) and used for fluorescence-activated cell sorting (FACS). DNA was extracted from sorted populations and the *SMB* and *GFP* target loci were PCR amplified and Sanger sequenced. TIDE analysis (Brinkman et al., 2014) was performed to determine the frequency and type of knockout alleles generated. The mCherry-positive populations (Cas9 expressing) had indel frequencies (TIDE Score) >95% for the GFP-1, SMB-1 and SMB-2 target loci (**Supplementary File 5**). In contrast, the mCherry-negative cell populations had indel frequencies of 1-5%, which are equivalent to wild-type or background levels when using TIDE analysis. The alleles generated were largely consistent across events, with 1-bp insertions being the predominant outcome (58-94%) followed by 1-bp deletions (3-38%; **Supplementary File 5**) for the GFP-1 target locus. A small, but significant, proportion of alleles were in-frame (3-bp deletions), but, as the GFP-1 gRNA targets the essential residue Gly67 (Fu et al., 2015), these alleles likely result in no GFP fluorescence. For the two *SMB*-targeting gRNAs, 1-bp insertions were the predominant repair outcome (78-88%) and a minority (∼10%) of alleles being 1-bp deletions for SMB-1 and 3-bp deletions for SMB-2 (**Supplementary File 5**). Thus the consistent *GFP* and *SMB* knockout phenotypes observed are due to an active and heritable Cas9-induced somatic mutagenesis specifically in root cap cells.

### Cell-lineage-specific gene knockout in the stomatal lineage

To test the possibility of using CRISPR-TSKO in a different somatic context, we utilized two promoter elements active in the stomatal cell lineage. The promoters of *TOO MANY MOUTHS* (*TMM*; AT1G80080) and *FAMA* (AT3G24140) control gene expression in the stomatal lineage, with *pTMM* expressing early in the lineage (Nadeau and Sack, 2002) and *pFAMA* expressing later during the formation of guard mother cells and young guard cells (Ohashi-Ito and Bergmann, 2006). These two promoters were used to produce CRISPR-TSKO constructs simultaneously targeting *GFP* and *PHYTOENE DESATURASE 3* (*PDS3*; AT4G14210) in the stomatal lineage as they should give clear knockout phenotypes. PDS3 is essential for chlorophyll, carotenoid and gibberellin biosynthesis. Null mutants show a dwarfed and albino phenotype and cannot survive on soil (Qin et al., 2007). Consistent with this, the ubiquitously-expressed Cas9-mCherry (*pPcUbi*:Cas9-mCherry;GFP-1,PDS3) gave rise to the expected severe phenotypic effects ranging from full albino to variegated leaves and stunted plants (Figure 3A**, Supplementary File 6A**).

**Figure 3:**
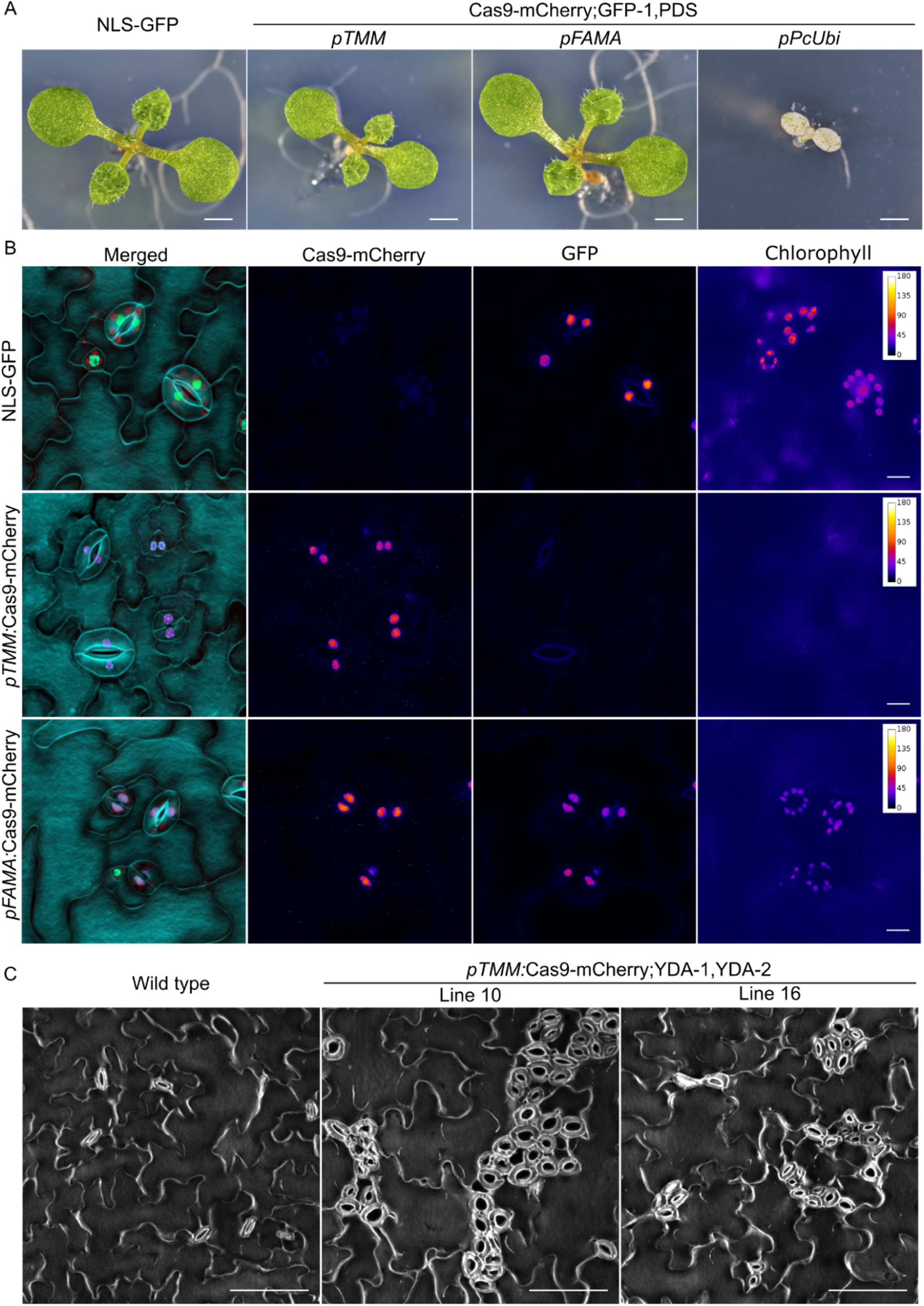
Stomatal-lineage specific knockout with CRISPR-TSKO. **A**, 9 DAG seedlings showing the partial rescue when *PDS3* is knocked out only in stomatal lineage (*pTMM*) in comparison with the arrested albino seedlings of ubiquitous knockout (*pPcUbi*). Scale bars represent 1 mm. **B**, Simultaneous stomata-lineage specific knockout of GFP and chlorophyll biosynthesis in 5 DAG T1 seedlings. Shown are stomata at the abaxial face of cotyledons. While both GFP and chlorophyll signals are lost in stomata in lines under the control of *pTMM*, signal is still present in stomata in lines under the control of *pFAMA*. In merged image, cell outlines are in cyan, GFP in green, Cas9-mCherry in magenta, and chlorophyll fluorescence in red. Epidermal cell patterning is shown using DAPI staining. Scale bars represent 10 µm. **C**, Targeting of *YDA* only in stomatal lineage (*pTMM*) is sufficient to cause clustering of stomata. Clusters of stomata are shown on the adaxial face of cotyledons of 15 DAG T2 seedlings. Scale bars represent 100 µm.

Five days after germination, cotyledons of T1 seedlings were assessed for chlorophyll and GFP fluorescence by epifluorescence microscopy. Eighteen out of 20 *pTMM:*Cas9-mCherry;GFP-1,PDS-1 T1 seedlings were clearly lacking both GFP and chlorophyll fluorescence (Figure 3B), indicative of successful knockouts in both genes. In a separate experiment, two out of 23 T1 seedlings did exhibit some mild bleaching similar to the ubiquitous knockout events (**Supplementary File 6A**), suggesting that *pTMM* can drive Cas9-mCherry expression in mesophyll cells at a low frequency. Independent T2 *pTMM*:Cas9 plants were generally smaller than the NLS-GFP background line (**Supplementary File 7**), but were otherwise not affected in vegetative and reproductive development. Thus, restricting the loss of PDS3 to the stomatal lineage did not markedly affect plant development.

In contrast to the high frequency of *GFP* and *PDS3* knockout phenotypes in *pTMM*:Cas9-mCherry;GFP-1;PDS3 events, we observed neither a loss of GFP nor chlorophyll fluorescence in 21 mCherry-expressing *pFAMA* T1 seedlings evaluated (Figure 3B). We hypothesized that the later induction of Cas9 by *pFAMA* allowed for residual *PDS3* and *GFP* mRNA and/or protein to persist in the targeted cells and these pools would have to be depleted before a loss of signal could be observed. Therefore, we investigated cotyledons ten days after germination in five *pTMM*:Cas9 and eight *pFAMA*:Cas9 T1 events. Despite this extended cultivation time, mCherry-positive guard cells still showed clear GFP and chlorophyll fluorescence signals in the *pFAMA*:Cas9 lines (**Supplementary File 6B).**

To determine if DNA mutations were induced in both the *pTMM*:Cas9 and *pFAMA*:Cas9 lines, protoplasts were prepared from T2 cotyledons of two independent lines for each genotype and sorted for mCherry. Surprisingly, mCherry-positive cells from one *pFAMA*:Cas9 line (13652.12) showed a reduced GFP fluorescence intensity during cell sorting (**Supplementary File 8C**). Genotyping results of the mCherry-positive and - negative protoplast populations determined an indel frequency of ∼80% for the *GFP* and *PDS3* target loci for the *pTMM*:Cas9 lines and 30-74% for the *pFAMA:*Cas9 lines (**Supplementary File 9**). The indel spectra for the *pTMM*:Cas9 line showed a preference for the 1-bp insertion. While the *pFAMA*:Cas9 lines have the same preference for the 1-bp insertion, they also have a greater variety of alleles from 3-bp deletions to 2-bp insertions (**Supplementary File 9**).

The detection of mutations in both *pFAMA*:Cas9 lines and the reduced GFP intensity detected by flow cytometry in one line was surprising given that a reduction of GFP signal was not observed by microscopy. To rule out technical errors, a second sorting experiment was performed on the two previously-sorted *pFAMA*:Cas9 T2 lines plus two additional lines. From these four lines, we clearly observed indel frequencies of 23-75% for *GFP* and 34-86% for *PDS3* (**Supplementary File 10**). Again, line 1365.12 had the highest indel frequencies and a reduction of GFP intensity (**Supplementary File 11).** These results indicate that DNA mutations were generated in the mCherry-expressing cells for both CRISPR-TSKO constructs, but do not resolve why knockout phenotypes were not observed in *pFAMA*:Cas9 lines.

To test if CRISPR-TSKO can be used to manipulate cell fate decisions within the stomatal lineage, we targeted *YODA* (*YDA*; AT1G63700), a mitogen-activated protein kinase kinase kinase. Knockout mutants of *YDA* have clustered stomata, severe developmental defects, frequent seedling growth arrest and, if *yda* mutants do manage to survive until flowering, sterility (Bergmann et al., 2004; Lukowitz et al., 2004). When we targeted *YDA* with a pair of gRNAs in a single, ubiquitously-expressed Cas9 construct (*pPcUbi*:Cas9-mCherry;YDA-1,YDA-2), 33 out of 35 mCherry-positive T1 seedlings contained clustered stomata on the cotyledonary epidermis to varying degrees of severity (**Supplementary File 12A**). Eight out of 19 T1 plants transferred to soil were sterile, consistent with the strong pleiotropic effects observed in reported *yda* mutants. When *YDA* was targeted by *pTMM*:Cas9-mCherry;YDA-1,YDA-2, all 40 T1 mCherry-positive seedlings had a clustered-stomata phenotype similar to that of the *yda-1* mutant, yet without the corresponding growth arrest (Figure 3C**, Supplementary File 12C**). All 19 plants transferred to soil developed similarly to wild-type and were fertile.

PCR and DNA sequence analysis confirmed efficient mutagenesis of *YDA* in T1 seedlings transformed with both *pPcUbi*:Cas9 and *pTMM*:Cas9 vectors. As a pair of gRNAs target *YDA*, an 813-bp deletion can be expected by the excision of the intervening DNA sequence. Such deletion events were observed in events transformed with both vectors (**Supplementary File 12B**). Indel frequencies for *pPcUbi*:Cas9 events were higher than *pTMM*:Cas9 as is expected for ubiquitous versus stomata-specific targeting (**Supplementary File 13**). These results illustrate that by utilizing the stomatal lineage-specific *pTMM*, we are able to uncouple the pleiotropic growth defects and sterility in systemic *YDA* knockouts (Lukowitz et al., 2004), but still retain the characteristic clustered-stomata phenotype.

### Organ-specific gene knockout in lateral roots

Next to gene-knockout in particular tissues and cell lineages, we tested the potential of CRISPR-TSKO to generate mutant organs on otherwise wild-type plants. To this end we made use of the promoter sequence of *GATA23,* a gene that marks the onset of lateral root organogenesis and is expressed in pericycle cells primed to become involved in lateral root formation in Arabidopsis (De Rybel et al., 2010). *GATA23* expression is transient and disappears prior to the emergence of the primordium from the primary root, except for some remaining expression at the base of the primordium (De Rybel et al., 2010) (Figure 4A). When targeting *GFP* with *pGATA23:*Cas9-mCherry;GFP-1, 20 out of 23 mCherry-positive T1 seedlings showed a complete or partial loss of GFP fluorescence in lateral roots while maintaining normal GFP expression in the primary root (Figure 4A,B, Table 4). In contrast, lines with undetectable mCherry expression showed chimeric or normal GFP expression in lateral roots (Table 4). Sequence analysis of lateral roots from six independent knockout events confirmed >93% of the alleles were mutated in those organs (**Supplementary File 14**). The indel spectrum was similar as the other tissue types, with the 1-bp insertion being the dominant repair outcome (**Supplementary File 14**).

**Figure 4:**
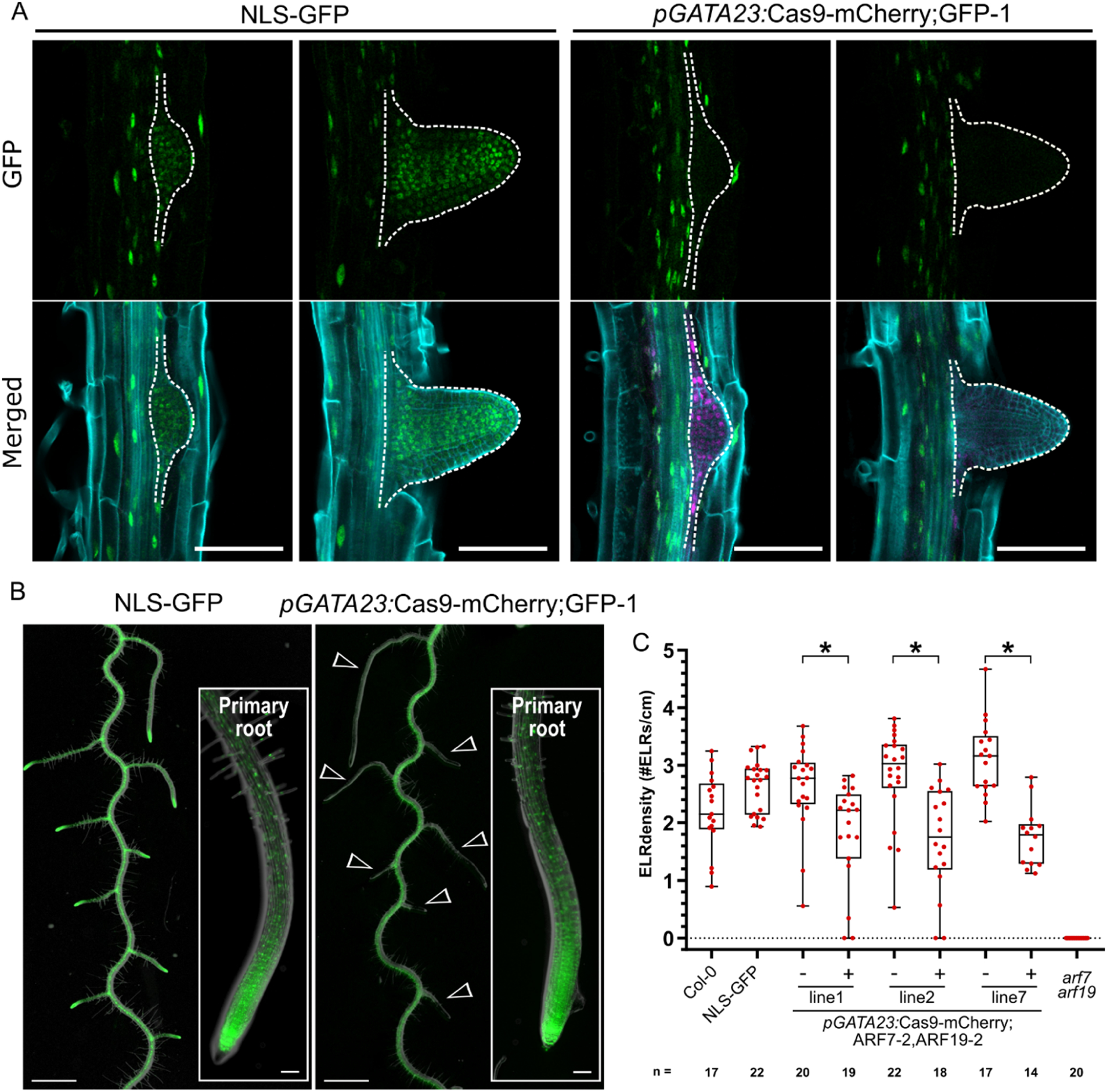
Organ-specific gene knockout using *pGATA23*-CRISPR-TSKO. **A,** Specific knockout of the GFP signal in emerging lateral roots (dashed outline). Representative images of the NLS-GFP control and *pGATA23*:Cas9-mCherry;GFP-1. GFP in green, mCherry in magenta, and cell wall stained with calcofluor white displayed in cyan. Scale bars represent 100 µm. **B,** GFP knockout is specific to lateral roots. Overlay of root morphology and GFP signal is shown for a representative NLS-GFP control and a T2 *pGATA23*:Cas9-mCherry;GFP-1 seedling. Arrowheads indicate GFP negative lateral roots. Insets are the tip of primary roots. Scale bars represent 1 mm for overview and 100µm for inset. **C,** Quantification of the emerged lateral root (ELR) density for Col-0 and FAST-positive (+) and –negative (-) T2 seedlings of *pGATA23*-CRISPR-TSKO lines targeting *ARF7* and *ARF19* simultaneously. ELR density was compared between FAST positive and negative seedlings within each line via Poisson regression analyses. * indicates *p* values smaller than 2×10^−3^.

**Table 4.** GFP phenotype in lateral roots of *pGATA23*:Cas9-mCherry;GFP-1

| GFP signal in LR |  |  |  |
| --- | --- | --- | --- |
| mCherry | No | Chimeric | Normal |
| Positive | 15 | 5 | 3 |
| Negative | 0 | 3 | 27 |
| Total seedlings | 15 | 8 | 30 |

Knockout phenotypes were scored in three segregating lines in the T2 generation. For two lines, all FAST-positive plants had no GFP expression in the lateral roots, while in line 3, 15 out of 17 plants had no GFP expression in lateral roots (Table 5). Together, these experiments demonstrate that organ-specific gene knockout in lateral roots is highly efficient via the xylem-pole pericycle-expressed Cas9 controlled by *pGATA23*.

**Table 5.**
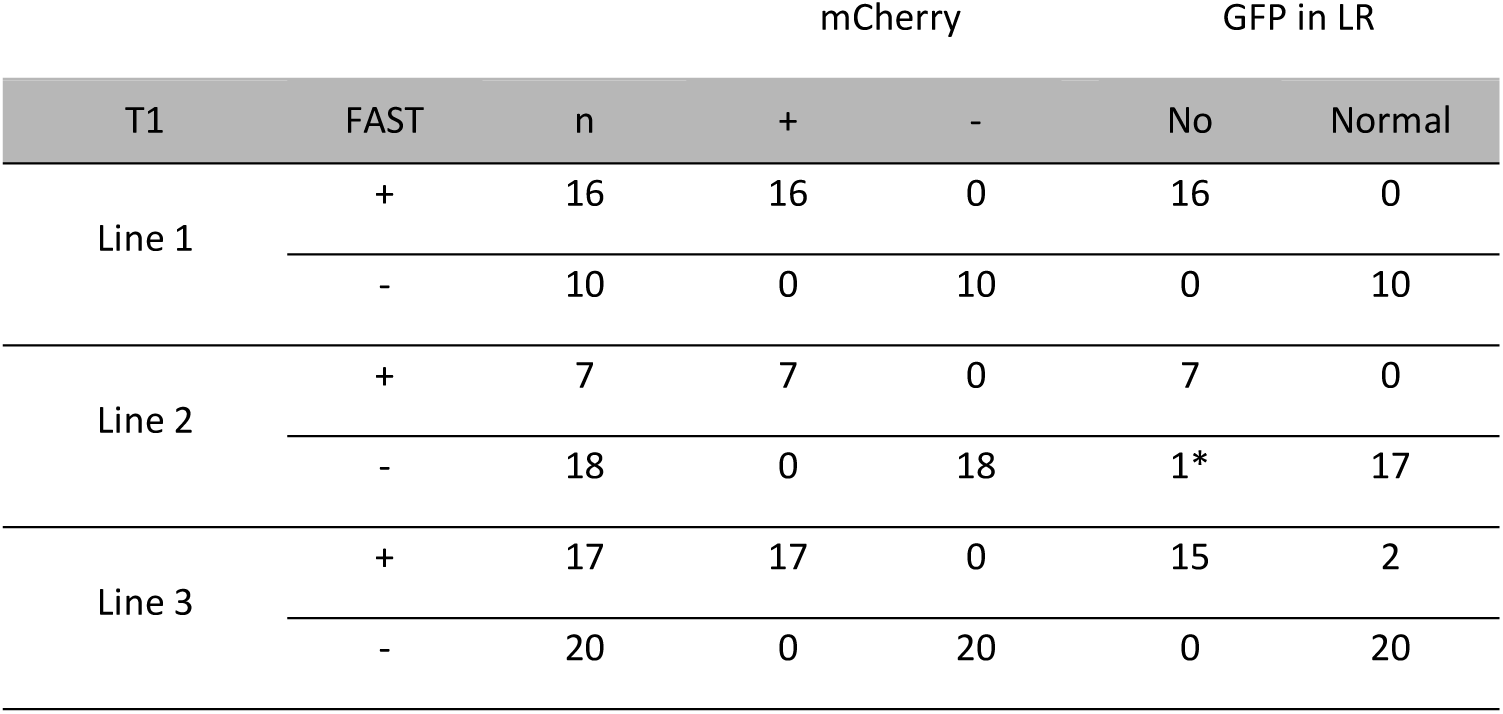
Phenotypic analysis of T2 seedlings of *pGATA23*:Cas9-mCherry;GFP-1. Plant indicated with an asterisk (*) showed no GFP signal in entire plant.

Lateral root organogenesis depends on the partially redundant action of *AUXIN RESPONSE FACTOR (ARF) 7* (AT5G20730) and *ARF19* (AT1G19220) as lateral root initiation is strongly inhibited in *arf7arf19* double-knockout mutants (Okushima et al., 2007). As both *ARFs* are broadly expressed in Arabidopsis seedlings, it is unclear whether this phenotype depends on ARF7 and ARF19 function strictly in xylem-pole pericycle cells (Okushima et al., 2005). To test this hypothesis, we used CRISPR-TSKO with *pGATA23* to target both *ARF7* and *ARF19*. We first recapitulated the ubiquitous double-knockout line *arf7arf19* with two ubiquitously-expressed Cas9 constructs each containing two different gRNAs targeting both *ARF*s (*pPcUbi*:Cas9-mCherry;ARF7-1,ARF19-1 and *pPcUbi*:Cas9-mCherry;ARF7-2,ARF19-2). While, no obvious reduction in lateral root density was observed in the T1 plants containing the first construct (*pPcUbi*:Cas9-mCherry;ARF7-1,ARF19-1), 18 out of 26 T1 plants containing the second construct (*pPcUbi*:Cas9-mCherry;ARF7-2,ARF19-2) completely lacked lateral roots (**Supplementary File 15A**) which is consistent with the phenotype of *arf7arf19* seedlings (Okushima et al., 2005). In agreement with these phenotyping results, sequencing of the target loci in whole roots showed that the *pPcUbi*:Cas9-mCherry;ARF7-1,ARF19-1 construct was particularly ineffective as the ARF19-1 target locus had an indel frequency of only 9-13% (**Supplementary File 16**), explaining the lack of a mutant phenotype in those T1s. In comparison, the indel frequencies were >93% for most events with the *pPcUbi*:Cas9-mCherry;ARF7-2,ARF19-2 construct (**Supplementary File 16**).

Emerged lateral root density was quantified in three segregating T2 lines transformed with *pGATA23*:Cas9-mCherry;ARF7-2,ARF19-2. Slight but significant reductions in emerged lateral root density was observed in FAST-positive T2 plants (Figure 4C **and Supplementary File 15B**). As these results were inconsistent with those of the ubiquitously-expressed construct, we sequenced the *ARF7 and ARF19* target loci in lateral roots of at least three plants per line. TIDE analysis revealed indel frequencies of >83% for ARF7-2 and >92% for ARF19-2 for nine out of ten plants and no wild-type alleles for either gene could be detected in lateral roots of four plants (**Supplementary File 17)**. Thus lateral root initiation is only mildly affected when *ARF7* and *ARF19* are knocked out in *GATA23*-expressing pericycle cells.

The central cell cycle regulator *CYCLINE-DEPENDENT KINASE A1* (*CDKA;1*; AT3G48750) is homologous to *CDK1* and *CDK2* in mammals and cell proliferation in *cdka;1* null mutants is severely affected (Nowack et al., 2012). Mutant embryos are superficially normal in appearance, but only contain a fraction of the number of cells that make up the wild-type embryo. Mutant seedlings are not viable on soil, but can be cultivated as sterile dwarf plants without a root system in axenic liquid cultures (Nowack et al., 2012). We generated CRISPR-TSKO constructs to specifically knockout *CDKA;1* in the lateral root primordia to allow us to study the effect of CDKA;1 in the context of lateral root formation. We started by testing the efficiency of our gRNAs using ubiquitously-expressed Cas9 with a paired-gRNA construct (*pPcUBI*:Cas9-mCherry;CDKA1-1,CDKA1-2). T1 seedlings reproduced the reported dwarf-seedling phenotype (Nowack et al., 2012) and genotyping revealed a 171-bp deletion, corresponding to the excision of the intervening DNA sequence (**Supplementary File 18**). TIDE analysis of the upper band showed higher indel frequencies with the gRNA CDKA1-1 (48-99%) than for gRNA CDKA1-2 (16-79%; **Supplementary File 18**). The same gRNAs were also used with a *pGATA23* construct. All T1 transgenic plants grew normally and were fertile. In T2, we were surprised that lateral root development was not as severely affected as anticipated; lateral root density was unaffected and lateral roots of FAST-positive T2 plants were 62-69% the length of their null-segregant siblings (Figure 5A, C).

**Figure 5:**
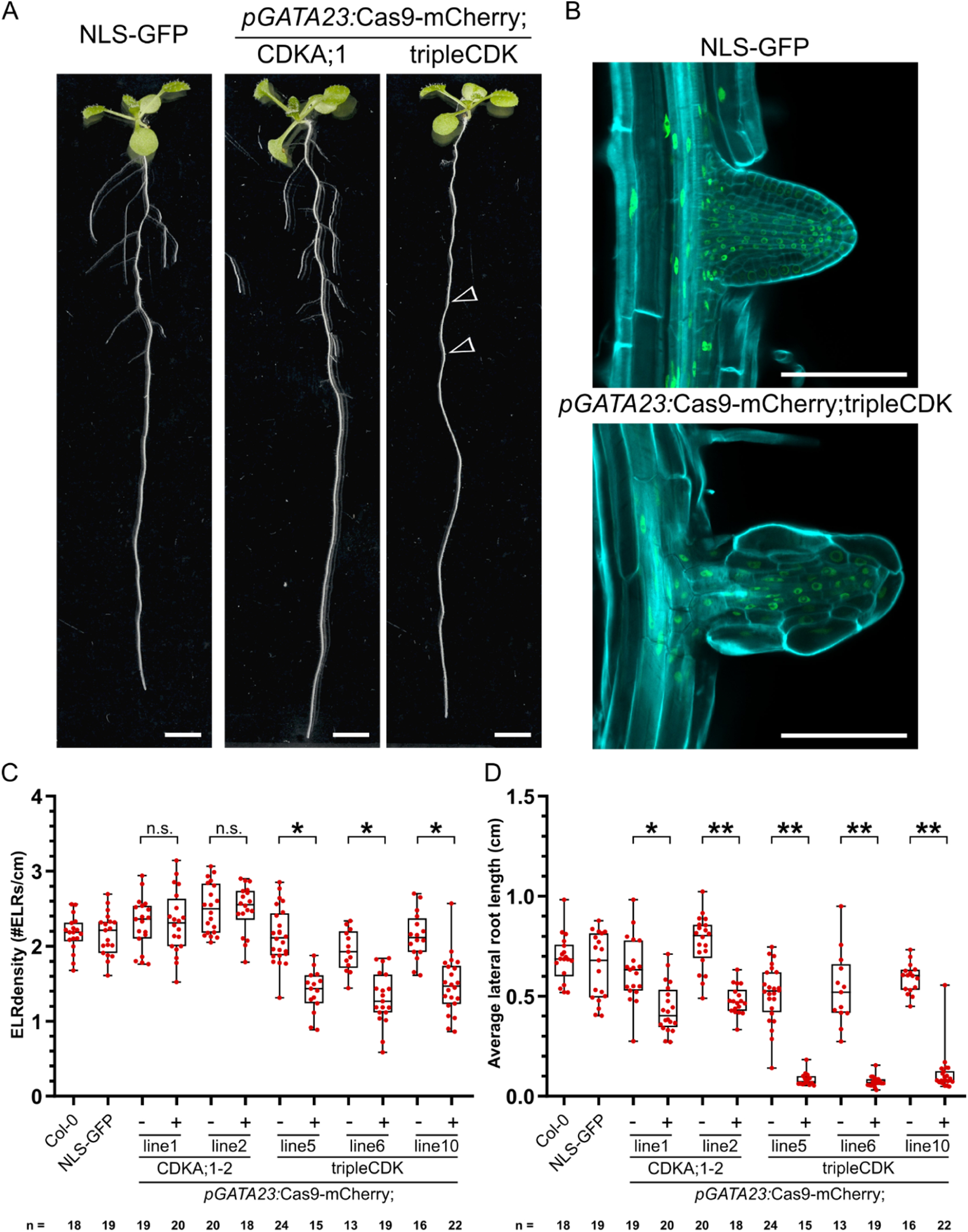
Lateral root-specific gene knockout of cell cycle regulators using *pGATA23*-CRISPR-TSKO. **A**, Representative 12 day old seedlings of NLS-GFP, T2 seedling of *pGATA23*:Cas9-mCherry;CDKA;1-2, and T2 seedling of *pGATA23*:Cas9-mCherry;CDKA;1-2,CDKB1-1 (tripleCDK). Arrowheads show emerged lateral roots with an extremely reduced cell number. Scale bars represent 0.5cm. **B**, Confocal images of an emerged lateral root in NLS-GFP and *pGATA23*:Cas9-mCherry;tripleCDK. GFP in green, and cell wall stained with calcofluor white displayed in cyan. Scale bars represent 100 µm. **C**, Quantification of the emerged lateral root (ELR) density of Col-0, NLS-GFP, and FAST negative (-) and positive (+) T2 seedlings of *pGATA23*-CRISPR-TSKO lines targeting either only *CDKA;1* or simultaneously *CDKA;1*, *CDKB1;1*, and *CDKB1;2.* ELR density was compared between FAST positive and negative seedlings within each line via Poisson regression analyses. n.s. indicates not significant with an α=0.05. * indicates *p* values smaller than 4×10^−4^. **D**, Quantification of average lateral root length of same seedlings as in C. A random effects model was used to estimate the effect of CRISPR-TSKO on the lateral root lengths between FAST positive and negative seedlings within each line. * indicates *p* values smaller than 6×10^−3^, ** indicates *p* values smaller than 1×10^−4^.

As the *CDK* gene family in Arabidopsis is composed of 10 partially redundant members (De Veylder et al., 2007), we hypothesized that the elimination of *CDKA;1* in lateral roots was being compensated for by the action of two B-type CDKs (*CDKB1;1*, AT3G54180 and *CDKB1;2*, AT2G38620). In contrast to the *cdka;1* single mutant, *cdka;1 cdkb1;1* double mutants are embryo lethal, with embryo development arresting after a few rounds of cell divisions (Nowack et al., 2012). Therefore, we combined the gRNA CDKA1-1 with one of two different gRNAs that simultaneously target both *CDKB1* genes to generate triple knockouts (*pPcUBI*:Cas9-mCherry;CDKA1-1,CDKB1-1 and *pPcUBI*:Cas9-mCherry;CDKA1-1,CDKB1-2). If effective, we expected severe seedling phenotypes or even failure to recover FAST-positive T1 seeds. Indeed, the few FAST-positive T1 seedlings we could recover showed severe developmental defects and many died in axenic culture (**Supplementary File 19**). We were able to isolate DNA from some of these seedlings and confirmed that indel frequencies were >90% for all three genes in four out of seven independent *pPcUBI*:Cas9-mCherry;CDKA1-1,CDKB1-1 lines (**Supplementary File 19 and 20**). The second *CDKB1* gRNA was less effective as most events had indel frequencies of only ∼20-70%. It is important to note that due to the severe growth defects in these mutants, we observed a negative selection pressure against events transformed with the *pPcUBI*:Cas9-mCherry;CDKA1-1,CDKB1-1 (**Supplementary File 21**) and many of the most severely affected plants did not yield sufficient DNA for genotyping. Nevertheless, these results indicate that both gRNAs are effective, with the CDKB1-1 gRNA being more efficient.

We next generated triple *CDK* lateral root knockouts via a *pGATA23* construct. Macroscopically, the transgenic lines exhibited an apparent lack of lateral roots (Figure 5A). However, upon closer inspection, we found that lateral roots did form, but they arrested growth soon after emergence (Figure 5B, D). These stunted lateral roots show the characteristic reduced number of cells and the presence of grossly enlarged epidermal and cortex cells in mutants severely affected in cell cycle progression (Nowack et al., 2012). Furthermore, we detected a slight but significant reduction in emerged lateral root density in FAST-positive segregants in three independent lines (Figure 5C), suggesting that some lateral roots did arrest before emergence.

## Discussion

### CRISPR-TSKO of essential genes enables their study in specific contexts

Targeted gene knockout experiments in plants typically have the objective of generating inheritable mutant alleles that will be transmitted to the offspring. The generation of such knockout lines is a powerful tool for the functional analysis of many genes of interest. However, this approach is difficult to apply to genes that are essential for cell survival, reproduction or those that have severe pleiotropic effects when mutated. Moreover, context-specificity of key regulators in developmental processes is often assumed by researchers, without experimental proof, while being aware of the non-context-specific expression. In this report we describe the design and validation of CRISPR-TSKO, a tissue-specific gene knockout approach in plants that can be used to overcome these limitations.

In total, we targeted nine genes using four different tissue-specific promoters. Several of the target genes (*PDS3, YDA*, *CDKA;1*) are essential for plant growth, development and/or reproduction. Mutations in *PDS3* induced by *pTMM*:Cas9-mCherry;GFP-1;PDS3 led to the expected defects in chlorophyll content and chloroplast formation (Qin et al., 2007) specifically in the stomatal lineage. Importantly, the active photosynthetic mesophyll tissue was not markedly affected in non Cas9-expressing cells which allowed these plants to develop similarly as the wild-type. This stands in contrast to the ubiquitous CRISPR knockout plants that were primarily dwarfed, albino and not viable in soil (Figure 3a**, Supplementary File 6A**).

In wild-type Arabidopsis, chlorophyll-containing chloroplasts are formed in epidermal pavement cells as well as stomatal guard cells, though they are much smaller than mesophyll chloroplasts (Barton et al., 2016). The function of chloroplasts in guard cells has been the subject of debate (Lawson, 2009). Recently, the discovery of the *gles1* (*green less stomata 1)* mutant further supports the hypothesis that functional chloroplasts in guard cells are important for stomatal responses to CO2 and light, resulting in stomatal opening (Negi et al., 2018). CRISPR-TSKO plants with mutated *PDS3*, or other genes required for chloroplast development and/or function, specifically in the stomatal lineage can be powerful tools to test these and other hypothesized functions of chloroplasts in guard cells.

The mitogen-activated protein kinase YDA has a plethora of roles during plant development including embryogenesis, epidermal patterning and root development (Musielak and Bayer, 2014; Smekalova et al., 2014). Accordingly, *yda* mutants have severe pleiotropic phenotypes. Already soon after fertilization, *yda* mutants fail to establish the first asymmetric division of the zygote and ensuing embryo development is severely compromised (Lukowitz et al., 2004). Some *yda* embryos do continue to develop into seedlings, but these rarely survive on soil and the few *yda* plants that flower are severely dwarfed and completely sterile (Bergmann et al., 2004; Lukowitz et al., 2004). While loss-of-function *yda* lines can be maintained in a heterozygous state, previous reports and our own experience show that only a small proportion of homozygous seedlings can be obtained either due to germination issues or very early seedling lethality (Lukowitz et al., 2004). This low recovery rate poses a considerable obstacle when designing and conducting experiments.

Using CRISPR-TSKO to target *YDA* only in the stomatal lineage, all transgenic events expressed a range of clustered-stomata phenotypes while other aspects of plant development were not notably compromised. Critically, all lines transferred to soil were fertile and we were able to generate normally-segregating T2 populations. These results demonstrate that by using CRISPR-TSKO we are able to uncouple the pleiotropic defects caused by *YDA* mutations to study its functions in the stomatal lineage.

The specific cellular defects caused by mutations in essential genes such as central cell cycle regulators are challenging to investigate due to lethality in the gametophyte or embryonic stage, and accordingly, low transmission rates (Nowack et al., 2012). CRISPR-TSKO enabled us to generate presumably higher-order *CDK* mutant lateral roots with striking cell proliferation defects on otherwise wild-type plants. These mutant plant lines offer a convenient opportunity to investigate the cellular defects caused by depletion of CDK proteins in an easily accessible tissue in the T1 or later transgenic generations. Interestingly, the cell proliferation in the stele of triple-CDK lateral roots appeared to be less affected than in the epidermis and cortex. Whether this is caused by differential turnover of CDK mRNA and/or proteins in different cell types, or by differential requirement of CDK activity in different tissues remains to be tested. Depletion of the CDKA;1 target RETINOBLASTOMA-RELATED1 (RBR1) has been shown to attenuate the cell proliferation defect in *cdka;1* mutants (Nowack et al., 2012), suggesting that tissue-specific differences of gene expression levels might contribute to a differential response to loss of CDK function. Alternatively, other CDK classes might be able to partly compensate for CDKA and CDKB loss of function in specific cell types (Inze and De Veylder, 2006). These different scenarios could be addressed by CRISPR-TSKO in the future. Similarly, many other central cell cycle or cell division regulators, of which no homozygous plants can be recovered, become amenable for detailed cellular investigation by CRISPR-TSKO.

### Generation of entire mutant organs with CRISPR-TSKO

To generate entire mutant organs on otherwise wild-type plants we targeted primordial founder cells responsible for the generation of the root cap and whole lateral roots. The root cap is an organ that covers and protects the stem cells at the root tip. Although it has relatively low tissue complexity, it encompasses many aspects of plant development – generation by stem cells, proliferation, differentiation, and finally programmed cell death– in a compact spatial and temporal frame of a few hundred micrometers and a couple of days (Kumpf and Nowack, 2015). SMB is a key transcription factor required for root cap maturation and programmed cell death and its expression starts immediately after the formative stem cell division in root cap daughter cells (Willemsen et al., 2008; Fendrych et al., 2014). We show that even an early acting gene such as *SMB* itself can be efficiently targeted thereby affecting root cap development. In this model system, the *pSMB*-CRISPR-TSKO vector toolkit could be particularly useful to study genes essential for basic cellular functions in this easily-accessible and nonessential root organ.

Lateral roots arise from a subset of stem cells situated in the pericycle at the xylem poles. These cells express *GATA23*, a gene that marks the onset of lateral root organogenesis and undergo tightly coordinated asymmetric cell divisions to generate cell diversity and tissue patterns, resulting in the development of a lateral root primordium (De Rybel et al., 2010). By targeting *GATA23*-expressing pericycle cells, we were able to generate plants with entirely mutated lateral roots. The generation of completely GFP-negative lateral roots in 87% of T1 events demonstrates the high efficiency of CRISPR-TSKO under *pGATA23*. Having a promoter at hand that is activated in the precursor cells of a new organ thus represents an effective means to generate whole organs devoid of the function of a gene of interest. Thus the use of CRISPR-TSKO may be an attractive alternative to grafting in certain experimental systems. Moreover, essential genes for primary root development such as *MONOPTEROS* hinder loss-of-function analysis during lateral root development (De Smet et al., 2010). The *pGATA23*-CRISPR-TSKO will enable us to uncouple the function of such genes involved in primary and lateral root development.

Auxin signaling is essential for lateral root initiation and development. The auxin response factors ARF7 and ARF19 are required for the auxin-induced pericycle cell divisions that constitute a lateral root initiation event. These divisions are strongly inhibited in *arf7arf19* double knockout mutants, which hardly produce any lateral roots (Okushima et al., 2007; Lavenus et al., 2013). *ARF7* is expressed in the initials of the vasculature and pericycle cells starting from the elongation zone, while *ARF19* is much more broadly expressed in the primary root (Okushima *et al*., 2005; Rademacher *et al*., 2011). Given their expression beyond the cells that actually contribute to lateral root formation, it has so far remained unresolved whether or not the role of ARF7 and ARF19 in lateral root initiation depends strictly on their activity in xylem-pole pericycle cells. Interestingly, targeted mutagenesis of *ARF7* and *ARF19* using the *pGATA23* does not result in a strong inhibition of lateral root initiation as observed in *arf7arf19* seedlings (Figure 4C).

This suggests that the function of ARF7 and ARF19 in lateral root precursor cells is not essential for lateral root development, and that their activity is required before the initiation of lateral root organogenesis and thus prior to the activation of *pGATA23*, or even in another tissue. This raises the question of when and in which cells of the primary root these ARFs are necessary for lateral root development. Alternatively, the ARF7/ARF19 mRNA and/or protein may persist in *GATA23*-expressing cells long enough to promote lateral root initiation. To test these hypotheses, we will be able to utilize CRISPR-TSKO with different promoters with unique spatio-temporal expression patterns. Furthermore, the use of fluorescently-tagged translational fusion ARF7/ARF19 lines will allow us to track their depletion upon CRISPR-TSKO targeting and establish the precise developmental window of ARF7/ARF19 signaling necessary for lateral root initiation.

### CRISPR-TSKO; A powerful and versatile tool

We developed a modular vector-cloning scheme based on the GreenGate system (Lampropoulos et al., 2013) to facilitate the construction of CRISPR-TSKO reagents. The modularity of the cloning system allows for the rapid assembly of new promoter sequences with Cas9 or any nuclease of choice. To further enable the use of this technology, we have developed a Cas9-P2A-mCherry;GFP-1 destination vector (pFASTR-BsaI-*Cm*R-*ccdB*-BsaI-Cas9-P2A-mCherry-G7T-AtU6-GFP-1) with an empty promoter module containing a *ccd*B/*Cm*R cassette flanked by BsaI restriction sites. Any promoter in a GreenGate entry vector can be inserted into this destination vector with a single Golden Gate reaction and researchers can immediately test the suitability of their promoter for CRISPR-TSKO in a GFP-expressing background line. Targeting specifically in spatial and temporal contexts can also be readily achieved with the inducible and tissue-specific plant expression systems that utilize GreenGate technology (Schurholz et al., 2018).

The cloning reagents used here are inexpensive, require minimal hands-on time and can be readily adopted by any laboratory. The system can currently accommodate up to twelve gRNAs by the use of an AarI linker and additions of six paired-gRNA entry modules (**Supplementary File 2 and Supplementary Methods**). By recycling the AarI-linker it is possible to clone even more gRNAs. For ease of use, a workflow was developed to substitute the AarI restriction sites in the linker for BsaI restriction sites flanking the *ccd*B and *Cm*R selectable markers (Figure 1**, Supplementary File 2, and Supplementary Methods**). This strategy avoids the need for separate AarI digestions of regularly-used destination vectors and provides a negative selection marker for the original destination vector in *ccd*B-sensitive *Escherichia coli* cells (*e.g.* DH5α). Alternatively, additional expression cassettes can be sequentially inserted into the AarI-*SacB* or BsaI-*ccdB* linkers. Hence, the system presented here can be easily used and modified for a variety of genome engineering applications such as transcriptional regulation (Lowder et al., 2015) and base editing (Marzec and Hensel, 2018).

### Considerations for use of CRISPR-TSKO

One general characteristic of CRISPR-TSKO is the continuous *de-novo* generation of mutations in cells that start to express Cas9. In the case of the root cap, every newly-generated root cap cell starting to express SMB can create a novel gene knockout event. In the stomatal lineage, every cell that starts expressing TMM or FAMA generates independent lineages, and in the case of the lateral root, every *GATA23*-expressing founder cell will contribute individual mutations to the lateral root primordium (von Wangenheim et al., 2016). Therefore, unlike in ubiquitous, inheritable mutant approaches, no defined mutant alleles are generated. Most mutations are small (1-3 bp) indels causing frame shifts and early stop codons, but, depending on the gRNA, some will also lead to in-frame missense mutations. Despite this source of variation, we are able to observe knockout phenotypes of varying degrees for all of the genes investigated. Furthermore, DNA repair outcomes from Cas9-mediated cleavage are non-random (Allen et al., 2019). Indeed, mutation analysis of the CRISPR-TSKO events show that gRNAs largely give the same indel spectrum regardless of the tissue-specific promoter used. For example, the most common indel generated for the GFP-1 target locus in *SMB*-, *TMM*-, *FAMA*-, and *GATA23*-expressing cells is a 1-bp insertion followed by 1- and 3-bp deletions (**Supplementary Files 5, 9, 10, and 14**).

Some gRNAs do not induce mutations with a high efficiency. The ARF7-1 and ARF19-1 gRNAs are a clear example of this, with ARF7-1 giving indel frequencies of 20-60% and ARF19-1 giving only 3-13%. These low indel frequencies are the most likely reason for the lack of a lateral root phenotype in the ubiquitously-expressed Cas9 T1 plants. Based on this, and that indel outcomes are non-random, we recommend that users of CRISPR-TSKO initially generate multiple gRNAs to target genes of interest as we have done in most of the experiments reported here. In cases where the targeted cells will be of low abundance (only a few cells targeted, e.g. *pSMB*, or knockout of essential genes, e.g. *CDKA;1*) and it would therefore be practically challenging to obtain sufficient material for genotyping, controlling Cas9 expression with a ubiquitous promoter such as *PcUbi* or *GATA23* (to make mutant lateral roots) is a reasonable way to test the efficiency of a gRNA. This experiment can also establish whether or not a gene is essential when an efficient gRNA is identified.

Users should also consider targeting functional domains as is generally recommended with any standard knockout strategy. For example, the gRNA GFP-1 used here targets the essential Gly67 residue for GFP fluorescence, so that even in-frame mutations result in a loss of fluorescence (Fu et al., 2015). Hence, gRNAs targeting genes of interest in particularly sensitive sites, such as crucial interaction domains or active sites, can further increase the likelihood of CRISPR-TSKO being effective.

We observed a strong correlation between gene knockout and Cas9-mCherry expression, which can be used to facilitate event selection. Furthermore, targeting of *GFP* alongside a gene of interest in a ubiquitously-expressing NLS-GFP background revealed that knockout of GFP strongly correlated with mutagenesis of the endogenous genes *SMB* and *PDS3*. Thus, both tagged Cas9 as well as knockout of reporter genes facilitate the selection of successful knockout events in tissues and organs. Moreover, the efficiency of new promoter sequences to drive expression of Cas9 can be evaluated by using both the gRNA GFP-1 and the NLS-GFP background plant line as used here. While loss of GFP signal should not be taken as definitive proof that the function of a gene of interest is also lost, it allows for an easy readout when testing CRISPR-TSKO in new cell types and developmental contexts.

### Limitations of CRISPR-TSKO

Depending on the promoter used or gene targeted, CRISPR-TSKO experiments might not always be straightforward. This is illustrated with our use of the *pFAMA* promoter sequence. While we initially were unable to observe an obvious microscopic phenotype when targeting *GFP* and *PDS3*, we did observe a reduced GFP signal by flow cytometry for one transgenic line and DNA mutations were detected in all four sorted lines. Therefore, Cas9 expression in these cells led to DNA mutations (albeit, at a lower frequency than other experiments), but conferred only a modest phenotypic effect. If the generation of indels are independent events and given the indel frequencies we observed (23-75%; **Supplementary File 10**), we would expect ∼5-56% of the guard cells to be knocked out for GFP. However, guard cells completely lacking GFP expression were not observed in our *pFAMA* experiments. In our experiments with the *pSMB*, residual GFP fluorescence is detectable in the two youngest root cap layers, with some overlap between mCherry and GFP signals (Figure 2A,B). Therefore, we hypothesize that mRNA and/or protein turnover is required before knockout phenotypes can be observed. The speed of these processes likely depends of the stability of the particular mRNA and protein, and, considering our *pFAMA* observations, might also depend on the cell type in question.

The negative results presented here highlight that these dynamics should be considered on a gene-by-gene and tissue-specific promoter basis when designing and analyzing CRISPR-TSKO experiments. Therefore, ideally more than one promoter should be evaluated when targeting novel cell types with CRISPR-TSKO before more labor-intensive phenotyping is performed for genes of interest. This test can simply be performed by analyzing the ubiquitously expressing NLS-GFP line transformed with the highly efficient gRNA GFP-1 under the control of a new promoter of choice. Alternatively, translational fluorescent fusion lines can be used to monitor the elimination of a protein of interest from a particular cell type.

In conclusion, cell type-, tissue-, or organ-specific gene knockout by targeted expression of Cas9 is a powerful means of functional genetic analysis in specific spatial and temporal contexts of plant development. This is especially true for genes that are widely expressed or have fundamental roles for cell survival or plant reproduction. CRISPR-TSKO allows for the rapid generation of stable transgenic lines with *de novo* somatic DNA mutations specifically to the cell, tissue or organ of interest. Due to its flexibility and ease of use, we foresee this tool as enabling the discovery of context-specific gene functions. Moreover, the scalability of the system allows for quick initial investigation of candidate genes with the reduced influence of pleiotropic effects. As with other CRISPR applications, CRISPR-TSKO is forward-compatible to incorporate upcoming future variations of CRISPR gene modification. Together with the virtually unlimited possibilities to combine different promoters, reporters, or tags in CRISPR-TSKO, this technology presents a powerful addition to the molecular genetics tool-box for plant biology research.

### Distribution of modules, plasmids, and protocols

All cloning modules and plasmids reported here are available via the VIB-UGent Center for Plant Systems Biology Gateway Vector website (**Supplementary File 3**; https://gateway.psb.ugent.be/search) or via Addgene (https://www.addgene.org/). See **Supplementary methods** for detailed cloning protocols.

Seeds for the *pHTR5:NLS-GUS-GFP* line are available upon request.

## Methods

### Cloning

All cloning reactions were transformed via heat-shock transformation into *ccdB*-sensitive DH5α *E. coli* or One Shot™ *ccd*B Survival™ 2 T1R Competent Cells (ThermoFisher Scientific). Depending on the selectable marker, the cells were plated on LB medium containing 100 µg/mL carbenicillin, 100 µg/mL spectinomycin, 25 µg/mL kanamycin or 10 µg/mL gentamycin. Colonies were verified via colony-touch PCR, restriction digest and/or Sanger sequencing by Eurofins Scientific using the Mix2Seq or TubeSeq services. All cloning PCR reactions were performed with either Q5^®^ High-Fidelity DNA Polymerase (New England Biolabs) or iProof^TM^ High-Fidelity DNA Polymerase (BioRad Laboratories). Gibson assembly reactions were performed using 2x NEBuilder Hifi DNA Assembly Mix (New England Biolabs). Column and gel purifications were performed with Zymo-Spin^TM^ II columns (Zymo Research).

#### Golden Gate entry modules

Golden Gate entry modules were made by PCR amplification of gene fragments and inserting the purified PCR product into BsaI-digested GreenGate entry vector (Lampropoulos et al., 2013) via restriction-ligation using BsaI (New England Biolabs) or Gibson assembly. See **Supplementary File 22** for all primers used. All generated clones were verified via Sanger sequencing.

The coding sequence for mTagBFP2, based on a previously reported mTagBFP2 (Pasin et al., 2014), was PCR amplified with primers RB42 and RB43 from a synthesized fragment (Gen9) and inserted into a BsaI-digested pGGC000 plasmid via ligation.

The unarmed gRNA modules were cloned by amplifying the AtU6-26 promoter and gRNA scaffold from previously-described Golden Gate entry vectors (Houbaert et al., 2018). The amplification was done using the forward primer 120 and the reverse primers 283, 284, 230, 231, 232 and 233 for the B to G overhangs, respectively. This removed an unwanted attB2 site. PCR products were digested with BsaI and ligated into the respective empty entry vectors. The unarmed gRNA modules were further adapted by adding the *ccdB* negative selectable marker. The *ccdB* gene was PCR amplified from pEN-L1-AG-L2 with oligos 391 and 392 and inserted into BbsI-digested unarmed gRNA modules via Gibson assembly. pGG-F-AtU6-26-AarI-AarI-G was made by annealing oligos 345 and 346, and ligating these into the BbsI-digested vector pGG-F-AtU6-26-BbsI-Bbs-G.

The linker modules for Golden Gate were constructed as previously described (Houbaert et al., 2018). The entry module pGG-F-A-AarI-AarI-G-G was made by annealing oligos 361 and 362, and ligating into the BsaI-digested entry vector pGGF000. The variable linker pGG-F-A-AarI-SacB-AarI-G-G, based on the SacB sequence from pMA7-SacB (Lennen et al., 2016), was synthesized on the BioXP3200 DNA synthesis platform (SGI-DNA) and inserted into a BsaI-digested pGGF000 plasmid via Gibson assembly.

The variable linker modules were made by PCR-amplifying the AarI-SacB fragment from pGG-F-A-AarI-SacB-AarI-G-G with the respective primers 1589-1600 (Supplementary Table 3). PCR products were gel purified and inserted via Gibson assembly into pGG-A-m43GW-B (*unpublished*) pre-digested with ApaI and SacI. Clones were verified with oligo 1658.

#### GATEWAY^TM^ destination vectors

pGG-A-pOLE1-B, pGG-B-OLE1-C, pGG-D-linker-E (Lampropoulos et al., 2013), pGG-E-NOST-F and pGG-F-LinkerII-G were assembled with either pGG-C-mRuby3-D or pGG-C-GFP-D (Lampropoulos et al., 2013) into pEN-L1-A-G-L2. The ligation reactions were used as templates for PCR with the primers 195 and 196. The PCR products were cloned via Gibson assembly into pGGK7m24GW (Karimi et al., 2005) linearized with KpnI and XbaI. Clones were verified by Sanger sequencing. The resulting vectors containing the red and green fluorescent FAST markers were named pFASTRK24GW and pFASTGK24GW, respectively.

#### Proof-of-concept vectors

The Golden Gate entry modules pGG-A-*pSMB*-B, pGG-B-Linker-C, pGG-C-Cas9PTA*-D, pGG-D-Linker-E, pGG-E-G7T-F and pGG-F-linkerII-G were assembled in pEN-L4-AG-R1 (Houbaert et al., 2018), resulting in the vector pEN-L4-*pSMB*-Cas9PTA-G7T-R1. The Golden Gate entry module pGG-A-AtU6-26-BbsI-BbsI-B and pGG-B-linkerII-G were assembled in pEN-L1-A-G-L2 (Houbaert et al., 2018), resulting in the vector pEN-L1-AtU6-26-BbsI-BbsI-L2. The BbsI restriction sites were swapped with a fragment containing the *ccdB* and *CmR* selectable markers flanked with BsaI sites. This fragment was PCR amplified from the plasmid pEN-L4-A-G-R1, using primers 1436 and 1437. The fragment was BsaI-digested and ligated with T4 DNA Ligase in the BbsI-digested vector pEN-L1-AtU6-26-BbsI-BbsI-L2, resulting in the vector pEN-L1-AtU6-26-BsaI-BsaI-L2. pEN-L4-*pSMB*-Cas9PTA-G7T-R1 and pEN-L1-AtU6-26-BsaI-BsaI-L2 were recombined in pGGB7m24GW (Karimi et al., 2005) via a MultiSite Gateway reaction according to manufacturer’s recommendations. This vector was called pB-*pSMB*-Cas9-G7T-AtU6-BsaI-BsaI-gRNA scaffold.

Oligos 138 and 139 (GFP-1 target) and oligos 134 and 135 (GFP-2 target) were annealed by adding 1 µL of each 100 µM oligonucleotide in 48 µL of MQ and incubating with a slow cooling program on the thermal cycler (5 minutes at 95°C; 95-85°C, −2°C/second; 85-25°C, −0.1°C/second). These annealed oligonucleotides were cloned via a Golden Gate reaction into pB-*pSMB*-Cas9-G7T-AtU6-BsaI-BsaI-gRNA scaffold. The Golden Gate reaction conditions are described in **Supplementary Methods**. The resulting vectors were named pB-*pSMB*-Cas9-G7T-AtU6-GFP-1 and GFP-2.

#### Golden Gate destination

The Golden Gate destination vectors were cloned by amplifying the *CmR* and *ccdB* selection cassettes, flanked by the Golden Gate cloning sites A-G, from pEN-L1-AG-L2 using primers 298 & 313. PCR products were column purified and cloned via Gibson assembly in the HindIII and PstI linearized Gateway destination vectors pGGP7m24GW, pGGK7m24GW, pGGB7m24GW, pGGPH7m24GW (Karimi et al., 2005), pFASTRK24GW and pFASTGK24GW. The resulting vectors were named respectively pGGP A-G, pGGK A-G, pGGB A-G, pGGH A-G, pFASTRK A-G, pFASTGK A-G. All clones were verified by Sanger sequencing and diagnostic digest with NotI.

To generate the pFASTR-A-G destination vector, the Golden Gate entry modules pGG-A-pOLE1-B, pGG-B-OLE1-C, pGG-C-mRuby3-D, pGG-D-Linker-E, pGG-E-NOST-F, pGG-F-linkerII-G were assembled in pGGP-A-G. Subsequently the *CmR* and *ccdB* selection cassettes, flanked by the Golden Gate cloning sites, were PCR amplified using primers 298 and 430 from pEN-L1-AG-L2 and inserted via Gibson assembly.

#### One-step CRISPR-TSKO cloning vectors

For cloning two gRNAs in a destination vector, we followed a similar approach as previously described (Xing et al., 2014), with some modifications. A plasmid was generated to serve as a PCR template for 2-gRNA vectors. The Golden Gate entry modules pGG-A-AtU6-26-BbsI-BbsI-B, pGG-B-Linker-C (Lampropoulos et al., 2013), pGG-B-AtU6PTA-C (Houbaert et al., 2018) and pGG-D-linkerII-G were assembled in pEN-L1-A-G-L2 to generate pEN-2xAtU6 template. The clone was verified by Sanger sequencing.

The extended protocol for making one-step, CRISPR-TSKO cloning vectors can be found in **Supplementary Methods**. In summary, six different entry modules are combined via Golden Gate assembly in a destination vector. The A-B entry module contains the tissue-specific promotor, the C-D module contains the Cas9 endonuclease and can be combined with an N-terminal tag (B-C) or C-terminal tag (D-E), and the E-F entry module contains the terminator. For making a vector that is compatible with cloning one or two gRNAs, the F-G module pGG-F-AtU6-26-AarI-AarI-G is used (Figure 1). Upon digestion with AarI, this vector can be loaded directly with one or two gRNAs. Alternatively, the AarI sites can be replaced for a fragment containing BsaI sites flanking the *ccdB* and *CmR* selectable markers. Two gRNAs can be cloned via a PCR reaction on the pEN-2xAtU6 template using primers that contain gRNA spacer sequences via a Golden Gate reaction. More details can be found in **Supplementary Methods**.

For making a vector that is compatible with multiple gRNA’s (up to 12) the Golden Gate cloning is slightly modified. The initial Golden Gate reaction is performed with an F-G linker containing AarI restriction sites (**Supplementary File 2**). Upon AarI digestion, this vector can be directly loaded with six Golden Gate entry modules containing one or two AtU6-26 promotors and gRNAs. Alternatively, a similar strategy to replace the AarI sites by the *ccdB* and *CmR* selectable markers flanked with BsaI sites can be followed. All gRNA target sequences are in **Supplementary File 23**.

#### Expression vector with an empty promoter module

The armed gRNA module pGG-F-AtU6-26-GFP-1-G was made by annealing oligos 138 and 139 (GFP-1 target), and ligating these via a Golden Gate reaction in pGG-F-AtU6-26-BbsI-ccdB-BbsI-G. The entry modules pGG-A-AarI-SacB-AarI-B, pGG-B-Linker-C, pGG-C-Cas9PTA*-D, pGG-D-P2A-mCherry-NLS-E, pGG-E-G7T-F and pGG-F-AtU6-26-GFP-1-G were cloned into pFASTR A-G via a Golden Gate reaction. This vector was digested with AarI and the upper band was gel purified. The PCR product of the reaction on pEN-L4-AG-R1 using oligos 1879 and 1880, was cloned into the AarI-digested fragment using Gibson assembly. The resulting vector, pFASTR-BsaI-CmR-ccdB-BsaI-Cas9-P2A-mCherry-G7T-AtU6-GFP-1, was verified by restriction digest using PvuII and sequencing. Entry modules containing a promoter can easily be cloned in this vector via a Golden Gate reaction.

### Plant lines used in this study

The *smb-3* line is derived from the SALK collection (SALK_143526C). The NLS-GFP line (pHTR5:NLS-GUS-GFP) was previously reported (Ingouff et al., 2017). The *arf7 arf19* double mutant (Okushima et al., 2005) is derived from the ABRC collection (CS24629). The *yda-1* mutant was previously reported (Lukowitz et al., 2004).

### Plant transformation

Plant vectors were transformed in *Agrobacterium tumefaciens* C58C1 by electroporation. Transformation in pHTR5:NLS-GUS-GFP was performed via the floral dip method (Clough and Bent, 1998). For constructs containing the bar selectable marker, the T1 seed selection was done on 1/2 MS medium + 10 mgL^-1^ Glufosinate-ammonium (Sigma-Aldrich). For construct containing the FASTR screenable marker, the T1 transgenic seeds were selected under a Leica M165FC fluorescence stereomicroscope. Resistant seedlings or FASTR-positive seeds were transferred to Jiffy-7 pellets® and grown in a greenhouse at 21°C under a 16-hour day regime.

### DNA extraction and molecular analysis

Seedling, leaves or roots were frozen and disrupted to powder using a TissueLyser (Retsch MM300). DNA was extracted using a modified version of the protocol from Edwards et al. (Edwards et al., 1991). The modifications consisted of an adapted extraction buffer (100 mM Tris HCl pH 8.0, 500 mM NaCl, 50 mM EDTA, 0.7% SDS) and a 70% ethanol washing step before dissolving the pellet. A region around the CRISPR/Cas9 target site was PCR amplified using the ALLin™ Red Taq Mastermix, 2X (highQu GmbH) with the following program on the thermocycler: 95°C for 3 minutes, followed by 33 cycles (30 seconds at 95°C, 30 seconds at the annealing temperature, 1 minute/kb at 72°C), 72°C for 5 minutes. The PCR products were analyzed via agarose gel electrophoresis and the clean-up was either done by bead purification with HighPrep^TM^ PCR (MAGBIO) or column purification with the DNA Clean & Concentrator^TM^ kit (Zymo Research). The purified samples were send for Sanger sequencing (Eurofins Scientific) and analyzed using TIDE (version 2.0.1) (Brinkman et al., 2014).

### Confocal microscopy for original proof of concept

T1 seedlings were imaged on a Zeiss LSM710 confocal microscope. GFP was excited at 488 nm and acquired between 500-550 nm. T2 seedlings were imaged on a Leica SP8X confocal microscope. GFP was excited at 488 nm and acquired between 500-530 nm.

### Confocal microscopy

Seedlings were imaged on a Leica SP8X confocal microscope. For root imaging, GFP was excited at 488 nm and acquired between 500-530nm. mCherry was excited at 594 nm and acquired between 600-650nm. Samples were either stained with 20 µg/mL DAPI or with 10 µg/mL propidium iodide in 0.43 gL^-1^ Murashige and Skoog salts with 94 µM MES.H2O medium. DAPI was excited at 405 nm and acquired between 410-480 nm in sequential mode.

For stomata imaging, cotyledons were vacuum infiltrated with 20 µg/mL of DAPI in 0.43 gL^-1^ Murashige and Skoog salts with 94 µM MES.H2O medium. Samples were imaged in sequential mode. DAPI was excited at 405 nm and acquired between 410-450 nm. GFP was excited at 488 nm and acquired between 500-530 nm. mCherry was excited at 594 nm and acquired between 600-650 nm. Chlorophyll fluorescence was excited at 488 nm and acquired between 680-730 nm. Images were analyzed using Fiji (Schindelin et al., 2012).

To image lateral root primordia, seedlings were cleared using the ClearSee protocol (Kurihara et al., 2015; Ursache et al., 2018) in combination with cell wall staining using Calcofluor White M2R (Sigma) on a Leica SP8X confocal microscope. Calcofluor White was excited at 405 nm and acquired between 430-470 nm. GFP was excited at 488 nm and acquired between 500-525 nm. mCherry was excited at 594 nm and acquired between 600-630 nm.

### Epifluorescence microscopy

Cotyledons of FASTR positive seedlings were mounted on distilled water and observed on a Zeiss Observer.Z1 using a Plan-Apochromat 20x/0.8 DICII objective. GFP fluorescence was observed with a BP 470/40 filter for excitation, a FT 495 beam splitter, and a BP 525/50 emission filter. mCherry was observed with a BP 545/25 filter for excitation, a FT 570 beam splitter, and a BP 605/70 emission filter.

### Segmentation and analysis of root cap nuclei

Root tip image stacks were segmented and nuclei intensity measurements performed using the interactive learning and segmentation toolkit ilastik 1.3.0 (Sommer et al., 2011). Intensity of GFP and mCherry were measured for segmented nuclei with a probability equal or higher than 0.95 of belonging to root cap cells. Based on mCherry measurements in the NLS-GFP line (**Supplementary File 24**), a threshold of 25 was established as a minimum signal for mCherry.

### Protoplast preparation and cell sorting

Protoplasting was performed as previously described (Bargmann and Birnbaum, 2010). Briefly, for *pSMB*-CRISPR-TSKO lines, root tips of 5 day old seedlings grown under continuous light on 0.43 gL^-1^ Murashige and Skoog salts with 94 µM MES.H2O medium were incubated in protoplasting solution consisting of 1.25% cellulase (Yakult, Japan), 0.3% Macerozyme (Yakult, Japan), 0.4 M mannitol, 20 mM MES, 20 mM KCl, 0.1% BSA and 10 mM CaCl2 at pH 5.7 for 3 hours. Samples were then filtered through a 40 µm filter and the flow through centrifuged at 150xg for 10min. Supernatant was discarded and protoplasts were recovered in ice-cold resuspension buffer. Resuspension buffer was of same constitution as protoplasting buffer with the omission of both cellulase and macerozyme. For lines targeting stomatal lineages, cotyledons of 5-day old seedlings were processed as above but with a 12 hours incubation time to get proper release of guard cells.

Root tip protoplasts were sorted into 1.5 ml Eppendorf tubes containing 500 µl of resuspension buffer using a BD FACSAriaII, equipped with 3 lasers (405 nm, 488 nm and 633 nm). To account for the double presence of GFP and mCherry in some samples, the cotyledon protoplasts were sorted into 1.5 ml Eppendorf tubes containing 500 µl of resuspension buffer using a BD FACSMelody, equipped with 3 lasers (405 nm, 488 nm and 561 nm). The 561 nm laser in the BD FACSMelody made a better separation possible due to a better excitation of the mCherry. All FACS sorting reports can be found in **Supplementary Files 8 and 11**.

### Quantification of lateral root density

Seeds were sown on half-strength Murashige and Skoog (MS) medium (Duchefa Biochemie B.V.), supplemented with 1% (w/v) sucrose and 0,8% (w/v) agar, at pH 5,7 and stratified for 2 days in the dark at 4°C. Seedlings were grown vertically for 12 days in continuous light (100 μmol m^-2^s^-1^) at 22°C. Presence/absence of Cas9-mCherry signal was scored using a Leica M165FC fluorescence stereomicroscope. The number of emerged lateral roots was determined for every seedling using a stereo microscope and root lengths were measured via Fiji (ImageJ 1.52n) (Schindelin et al., 2012) using digital images obtained by scanning the petri dishes.

### Stomata analysis of cotyledons in *YDA* targeting lines

The cotyledon epidermis of seedlings 10 days post germination was visualized by clearing cotyledons in 100% ethanol and incubation at 60 degrees in 90% ethanol / 10% acetic acid for 30 minutes and ethanol / 1,25 M sodium hydroxide (1:1 v/v) for 2 hours. Next, cotyledons were incubated overnight at room temperature in lactic acid saturated with chloral hydrate, washed in 100% lactic acid and mounted for differential interference contrast microscopy (Olympus BX51). Images (430 µm x 566 µm) from the midline to the margin on abaxial surfaces were generated. Thirty five to 40 cotyledons of individual seedlings were evaluated per genotype.

### Statistical analysis

For segmentation and analysis of root cap nuclei, Spearman’s Correlation Coefficient between median root cap signal of GFP and mCherry was calculated using SAS (Version 9.4, SAS Institute Inc., 2013 Cary, North Carolina). For the comparison of emerged lateral root densities, the number of emerged lateral roots was modelled by Poisson regression using the primary root length as an offset variable and genotype as fixed effect. In the presence of overdispersion, the negative binomial distribution was used instead of the Poisson distribution. The analysis was performed with the genmod procedure from SAS (SAS/STAT analytical product 14.3, SAS Institute Inc., 2017, Cary, North Carolina). Post-hoc comparison tests were done using the capabilities of the plm procedure. In case of multiple testing, P-values were adjusted using the Dunnett’s method. For the comparison of lateral root lengths, a random effects model was used to estimate the effect within each line. The root length was log transformed to stabilize the variance. Numerator degrees of freedom for the type III test of effect were calculated according to Kenward-Rogers as implemented in the mixed procedure from SAS (Version 9.4, SAS Institute Inc., 2013 Cary, North Carolina). The assumptions were checked by residual diagnostics. The SAS code is available upon request.

## Acknowledgements

We thank Dominique Bergmann, Camila Lopez-Anido, Michael Raissig, and the members of the Programmed Cell Death and Plant Genome Editing teams at VIB for constructive discussions. We also thank Veronique Storme for assistance with statistical analysis. The GreenGate plasmid kit used for generation of plant transformation constructs was a gift from Jan Lohmann (Addgene kit # 1000000036). The coding sequence for mRuby3 was derived from pNCS-mRuby3, which was a gift from Michael Lin (Addgene plasmid # 74234). We thank Carina Braeckman (VIB-UGent Center for Plant Systems Biology) for the help with the A. thaliana floral dip transformation. We thank Nico Smet, Miguel Riobello y Barea, Thomas Farla, and Sandra Lefftz for greenhouse support. We thank Debbie Rombaut and Freya De Winter for the help with the preparation of root protoplasts. We thank Eugenia Russinova for a critical review of the manuscript. We gratefully acknowledge funding by the ERC StG PROCELLDEATH (Project 639234) to M.K.N., and by the FWO 1174119N to M.L.P.

## Author contributions

W.D., R.A.B., M.K.N., and T.B.J. conceived and devised the study. M.K. adapted the GreenGate vectors and provided additional cloning support. W.D. constructed the vectors and performed the genotyping analysis. R.A.B., M.L.P., N.V. and J.J. performed the experiments, imaging and analysis. W.D., J.J. and R.A.B. performed statistical analysis.

G.V.I. and R.A.B. performed the FACS experiments and analysis. W.D., T.B.J., R.A.B.,

M.K.N. and T.B. wrote the manuscript with contributions from all other authors.

## Additional information

**Supplementary information** accompanies this paper at XXX

