## Supplementary Files for "CRISPR-TSKO facilitates efficient cell type-, tissue-, or organ-specific mutagenesis in Arabidopsis"

**Supplementary File 1**

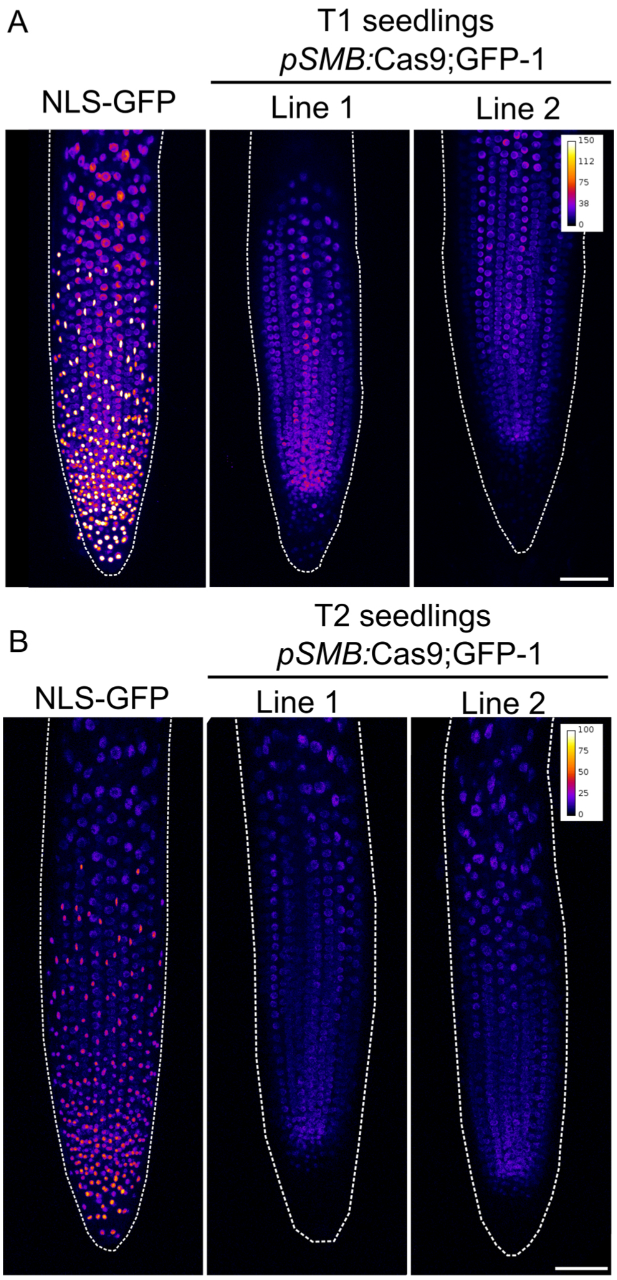

**Supplementary File 1 | Root cap-specific knockout of GFP**

**A**, Two independent T1 seedlings of *pSMB*:Cas9;GFP-1 displaying the root cap-specific knockout of GFP. **B**, Representative T2 progeny of lines shown in **A** displaying the same expected phenotype of loss of GFP signal in root cap cells. White dashed lines indicate root tip outline. Scale bars represent 50 µm.

#### Supplementary File 2

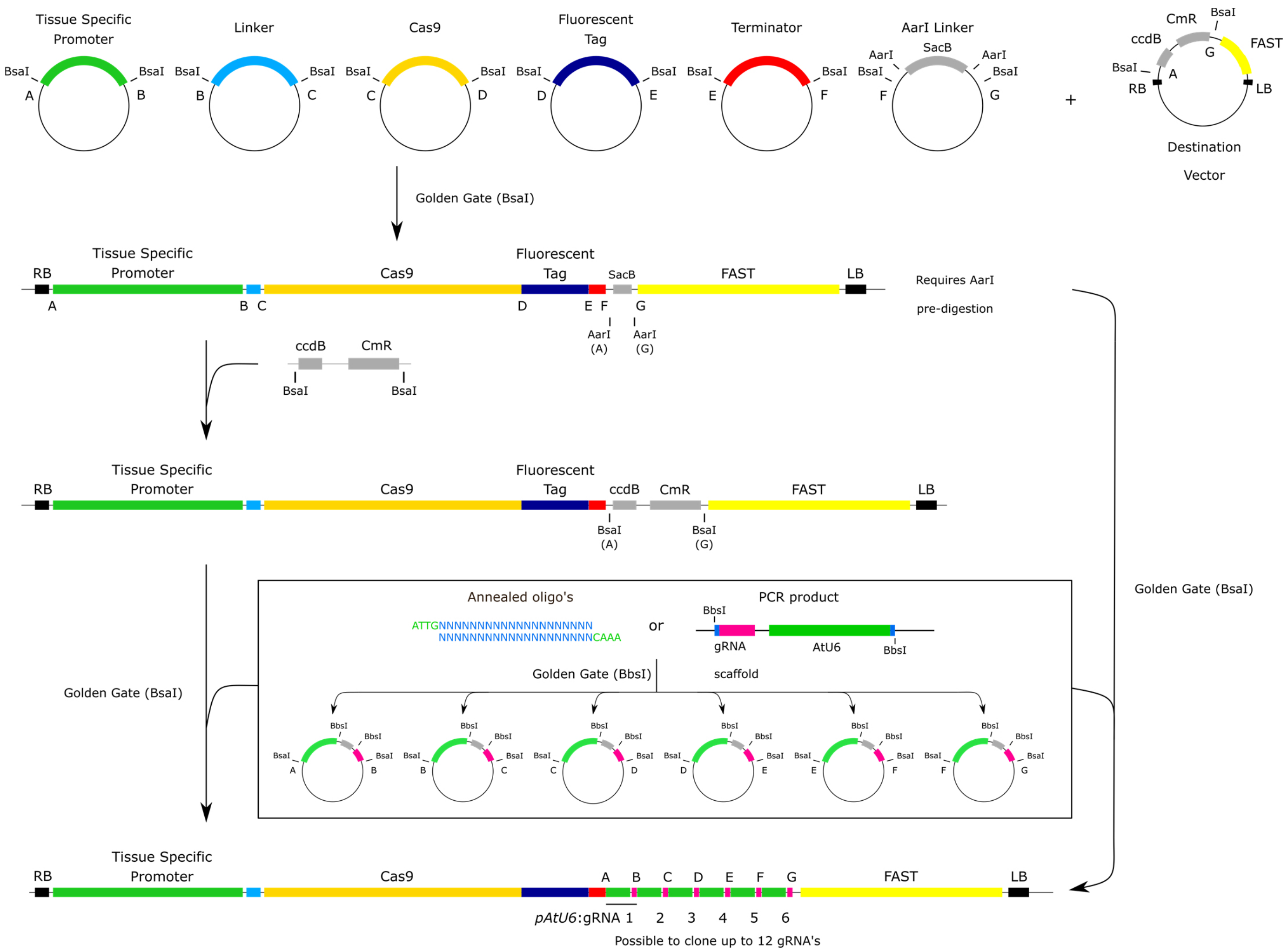

**Supplementary File 2 | Overview of the cloning scheme for making a vector with multiple gRNAs**

Six entry modules are combined in a destination vector via a Golden Gate reaction. Via the Golden Gate entry F-G module a linker with AarI restriction sites enters the destination vector. Upon digestion with AarI, Golden Gate A-G overhangs are generated allowing for a second Golden Gate reaction. Thus, upon pre-digestion with AarI, this vector can be directly used as a destination vector. More conveniently however, the AarI restriction sites can be exchanged for BsaI restriction sites to avoid the pre-digestion. To clone the gRNAs into the destination vector, six different Golden Gate entry modules can be 'loaded' with one (annealed oligos) or two gRNA (PCR product) cassettes. By combining these entry modules an expression vector can be built that contains up to 12 gRNAs. Detailed protocol in **Supplemental Methods**.

**Supplementary File 3. Plasmid Overview**

| Full Name | Type | Cloning sites[a] | Bacterial Selection | Plant Selection | Screenable Marker | Reference | Plasmid Deposit |
|---|---|---|---|---|---|---|---|
| **Gateway destination vectors** (*ccdB* $^{+}$) | | | | | | | |
| pGGP7m24GW | | R2-R4 | Spectinomycin | | | Karimi et al. (2005) | |
| pGGK7m24GW | | R2-R4 | Spectinomycin | Kanamycin | | Karimi et al. (2005) | |
| pGGB7m24GW | | R2-R4 | Spectinomycin | Glufosinate-ammonium | | Karimi et al. (2005) | |
| pGGH7m24GW | | R2-R4 | Spectinomycin | Hygromycin | | Karimi et al. (2005) | |
| pFASTRK24GW | | R2-R4 | Spectinomycin | Kanamycin | FAST Red | This work | |
| pFASTGK24GW | | R2-R4 | Spectinomycin | Kanamycin | FAST Green | This work | |
| **Gateway empty entry vectors** (*ccdB* $^{+}$) | | | | | | | |
| pEN-L4-AG-R1 | | L4-R1 | Kanamycin | | | Houbaert et al. (2018) | |
| pEN-L1-AG-L2 | | L1-L2 | Kanamycin | | | Houbaert et al. (2018) | |
| **Gateway entry vectors** | | | | | | | |
| pEN-L4-pSMB-Cas9PTA-G7-R1 | | L4-R1 | Kanamycin | | | This work | |
| pEN-L1-AtU6-26-BsaI-L2 | | L1-L2 | Kanamycin | | | This work | |
| **Golden Gate destination vectors** (*ccdB* $^{+}$) | | | | | | | |
| pGGP A-G | | A-G | Spectinomycin | | | This work | |
| pGGK A-G | | A-G | Spectinomycin | Kanamycin | | This work | |
| pGGB A-G | | A-G | Spectinomycin | Glufosinate-ammonium | | This work | |
| pGGH A-G | | A-G | Spectinomycin | Hygromycin | | This work | |
| pFASTR A-G | | A-G | Spectinomycin | | FAST Red | This work | |
| pFASTRK A-G | | A-G | Spectinomycin | Kanamycin | FAST Red | This work | |
| pFASTGK A-G | | A-G | Spectinomycin | Kanamycin | FAST Green | This work | |
| **Empty entry vectors** (*ccdB* $^{+}$) | | | | | | | |
| pGGA000 | | A-B | Ampicillin | | | Lampropoulos et al. (2013) | Addgene ID 48856 |
| pGGB000 | | B-C | Ampicillin | | | Lampropoulos et al. (2013) | Addgene ID 48857 |
| pGGC000 | | C-D | Ampicillin | | | Lampropoulos et al. (2013) | Addgene ID 48858 |
| pGGD000 | | D-E | Ampicillin | | | Lampropoulos et al. (2013) | Addgene ID 48859 |
| pGGE000 | | E-F | Ampicillin | | | Lampropoulos et al. (2013) | Addgene ID 48860 |
| pGGF000 | | F-G | Ampicillin | | | Lampropoulos et al. (2013) | Addgene ID 48861 |
| **Plant promoters** | | | | | | | |
| pGG-A-pPcUBI-B | *PcUBI (Petroselinum crispum UBIQUITIN)* promoter | A-B | Ampicillin | | | Houbaert et al. (2018) | |
| pGG-A-pOLE1-B | *OLE1 (OLEOSIN 1)* promoter | A-B | Ampicillin | | | This work | |
| pGG-A-pSMB-B | *SMB (SOMBRERO)* promoter | A-B | Ampicillin | | | This work | |
| pGG-A-pFAMA-B | *FAMA* promoter | A-B | Ampicillin | | | This work | |
| pGG-A-pTMM-B | *TMM (TOO MANY MOUTHS)* promoter | A-B | Ampicillin | | | This work | |
| pGG-A-pGATA23-B | *GATA23 (GATA transcription factor 23)* promoter | A-B | Ampicillin | | | This work | |
| **N-tags** | | | | | | | |
| pGG-B-OLE1-C | *OLE1 (OLEOSIN 1)* | B-C | Ampicillin | | | This work | |
| **Coding sequences** | | | | | | | |
| pGG-C-GFP-D | | C-D | Ampicillin | | | Lampropoulos et al. (2013) | Addgene ID 48827 |
| pGG-C-mRuby3-D | | C-D | Ampicillin | | | This work | |
| pGG-C-Cas9PTA*-D | *Cas9-SV40* with stopcodon | C-D | Ampicillin | | | Houbaert et al. (2018) | |
| pGG-C-Cas9PTA-D | *Cas9-SV40* | C-D | Ampicillin | | | This work | |

**Supplementary File 3. Plasmid Overview (Continued)**

| Full Name | Type | Cloning sites[a] | Bacterial Selection | Plant Selection | Screenable Marker | Reference | Plasmid Deposit |
|---|---|---|---|---|---|---|---|
| **C-tags** | | | | | | | |
| pGG-D-P2A-mCherry-NLS-E | *P2A-mCherry-NLS* | D-E | Ampicillin | | | This work | |
| pGG-D-P2A-mTagBFP2-NLS-E | *P2A-mTagBFP2-NLS* | D-E | Ampicillin | | | This work | |
| pGG-D-P2A-GFP-NLS-E | *P2A-GFP-NLS* | D-E | Ampicillin | | | This work | |
| **Plant Terminators** | | | | | | | |
| pGG-E-G7T-F | *G7* terminator | E-F | Ampicillin | | | Houbaert et al. (2018) | |
| pGG-E-NOST-F | *NOS* terminator | E-F | Ampicillin | | | This work | |
| **Unarmed gRNA modules** | | | | | | | |
| pGG-A-AtU6-26-BbsI-BbsI-B | AtU6-26 promoter and 'unarmed' gRNA scaffold | A-B | Ampicillin | | | This work | |
| pGG-B-AtU6-26-BbsI-BbsI-C | AtU6-26 promoter and 'unarmed' gRNA scaffold | B-C | Ampicillin | | | This work | |
| pGG-C-AtU6-26-BbsI-BbsI-D | AtU6-26 promoter and 'unarmed' gRNA scaffold | C-D | Ampicillin | | | This work | |
| pGG-D-AtU6-26-BbsI-BbsI-E | AtU6-26 promoter and 'unarmed' gRNA scaffold | D-E | Ampicillin | | | This work | |
| pGG-E-AtU6-26-BbsI-BbsI-F | AtU6-26 promoter and 'unarmed' gRNA scaffold | E-G | Ampicillin | | | This work | |
| pGG-F-AtU6-26-BbsI-BbsI-G | AtU6-26 promoter and 'unarmed' gRNA scaffold | F-G | Ampicillin | | | This work | |
| pGG-F-AtU6-26-AarI-AarI-G | AtU6-26 promoter and 'unarmed' gRNA scaffold | F-G | Ampicillin | | | This work | |
| **Unarmed gRNA modules** *(ccdB* $^{+}$ *)* | | | | | | | |
| pGG-A-AtU6-26-BbsI-ccdB-BbsI-B | AtU6-26 promoter and 'unarmed' gRNA scaffold | A-B | Ampicillin | | | This work | |
| pGG-B-AtU6-26-BbsI-ccdB-BbsI-C | AtU6-26 promoter and 'unarmed' gRNA scaffold | B-C | Ampicillin | | | This work | |
| pGG-C-AtU6-26-BbsI-ccdB-BbsI-D | AtU6-26 promoter and 'unarmed' gRNA scaffold | C-D | Ampicillin | | | This work | |
| pGG-D-AtU6-26-BbsI-ccdB-BbsI-E | AtU6-26 promoter and 'unarmed' gRNA scaffold | D-E | Ampicillin | | | This work | |
| pGG-E-AtU6-26-BbsI-ccdB-BbsI-F | AtU6-26 promoter and 'unarmed' gRNA scaffold | E-G | Ampicillin | | | This work | |
| pGG-F-AtU6-26-BbsI-ccdB-BbsI-G | AtU6-26 promoter and 'unarmed' gRNA scaffold | F-G | Ampicillin | | | This work | |
| **Armed gRNA modules** | | | | | | | |
| pGG-F-AtU6-26-GFP-1-G | | F-G | Ampicillin | | | This work | |
| **Linkers** | | | | | | | |
| pGG-A-LinkerIII-B | | A-B | Gentamycin | | | This work | |
| pGG-A-LinkerIII-C | | A-C | Gentamycin | | | Houbaert et al. (2018) | |
| pGG-A-LinkerIII-D | | A-D | Gentamycin | | | This work | |
| pGG-A-LinkerIII-E | | A-E | Gentamycin | | | This work | |
| pGG-A-LinkerIII-F | | A-F | Gentamycin | | | This work | |
| pGG-B-Linker-C | | B-C | Ampicillin | | | Lampropoulos et al. (2013) | Addgene ID 48821 |
| pGG-B-linkerII-G | | B-G | Gentamycin | | | This work | |
| pGG-C-LinkerII-G | | C-G | Gentamycin | | | Houbaert et al. (2018) | |
| pGG-D-Linker-E | | D-E | Ampicillin | | | Lampropoulos et al. (2013) | Addgene ID 48834 |
| pGG-D-linkerII-G | | D-G | Gentamycin | | | This work | |
| pGG-E-linkerII-G | | E-G | Gentamycin | | | Houbaert et al. (2018) | |
| pGG-F-linkerII-G | | F-G | Gentamycin | | | This work | |
| **Variable Linkers** | | | | | | | |
| pGG-A-AarI-SacB-AarI-B | contains AarI restriction sites that produce A-B overhangs | A-B | Gentamycin | | | This work | |
| pGG-B-AarI-SacB-AarI-C | contains AarI restriction sites that produce B-C overhangs | B-C | Gentamycin | | | This work | |
| pGG-C-AarI-SacB-AarI-D | contains AarI restriction sites that produce C-D overhangs | C-D | Gentamycin | | | This work | |
| pGG-D-AarI-SacB-AarI-E | contains AarI restriction sites that produce D-E overhangs | D-E | Gentamycin | | | This work | |
| pGG-E-AarI-SacB-AarI-F | contains AarI restriction sites that produce E-F overhangs | E-F | Gentamycin | | | This work | |
| pGG-F-AarI-SacB-AarI-G | contains AarI restriction sites that produce F-G overhangs | F-G | Gentamycin | | | This work | |
| pGG-F-A-AarI-AarI-G-G | contains AarI restriction sites that produce A-G overhangs | F-G | Ampicillin | | | This work | |
| pGG-F-A-AarI-SacB-AarI-G-G | contains AarI restriction sites that produce A-G overhangs | F-G | Ampicillin | | | This work | |

**Supplementary File 3. Plasmid Overview (Continued)**

| Full Name | Type | Cloning sites[a] | Bacterial Selection | Plant Selection | Screenable Marker | Reference | Plasmid Deposit |
|---|---|---|---|---|---|---|---|
| **Expression vector with an empty promoter module** | | | | | | | |
| pFASTR-BsaI-CmR-ccdB-BsaI-Cas9-P2A-mCherry-G7T-AtU6-GFP-1 | | A-B | Spectinomycin | | FAST Red | This work | |
| **One-step cloning vectors compatible with 1 or 2 gRNA's** | | | | | | | |
| pB-pSMB-Cas9-G7T-AtU6-BsaI-BsaI-gRNA scaffold | | | Spectinomycin | Glufosinate-ammonium | | This work | |
| **One-step cloning vectors compatible with 1 or 2 gRNA's** *(ccdB⁺ )* | | | | | | | |
| pFASTR-pSMB-Cas9-P2A-mCherry-G7T-AtU6-BsaI-CmR-ccdB-BsaI-gRNA scaffold | | | Spectinomycin | | FAST Red | This work | |
| pFASTR-pPcUBI-Cas9-P2A-mCherry-G7T-AtU6-BsaI-CmR-ccdB-BsaI-gRNA scaffold | | | Spectinomycin | | FAST Red | This work | |
| pFASTR-pFAMA-Cas9-P2A-mCherry-G7T-AtU6-BsaI-CmR-ccdB-BsaI-sgRNA scaffold | | | Spectinomycin | | FAST Red | This work | |
| pFASTR-pTMM-Cas9-P2A-mCherry-G7T-AtU6-BsaI-CmR-ccdB-BsaI-gRNA scaffold | | | Spectinomycin | | FAST Red | This work | |
| pFASTR-pGATA23-Cas9-P2A-mCherry-G7T-AtU6-BsaI-CmR-ccdB-BsaI-gRNA scaffold | | | Spectinomycin | | FAST Red | This work | |
| pFASTR-pSMB-Cas9-P2A-mTagBFP2-G7T-AtU6-BsaI-CmR-ccdB-BsaI-gRNA scaffold | | | Spectinomycin | | FAST Red | This work | |
| **2x gRNA template** | | | | | | | |
| pEN-2xAtU6 template | | | Kanamycin | | | This work | |
| **One-step cloning vectors compatible with multiple gRNA's** *(ccdB⁺ )* | | | | | | | |
| pFASTR-SMBP-Cas9-P2A-mTagBFP2-G7T-BsaI-ccdB-CmR-BsaI | | A-G | Spectinomycin | | FAST Red | This work | |
| **Expression vectors** | | | | | | | |
| pB-pSMB-Cas9-G7T-AtU6-GFP-1 | | | Spectinomycin | Glufosinate-ammonium | | This work | |
| pB-pSMB-Cas9-G7T-AtU6-GFP-2 | | | Spectinomycin | Glufosinate-ammonium | | This work | |
| pFASTR-pPcUbi-Cas9-P2A-mCherry-G7T-AtU6-GFP-1,PDS3 | | | Spectinomycin | | FAST Red | This work | |
| pFASTR-pPcUBI-Cas9-P2A-mCherry-G7T-AtU6-ARF7-1,ARF19-1 | | | Spectinomycin | | FAST Red | This work | |
| pFASTR-pPcUBI-Cas9-P2A-mCherry-G7T-AtU6-ARF7-2,ARF19-2 | | | Spectinomycin | | FAST Red | This work | |
| pFASTR-pPcUBI-Cas9-P2A-mCherry-G7T-AtU6-YDA-1,YDA-2 | | | Spectinomycin | | FAST Red | This work | |
| pFASTR-pPcUBI-Cas9-P2A-mCherry-G7T-AtU6-CDKA1-1,CDKA1-2 | | | Spectinomycin | | FAST Red | This work | |
| pFASTR-pPcUBI-Cas9-P2A-mCherry-G7T-AtU6-CDKA1-1,CDKB1-1 | | | Spectinomycin | | FAST Red | This work | |
| pFASTR-pPcUBI-Cas9-P2A-mCherry-G7T-AtU6-CDKA1-1,CDKB1-2 | | | Spectinomycin | | FAST Red | This work | |
| pFASTR-pSMB-Cas9-P2A-mCherry-G7T-AtU6-GFP-1 | | | Spectinomycin | | FAST Red | This work | |
| pFASTR-pSMB-Cas9-P2A-mCherry-G7T-AtU6-GFP-1,SMB-1 | | | Spectinomycin | | FAST Red | This work | |
| pFASTR-pSMB-Cas9-P2A-mCherry-G7T-AtU6-GFP-1,SMB-2 | | | Spectinomycin | | FAST Red | This work | |
| pFASTR-pTMM-Cas9-P2A-mCherry-G7T-AtU6-GFP-1,PDS3 | | | Spectinomycin | | FAST Red | This work | |
| pFASTR-pTMM-Cas9-P2A-mCherry-G7T-AtU6-YDA-1,YDA-2 | | | Spectinomycin | | FAST Red | This work | |
| pFASTR-pFAMA-Cas9-P2A-mCherry-G7T-AtU6-GFP-1,PDS3 | | | Spectinomycin | | FAST Red | This work | |
| pFASTR-pGATA23-Cas9-P2A-mCherry-G7T-AtU6-GFP-1 | | | Spectinomycin | | FAST Red | This work | |
| pFASTR-pGATA23-Cas9-P2A-mCherry-G7T-AtU6-ARF7-1,ARF19-1 | | | Spectinomycin | | FAST Red | This work | |
| pFASTR-pGATA23-Cas9-P2A-mCherry-G7T-AtU6-ARF7-2,ARF19-2 | | | Spectinomycin | | FAST Red | This work | |
| pFASTR-pGATA23-Cas9-P2A-mCherry-G7T-AtU6-CDKA1-1,CDKA1-2 | | | Spectinomycin | | FAST Red | This work | |
| pFASTR-pGATA23-Cas9-P2A-mCherry-G7T-AtU6-CDKA1-1,CDKB1-1 | | | Spectinomycin | | FAST Red | This work | |
| pFASTR-pGATA23-Cas9-P2A-mCherry-G7T-AtU6-CDKA1-1,CDKB1-2 | | | Spectinomycin | | FAST Red | This work | |

[a] The cloning sites are either *att* sites for Gateway™ cloning or four-nucleotides overhangs for Golden Gate cloning (according to Lampropoulos et al. 2013)

**Supplementary File 4**

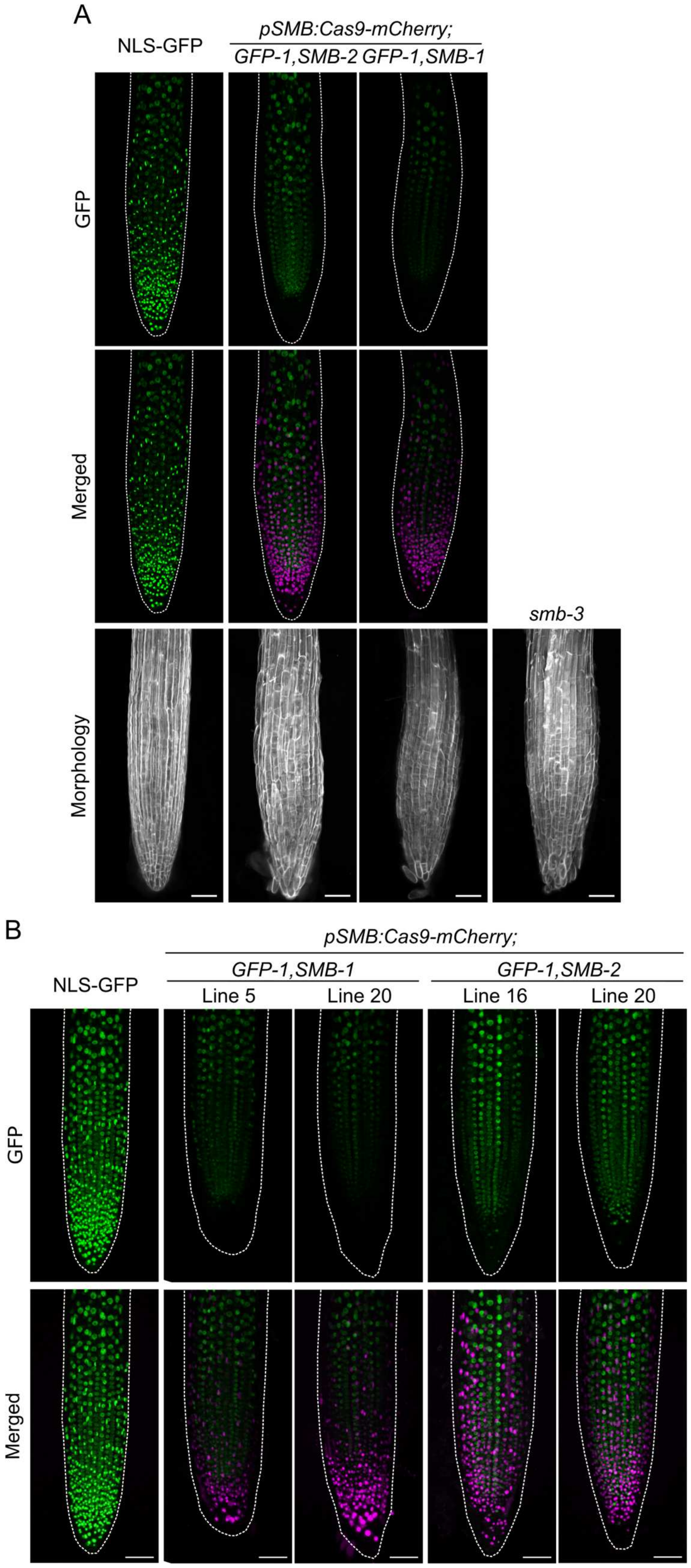

**Supplementary File 4 | *GFP* and *SMB* knockout in root cap at T1 and T2 generations**

**A**, Representative 5 DAG T1 seedlings for both gRNAs targeting *SMB* with simultaneous *GFP* targeting showing root cap specific GFP knockout and root cap specific Cas9-mCherry expression. The typical root cap enlargement observed in *smb* loss of function mutants is shown. Root morphology is shown using DAPI staining. **B**, Representative 6 DAG T2 seedlings targeting both *SMB* and *GFP* in the root cap. Two independent lines for *pSMB:Cas9-mCherry;GFP,SMB-1* and for *pSMB:Cas9-mCherry;GFP,SMB-2* are shown. Disappearance of GFP signal exclusively in root cap nuclei (top) and specific expression of Cas9-mCherry were observed. Accumulation of root cap nuclei observed in CRISPR-TSKO lines is consistent with root cap cell accumulation of a *smb* loss of function. GFP is shown in green and Cas9-mCherry in magenta. All scale bars represent 50 µm.

#### Supplementary File 5

### A

| T1 Line | Promoter | Target | Sorted Population | $R^2$ | TIDE Score (%) | -10 | -9 | -8 | -7 | -6 | -5 | -4 | -3 | -2 | -1 | 0 | 1 | 2 | 3 | 4 | 5 | 6 | 7 | 8 | 9 | 10 |
|---|---|---|---|---|---|---|---|---|---|---|---|---|---|---|---|---|---|---|---|---|---|---|---|---|---|---|
| 13650.2 | SMB | GFP-1 | mCherry | 0.99 | 99.1 | 0.0 | 0.0 | 0.0 | 0.0 | 0.0 | 0.7 | 0.0 | 2.4 | 0.2 | 3.5 | 0.0 | 92.3 | 0.0 | 0.0 | 0.0 | 0.0 | 0.0 | 0.0 | 0.0 | 0.0 | 0.0 |
| 13650.16 | SMB | GFP-1 | mCherry | 0.96 | 95.4 | 0.8 | 0.0 | 0.0 | 0.0 | 0.0 | 0.4 | 0.0 | 1.9 | 0.0 | 3.4 | 0.7 | 86.7 | 0.0 | 0.0 | 0.0 | 1.2 | 0.0 | 0.0 | 0.0 | 0.3 | 0.7 |
| 13715.5 | SMB | GFP-1 | mCherry | 0.97 | 96.7 | 0.0 | 0.0 | 0.0 | 0.0 | 0.0 | 0.0 | 0.0 | 0.0 | 0.0 | 38.3 | 0.0 | 58.3 | 0.0 | 0.0 | 0.0 | 0.0 | 0.0 | 0.1 | 0.0 | 0.0 | 0.0 |
| 13715.12 | SMB | GFP-1 | mCherry | 0.99 | 98.5 | 0.0 | 0.0 | 0.0 | 0.0 | 0.0 | 0.0 | 0.0 | 0.7 | 0.0 | 3.0 | 0.0 | 94.8 | 0.0 | 0.0 | 0.0 | 0.0 | 0.0 | 0.0 | 0.0 | 0.0 | 0.0 |
| 13650.2 | SMB | GFP-1 | GFP | 0.99 | 2.9 | 0.0 | 0.0 | 0.1 | 0.0 | 0.5 | 0.0 | 0.0 | 0.0 | 0.0 | 0.0 | 95.9 | 1.1 | 0.1 | 0.0 | 0.4 | 0.0 | 0.6 | 0.0 | 0.0 | 0.0 | 0.0 |
| 13650.16 | SMB | GFP-1 | GFP | 0.99 | 1.3 | 0.0 | 0.2 | 0.0 | 0.0 | 0.7 | 0.0 | 0.0 | 0.0 | 0.0 | 0.0 | 97.4 | 0.0 | 0.0 | 0.0 | 0.0 | 0.0 | 0.3 | 0.0 | 0.0 | 0.0 | 0.0 |
| 13715.5 | SMB | GFP-1 | GFP | 0.99 | 3.6 | 0.0 | 0.0 | 0.0 | 0.0 | 0.0 | 0.0 | 0.0 | 0.1 | 0.0 | 0.9 | 95.5 | 2.3 | 0.2 | 0.0 | 0.0 | 0.0 | 0.0 | 0.0 | 0.0 | 0.0 | 0.1 |
| 13715.12 | SMB | GFP-1 | GFP | 0.99 | 3.1 | 0.0 | 0.0 | 0.0 | 0.0 | 0.0 | 0.4 | 0.0 | 0.0 | 0.0 | 0.0 | 96.0 | 2.4 | 0.0 | 0.0 | 0.2 | 0.0 | 0.1 | 0.0 | 0.0 | 0.0 | 0.0 |

### B

| T1 Line | Promoter | Target | Sorted Population | $R^2$ | TIDE Score (%) | -10 | -9 | -8 | -7 | -6 | -5 | -4 | -3 | -2 | -1 | 0 | 1 | 2 | 3 | 4 | 5 | 6 | 7 | 8 | 9 | 10 |
|---|---|---|---|---|---|---|---|---|---|---|---|---|---|---|---|---|---|---|---|---|---|---|---|---|---|---|
| 13650.2 | SMB | SMB-1 | mCherry | 0.98 | 98.4 | 0.1 | 2.1 | 0.0 | 0.0 | 0.0 | 1.7 | 3.8 | 0.9 | 1.8 | 6.3 | 0.0 | 80.5 | 0.0 | 0.0 | 0.0 | 0.0 | 0.0 | 0.0 | 1.2 | 0.0 | 0.0 |
| 13650.16 | SMB | SMB-1 | mCherry | 0.98 | 98.1 | 0.0 | 4.7 | 0.0 | 0.3 | 0.0 | 1.4 | 2.3 | 0.7 | 0.4 | 9.9 | 0.0 | 78.2 | 0.0 | 0.2 | 0.0 | 0.0 | 0.0 | 0.0 | 0.0 | 0.0 | 0.0 |
| 13650.2 | SMB | SMB-1 | GFP | 0.99 | 5.1 | 0.6 | 0.0 | 0.0 | 0.0 | 0.9 | 0.5 | 1.0 | 0.0 | 0.1 | 0.0 | 93.5 | 1.2 | 0.8 | 0.0 | 0.0 | 0.0 | 0.0 | 0.0 | 0.0 | 0.0 | 0.0 |
| 13650.16 | SMB | SMB-1 | GFP | 1.00 | 1.3 | 0.3 | 0.0 | 0.0 | 0.0 | 0.3 | 0.3 | 0.2 | 0.0 | 0.0 | 0.1 | 98.4 | 0.2 | 0.0 | 0.0 | 0.0 | 0.0 | 0.0 | 0.0 | 0.0 | 0.0 | 0.0 |

### C

| T1 Line | Promoter | Target | Sorted Population | $R^2$ | TIDE Score (%) | -10 | -9 | -8 | -7 | -6 | -5 | -4 | -3 | -2 | -1 | 0 | 1 | 2 | 3 | 4 | 5 | 6 | 7 | 8 | 9 | 10 |
|---|---|---|---|---|---|---|---|---|---|---|---|---|---|---|---|---|---|---|---|---|---|---|---|---|---|---|
| 13715.5 | SMB | SMB-2 | mCherry | 0.97 | 97.4 | 0.0 | 0.0 | 0.0 | 0.0 | 0.0 | 0.0 | 0.0 | 9.0 | 0.0 | 0.7 | 0.0 | 87.8 | 0.0 | 0.0 | 0.0 | 0.0 | 0.0 | 0.0 | 0.0 | 0.0 | 0.0 |
| 13715.12 | SMB | SMB-2 | mCherry | 0.97 | 96.5 | 0.0 | 1.5 | 0.0 | 0.0 | 0.0 | 0.0 | 0.0 | 10.9 | 0.3 | 4.3 | 0.4 | 79.4 | 0.0 | 0.0 | 0.0 | 0.0 | 0.0 | 0.0 | 0.0 | 0.0 | 0.0 |
| 13715.5 | SMB | SMB-2 | GFP | 0.98 | 2.7 | 0.0 | 0.0 | 0.0 | 0.0 | 0.0 | 0.0 | 0.1 | 0.5 | 0.7 | 0.3 | 95.2 | 0.8 | 0.0 | 0.3 | 0.0 | 0.0 | 0.0 | 0.0 | 0.0 | 0.0 | 0.0 |
| 13715.12 | SMB | SMB-2 | GFP | 0.98 | 1.1 | 0.0 | 0.0 | 0.0 | 0.0 | 0.0 | 0.0 | 0.0 | 0.0 | 0.0 | 0.0 | 97.1 | 0.9 | 0.0 | 0.0 | 0.0 | 0.0 | 0.0 | 0.0 | 0.0 | 0.1 | 0.0 |

**Supplementary File 5 | Indel profile of protoplast from T2 root tips**

TIDE analysis of *pSMB:*Cas9-mCherry;GFP-1,SMB-1 (Line 2 and 16) and *pSMB:*Cas9-mCherry;GFP-1,SMB-2 (Line 5 and 12). Parameters TIDE analysis: **A**, Target: *GFP-1* (expected cut at 179 bp), Alignment window: 100-169, Decomposition window: 194-400, Indel size: 10. **B**, Target: *SMB-1* (expected cut at 197 bp), Alignment window: 100-187, Decomposition window: 212-245, Indel size: 10. **C**, Target: *SMB-2* (expected cut at 315 bp), Alignment window: 100-305, Decomposition window: 330-470, Indel size: 10. Significant values ($p<0.001$) are highlighted in green.

#### Supplementary File 6

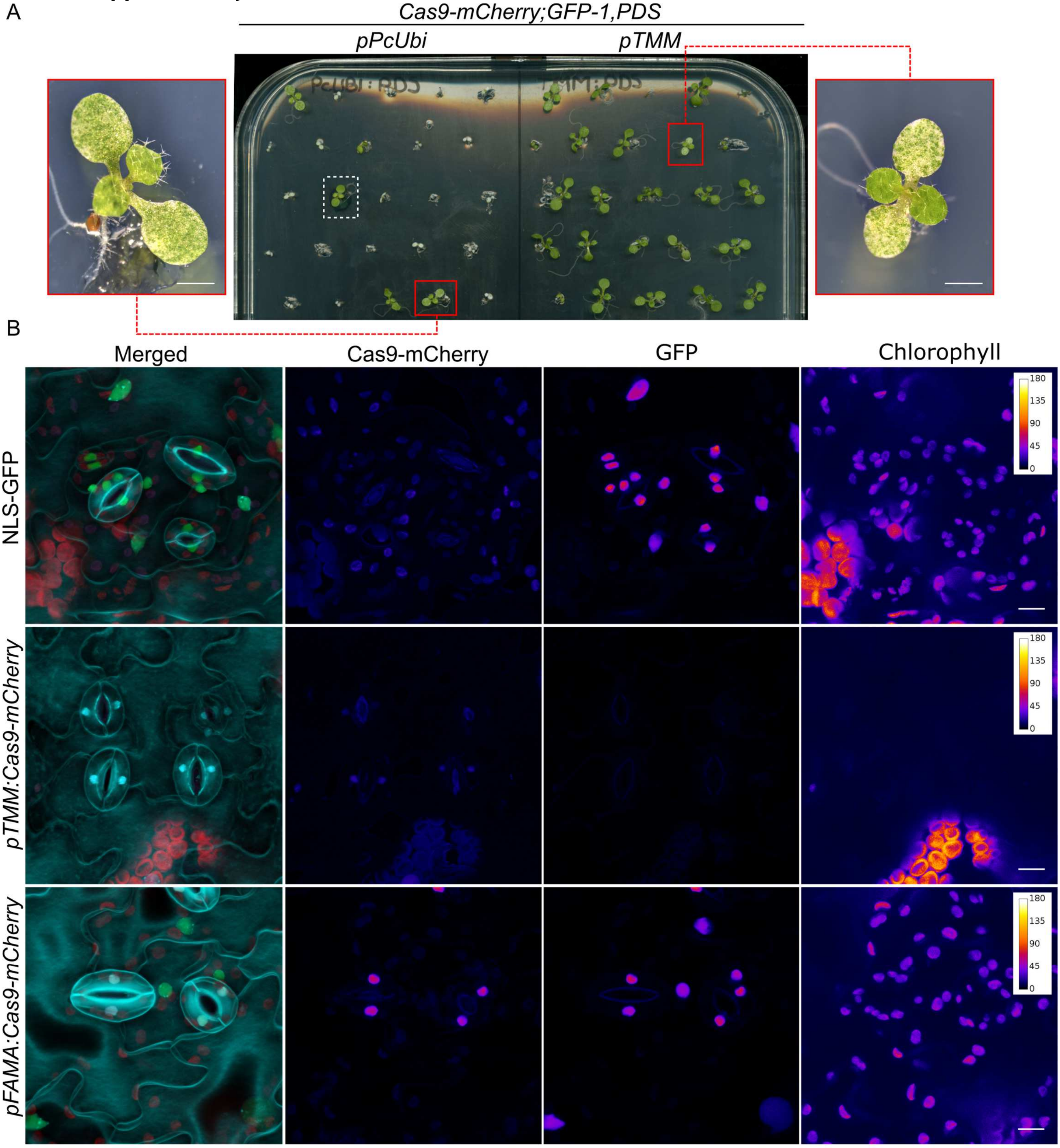

**Supplementary File 6 | Gene knockout in stomatal lineage**

**A**, T1 seedlings targeting *GFP* and *PDS3* simultaneously. Dashed white rectangle indicates a FAST negative seedling. Red boxes show chimeric bleached seedlings. Scale bars are 1mm. **B**, Simultaneous stomata lineage specific knockout of GFP and chlorophyll biosynthesis in 10DAG T1 seedlings. Shown are stomata at the abaxial face of cotyledons. While both GFP and chlorophyll signal is lost in stomata in lines under the control of *pTMM*, signal is still present in stomata in lines under the control of *pFAMA*. At 10 DAG, Cas9-mCherry signal is often weak or missing in *pTMM:Cas9-mCherry;GFP,PDS3* lines likely to decreased activity of the *pTMM* promoter fragment in stomata at this developmental stage. In merged image, cell outlines are in cyan, GFP in green, Cas9-mCherry in magenta, and chlorophyll fluorescence in red. Epidermal morphology is shown using DAPI staining. Scale bars represent 10µm.

#### Supplementary File 7

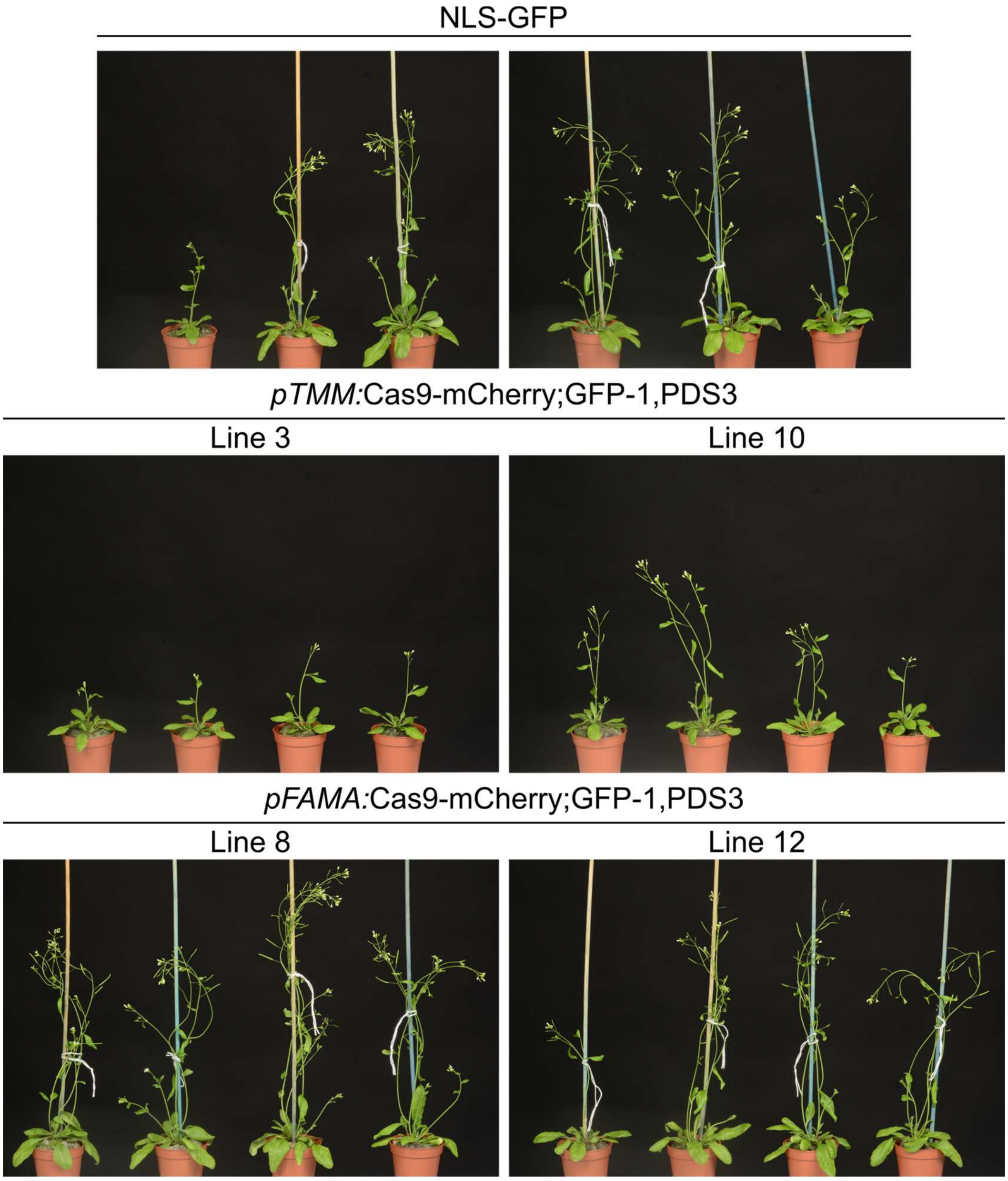

**Supplemental File 7 | CRISPR-TSKO of *GFP* and *PDS3* leads to viable plants**

Soil grown T2 plants of CRISPR-TSKO under control of *pTMM* and *pFAMA* targeting both *GFP* and *PDS3* in comparison to background line NLS-GFP. All plants are of same age.

#### Supplementary File 8

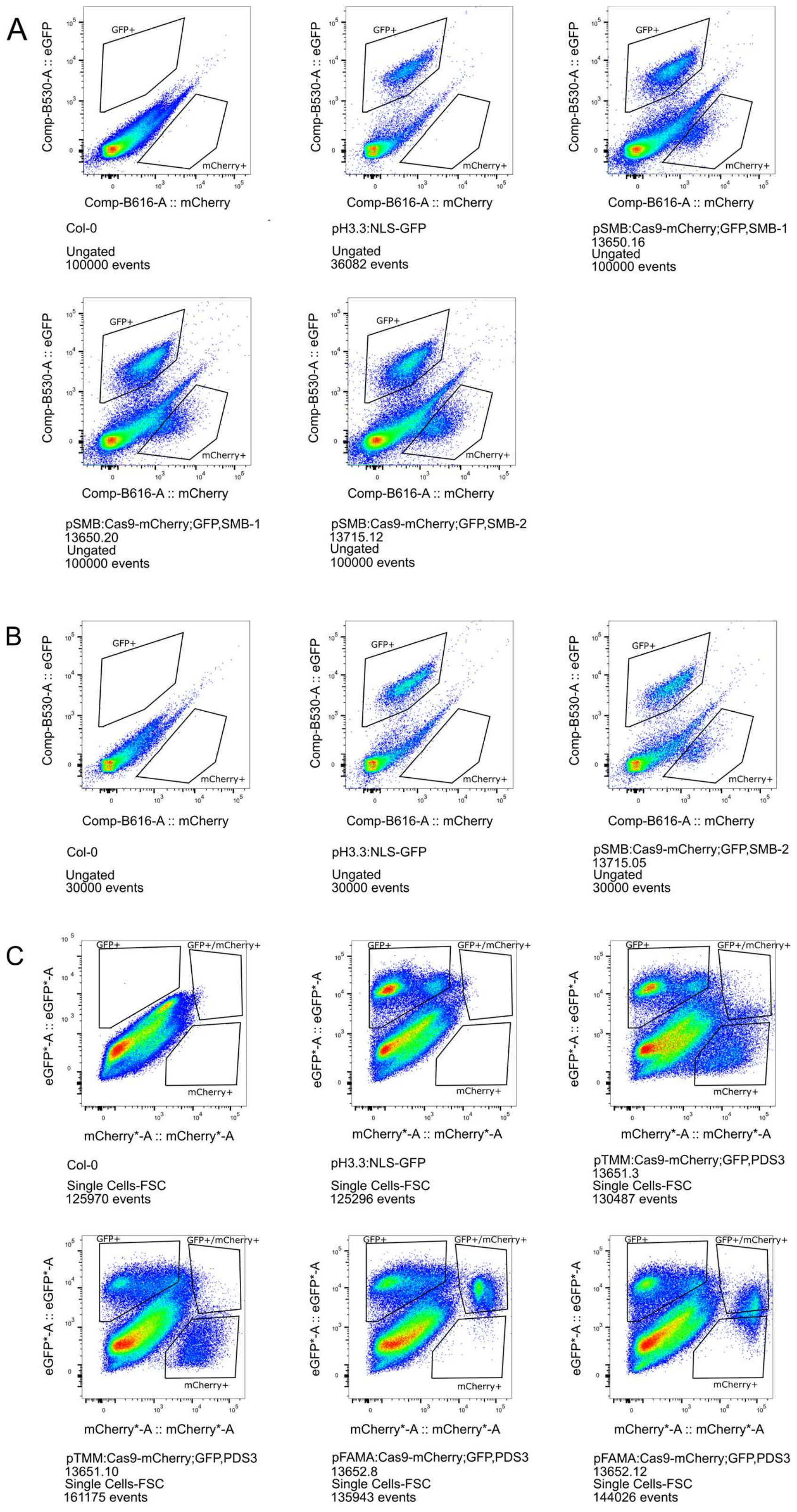

**Supplementary File 8 | Fluorescence-activated Cell Sorting of protoplasts used in genotyping**

Gating strategies for protoplasts from root tips (**A,B**) and cotyledons (**C**). Collected populations were used for DNA extraction and molecular analysis shown in **Supplementary File 9**.

### Supplementary File 9

## A

| T1 Line | Promoter | Target | Sorted Population | $R^2$ | TIDE Score (%) | -10 | -9 | -8 | -7 | -6 | -5 | -4 | -3 | -2 | -1 | 0 | 1 | 2 | 3 | 4 | 5 | 6 | 7 | 8 | 9 | 10 |
|---|---|---|---|---|---|---|---|---|---|---|---|---|---|---|---|---|---|---|---|---|---|---|---|---|---|---|
| 13651.3 | TMM | GFP-1 | mCherry | 0.99 | 78.6 | 0.0 | 0.0 | 0.0 | 0.0 | 0.0 | 0.0 | 0.0 | 0.5 | 0.0 | 0.0 | 20.0 | 78.1 | 0.0 | 0.0 | 0.0 | 0.0 | 0.0 | 0.0 | 0.0 | 0.0 | 0.0 |
| 13651.10 | TMM | GFP-1 | mCherry | - | - | | | | | | | | | | | | | | | | | | | | | |
| 13652.8 | FAMA | GFP-1 | mCherry/GFP | 0.98 | 30.0 | 0.0 | 0.0 | 0.0 | 0.0 | 0.0 | 0.0 | 0.0 | 3.0 | 0.0 | 1.2 | 68.4 | 21.0 | 4.8 | 0.0 | 0.0 | 0.0 | 0.0 | 0.0 | 0.0 | 0.0 | 0.0 |
| 13652.12 | FAMA | GFP-1 | mCherry/GFP | 0.98 | 66.3 | 0.0 | 0.0 | 0.0 | 0.0 | 0.0 | 0.3 | 0.6 | 4.7 | 1.8 | 4.1 | 31.6 | 54.2 | 0.0 | 0.0 | 0.0 | 0.0 | 0.5 | 0.0 | 0.0 | 0.0 | 0.0 |
| 13651.3 | TMM | GFP-1 | GFP | 0.99 | 0.2 | 0.0 | 0.0 | 0.0 | 0.0 | 0.2 | 0.0 | 0.0 | 0.0 | 0.0 | 0.0 | 98.8 | 0.0 | 0.0 | 0.0 | 0.0 | 0.0 | 0.0 | 0.0 | 0.0 | 0.0 | 0.0 |
| 13651.10 | TMM | GFP-1 | GFP | 0.99 | 0.5 | 0.0 | 0.0 | 0.0 | 0.0 | 0.0 | 0.0 | 0.0 | 0.0 | 0.5 | 0.0 | 98.3 | 0.0 | 0.0 | 0.0 | 0.0 | 0.0 | 0.0 | 0.0 | 0.0 | 0.0 | 0.0 |
| 13652.8 | FAMA | GFP-1 | GFP | 0.90 | 1.5 | 0.0 | 0.0 | 0.0 | 1.1 | 0.0 | 0.0 | 0.0 | 0.0 | 0.0 | 0.0 | 88.8 | 0.0 | 0.0 | 0.0 | 0.0 | 0.0 | 0.0 | 0.0 | 0.0 | 0.4 | 0.0 |
| 13652.12 | FAMA | GFP-1 | GFP | 0.99 | 1.3 | 0.0 | 0.0 | 0.0 | 0.0 | 0.1 | 0.1 | 0.0 | 0.0 | 0.8 | 0.0 | 97.7 | 0.1 | 0.0 | 0.0 | 0.3 | 0.0 | 0.0 | 0.0 | 0.0 | 0.0 | 0.0 |

## B

| T1 Line | Promoter | Target | Sorted Population | $R^2$ | TIDE Score (%) | -10 | -9 | -8 | -7 | -6 | -5 | -4 | -3 | -2 | -1 | 0 | 1 | 2 | 3 | 4 | 5 | 6 | 7 | 8 | 9 | 10 |
|---|---|---|---|---|---|---|---|---|---|---|---|---|---|---|---|---|---|---|---|---|---|---|---|---|---|---|
| 13651.3 | TMM | PDS | mCherry | 0.96 | 84.9 | 1.2 | 0.4 | 3.7 | 8.4 | 0.4 | 0.3 | 0.0 | 0.0 | 0.0 | 5.8 | 11.5 | 60.4 | 0.0 | 0.0 | 0.0 | 2.4 | 0.0 | 0.0 | 0.8 | 0.6 | 0.6 |
| 13651.10 | TMM | PDS | mCherry | 0.95 | 74.4 | 0.0 | 0.4 | 0.0 | 0.0 | 1.0 | 1.2 | 0.0 | 0.0 | 0.0 | 12.8 | 21.0 | 56.7 | 0.0 | 0.0 | 0.0 | 0.0 | 0.0 | 0.0 | 0.9 | 1.5 | 0.0 |
| 13652.8 | FAMA | PDS | mCherry/GFP | 0.96 | 49.1 | 0.3 | 0.4 | 0.9 | 1.4 | 0.0 | 0.0 | 0.0 | 0.5 | 0.3 | 6.1 | 46.6 | 37.8 | 0.0 | 0.0 | 0.0 | 0.0 | 0.0 | 0.5 | 0.2 | 0.5 | 0.1 |
| 13652.12 | FAMA | PDS | mCherry/GFP | 0.98 | 75.3 | 0.0 | 0.0 | 0.0 | 3.7 | 0.3 | 0.6 | 0.0 | 0.0 | 0.0 | 8.5 | 22.5 | 62.3 | 0.0 | 0.0 | 0.0 | 0.0 | 0.0 | 0.0 | 0.0 | 0.0 | 0.0 |
| 13651.3 | TMM | PDS | GFP | 0.98 | 8.8 | 0.0 | 0.3 | 0.3 | 1.1 | 0.7 | 0.0 | 1.2 | 0.0 | 1.3 | 0.7 | 89.5 | 0.6 | 0.0 | 0.0 | 0.2 | 0.0 | 0.6 | 0.0 | 0.5 | 0.4 | 0.8 |
| 13651.10 | TMM | PDS | GFP | 0.99 | 7.5 | 0.4 | 0.2 | 0.7 | 0.1 | 0.6 | 0.0 | 1.4 | 0.0 | 1.0 | 0.1 | 91.2 | 0.0 | 0.0 | 0.0 | 0.2 | 0.0 | 1.0 | 0.5 | 0.0 | 0.7 | 0.5 |
| 13652.8 | FAMA | PDS | GFP | 0.98 | 6.8 | 0.0 | 1.4 | 0.7 | 0.7 | 0.0 | 0.0 | 0.8 | 0.3 | 0.2 | 0.0 | 91.7 | 0.0 | 0.0 | 0.0 | 0.4 | 0.0 | 0.3 | 0.5 | 0.5 | 0.4 | 0.5 |
| 13652.12 | FAMA | PDS | GFP | 0.92 | 8.3 | 0.0 | 0.0 | 0.0 | 0.0 | 2.4 | 1.6 | 0.8 | 0.0 | 1.5 | 0.4 | 83.4 | 0.0 | 0.0 | 0.0 | 0.0 | 0.5 | 1.2 | 0.0 | 0.0 | 0.0 | 0.0 |

#### Supplementary File 9 | Indel profile of protoplast enriched for guard cells of T2 plants

TIDE analysis of *pTMM*:Cas9-mCherry;GFP-1,PDS3 (line 3 and 10) and *pFAMA*:Cas9-mCherry;GFP-1,PDS3 (line 3 and 10). Parameters TIDE analysis: **A**, Target: *GFP-1* (expected cut at 179 bp), Alignment window: 100-169, Decomposition window: 194-400, Indel size: 10. **B**, Target: *PDS3* (expected cut at 291 bp), Alignment window: 100-281, Decomposition window: 306-400, Indel size: 10. Significant values (p<0.001) are highlighted in green.

#### Supplementary File 10

**A**

| T1 Line | Promoter | Target | Sorted Population | $R^2$ | TIDE Score (%) | -10 | -9 | -8 | -7 | -6 | -5 | -4 | -3 | -2 | -1 | 0 | 1 | 2 | 3 | 4 | 5 | 6 | 7 | 8 | 9 | 10 |
|---|---|---|---|---|---|---|---|---|---|---|---|---|---|---|---|---|---|---|---|---|---|---|---|---|---|---|
| 13652.2 | FAMA | GFP-1 | mCherry/GFP | 0.98 | 29.1 | 0.0 | 0.2 | 0.0 | 0.0 | 0.0 | 0.0 | 0.0 | 2.1 | 0.2 | 3.8 | 69.1 | 22.8 | 0.0 | 0.0 | 0.0 | 0.0 | 0.0 | 0.0 | 0.0 | 0.0 | 0.0 |
| 13652.7 | FAMA | GFP-1 | mCherry/GFP | 0.98 | 22.7 | 0.0 | 0.3 | 0.0 | 0.0 | 0.0 | 0.0 | 0.0 | 2.0 | 0.0 | 4.2 | 75.8 | 16.2 | 0.0 | 0.0 | 0.0 | 0.0 | 0.0 | 0.0 | 0.0 | 0.0 | 0.0 |
| 13652.8 | FAMA | GFP-1 | mCherry/GFP | 0.98 | 37.8 | 0.0 | 0.4 | 0.0 | 0.0 | 0.0 | 0.1 | 0.0 | 2.2 | 0.2 | 5.3 | 60.6 | 29.2 | 0.0 | 0.0 | 0.0 | 0.0 | 0.3 | 0.0 | 0.0 | 0.0 | 0.0 |
| 13652.12 | FAMA | GFP-1 | mCherry/GFP | 0.98 | 74.7 | 0.0 | 0.0 | 0.1 | 0.0 | 0.0 | 0.0 | 0.0 | 3.4 | 0.4 | 5.8 | 23.3 | 64.1 | 0.0 | 0.4 | 0.2 | 0.0 | 0.0 | 0.0 | 0.0 | 0.4 | 0.0 |
| 13652.2 | FAMA | GFP-1 | GFP | 0.99 | 1.3 | 0.0 | 0.0 | 0.0 | 0.0 | 0.0 | 0.0 | 0.0 | 0.1 | 0.0 | 0.6 | 97.5 | 0.0 | 0.0 | 0.0 | 0.0 | 0.0 | 0.3 | 0.0 | 0.0 | 0.2 | 0.0 |
| 13652.7 | FAMA | GFP-1 | GFP | 0.99 | 0.9 | 0.0 | 0.1 | 0.0 | 0.0 | 0.0 | 0.0 | 0.0 | 0.0 | 0.0 | 0.2 | 98.0 | 0.0 | 0.1 | 0.0 | 0.0 | 0.0 | 0.4 | 0.0 | 0.1 | 0.0 | 0.0 |
| 13652.8 | FAMA | GFP-1 | GFP | 0.99 | 0.5 | 0.0 | 0.3 | 0.0 | 0.0 | 0.0 | 0.0 | 0.0 | 0.0 | 0.0 | 0.0 | 98.4 | 0.0 | 0.0 | 0.0 | 0.0 | 0.0 | 0.0 | 0.0 | 0.2 | 0.0 | 0.0 |
| 13652.12 | FAMA | GFP-1 | GFP | 0.99 | 1.6 | 0.0 | 0.1 | 0.0 | 0.0 | 0.0 | 0.0 | 0.0 | 0.2 | 0.1 | 0.5 | 97.6 | 0.0 | 0.0 | 0.0 | 0.0 | 0.0 | 0.7 | 0.0 | 0.0 | 0.0 | 0.0 |

**B**

| T1 Line | Promoter | Target | Sorted Population | $R^2$ | TIDE Score (%) | -10 | -9 | -8 | -7 | -6 | -5 | -4 | -3 | -2 | -1 | 0 | 1 | 2 | 3 | 4 | 5 | 6 | 7 | 8 | 9 | 10 |
|---|---|---|---|---|---|---|---|---|---|---|---|---|---|---|---|---|---|---|---|---|---|---|---|---|---|---|
| 13652.2 | FAMA | PDS | mCherry/GFP | 0.98 | 42.0 | 0.1 | 0.0 | 0.0 | 0.9 | 1.4 | 0.0 | 0.0 | 0.0 | 0.0 | 4.0 | 56.1 | 35.5 | 0.0 | 0.0 | 0.0 | 0.0 | 0.0 | 0.0 | 0.1 | 0.0 | 0.0 |
| 13652.7 | FAMA | PDS | mCherry/GFP | 0.99 | 34.4 | 0.3 | 0.3 | 0.0 | 2.0 | 1.4 | 0.0 | 0.8 | 1.0 | 1.6 | 5.0 | 67.2 | 17.5 | 0.0 | 0.0 | 0.0 | 0.2 | 0.6 | 0.2 | 0.0 | 0.0 | 0.5 |
| 13652.8 | FAMA | PDS | mCherry/GFP | 0.97 | 55.8 | 0.5 | 0.3 | 0.2 | 0.2 | 1.6 | 0.1 | 0.0 | 0.0 | 0.0 | 7.5 | 41.5 | 45.3 | 0.0 | 0.0 | 0.0 | 0.0 | 0.0 | 0.0 | 0.0 | 0.1 | 0.0 |
| 13652.12 | FAMA | PDS | mCherry/GFP | 0.98 | 85.7 | 0.2 | 0.5 | 1.8 | 6.9 | 0.4 | 1.3 | 0.8 | 1.1 | 0.8 | 8.0 | 12.4 | 62.0 | 0.0 | 0.0 | 0.0 | 1.2 | 0.3 | 0.5 | 0.0 | 0.0 | 0.0 |
| 13652.2 | FAMA | PDS | GFP | 0.99 | 1.6 | 0.1 | 0.0 | 0.0 | 0.4 | 0.3 | 0.0 | 0.0 | 0.0 | 0.0 | 0.7 | 97.5 | 0.0 | 0.0 | 0.0 | 0.0 | 0.0 | 0.1 | 0.0 | 0.0 | 0.0 | 0.0 |
| 13652.7 | FAMA | PDS | GFP | 0.98 | 5.7 | 0.3 | 0.6 | 0.0 | 0.5 | 1.6 | 0.2 | 0.0 | 0.2 | 0.1 | 0.5 | 92.7 | 0.0 | 0.0 | 0.0 | 0.2 | 0.0 | 1.1 | 0.0 | 0.0 | 0.0 | 0.4 |
| 13652.8 | FAMA | PDS | GFP | 0.99 | 5.9 | 0.1 | 0.4 | 0.0 | 0.7 | 0.7 | 0.4 | 0.4 | 0.2 | 1.2 | 0.4 | 93.1 | 0.0 | 0.0 | 0.0 | 0.4 | 0.1 | 0.4 | 0.0 | 0.1 | 0.3 | 0.2 |
| 13652.12 | FAMA | PDS | GFP | 0.90 | 5.1 | 0.0 | 0.3 | 0.1 | 0.0 | 1.6 | 0.5 | 0.1 | 0.0 | 0.0 | 0.2 | 93.8 | 0.0 | 0.0 | 0.0 | 0.9 | 0.7 | 0.7 | 0.0 | 0.0 | 0.0 | 0.0 |

**Supplementary File 10 | Indel profile of protoplast enriched for guard cells of *pFAMA*: Cas9-mCherry;GFP-1,PDS3 lines**

TIDE analysis of pFAMA:Cas9-mCherry;GFP-1,PDS3 (Lines 2, 7, 8 and 12). Parameters TIDE analysis: **A**, Target: *GFP-1* (expected cut at 179 bp), Alignment window: 100-169, Decomposition window: 194-400, Indel size: 10. **B**, Target: *PDS3* (expected cut at 289 bp), Alignment window: 100-279, Decomposition window: 304-400, Indel size: 10. Significant values ($p<0.001$) are highlighted in green.

#### Supplementary File 11

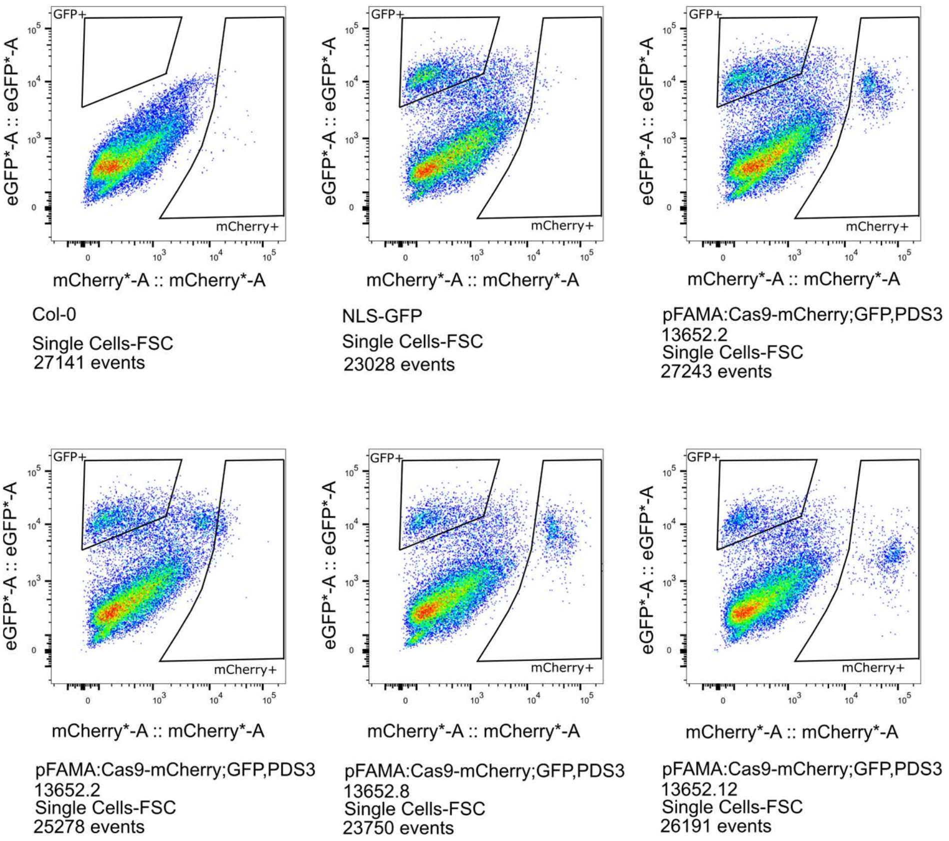

**Supplementary File 11 | Fluorescence-activated Cell Sorting of protoplasts of *pFAMA*: Cas9-mCherry;GFP-1,PDS3 lines**

Gating strategies for protoplasts from cotyledons. Collected populations were used for DNA extraction and molecular analysis shown in **Supplementary File 10**.

#### Supplementary File 12

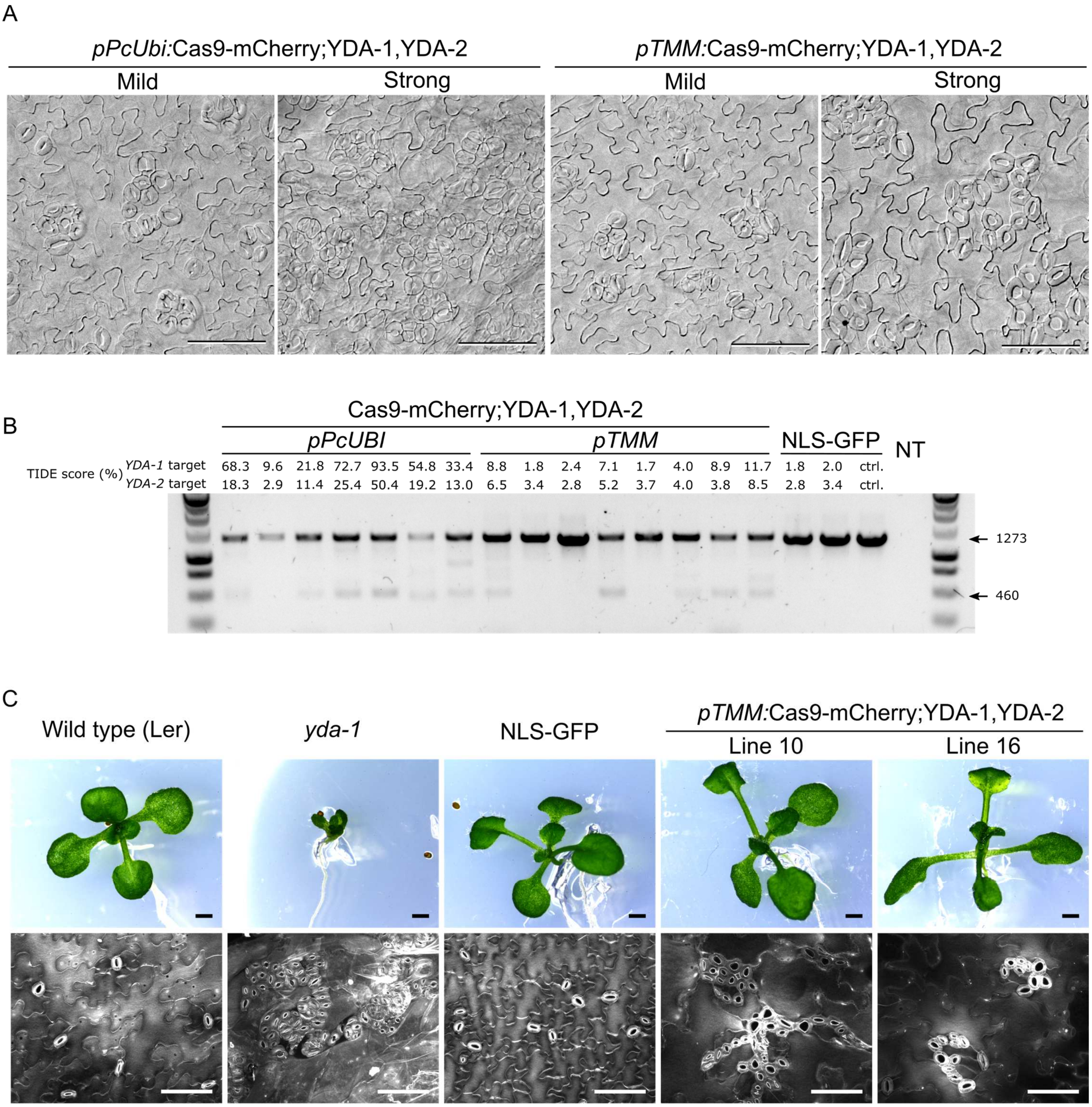

#### Supplementary File 12 | Stomatal lineage specific knockout of *YDA* with CRISPR-TSKO

**A**, Range of stomatal clustering phenotypes is shown on the abaxial surface of cotyledons of 10 day old T1 seedlings for both ubiquitous (*pPcUbi*) and stomatal lineage (*pTMM*) constructs. Scale bars are 100 µm. **B**, Genotyping results for T1 seedlings of both ubiquitous (*pPcUbi*) and stomatal lineage (*pTMM*) constructs. Lower band in gel is indicative of large deletion in *YDA*. TIDE scores are shown for upper band for both gRNAs used. **C**, representative 15 day old seedlings (up) and corresponding projection of adaxial face of cotyledon showing stomata and pavement epidermal cells. Wild type (Ler) is the background line for *yda-1.* NLS-GFP is the wild type background for both Line 10 and Line 16. Scale represent 1mm and 100 µm.

### Supplementary File 13

## A

| Sample | Promoter | Target | $R^2$ | TIDE Score (%) | -18 | -17 | -16 | -15 | -14 | -5 | -4 | -3 | -2 | -1 | 0 | 1 | 2 | 13 | 14 | 15 | 16 |
|---|---|---|---|---|---|---|---|---|---|---|---|---|---|---|---|---|---|---|---|---|---|
| 13984.1 | PcUBI | YDA-1 | 0.91 | 68.3 | 2.1 | 0.1 | 0.0 | 0.4 | 0.4 | 1.3 | 0.9 | 7.8 | 3.0 | 0.5 | 22.8 | 42.9 | 0.0 | 0.3 | 0.1 | 0.0 | 2.5 |
| 13984.2 | PcUBI | YDA-1 | 0.98 | 9.6 | 0.7 | 0.0 | 0.8 | 0.0 | 0.8 | 0.0 | 0.7 | 0.0 | 0.2 | 0.0 | 88.1 | 0.0 | 0.0 | 0.1 | 0.5 | 0.3 | 0.6 |
| 13984.3 | PcUBI | YDA-1 | 0.96 | 21.8 | 2.8 | 0.2 | 0.0 | 0.0 | 0.2 | 0.6 | 1.5 | 0.6 | 0.8 | 3.5 | 74.7 | 11.4 | 0.0 | 0.0 | 0.0 | 0.0 | 0.1 |
| 13984.4 | PcUBI | YDA-1 | 0.96 | 72.7 | 2.0 | 0.1 | 0.1 | 0.6 | 0.0 | 1.4 | 2.4 | 1.9 | 2.7 | 4.8 | 23.4 | 50.7 | 0.0 | 0.0 | 0.0 | 0.1 | 0.9 |
| 13984.5 | PcUBI | YDA-1 | 0.97 | 93.5 | 0.0 | 0.7 | 0.0 | 2.0 | 0.2 | 0.0 | 0.6 | 3.5 | 2.8 | 3.8 | 3.5 | 72.0 | 0.0 | 0.8 | 0.4 | 0.5 | 0.8 |
| 13984.6 | PcUBI | YDA-1 | 0.90 | 54.8 | 3.1 | 0.0 | 1.0 | 0.4 | 3.7 | 0.0 | 1.0 | 1.7 | 13.7 | 0.0 | 35.6 | 22.1 | 0.7 | 3.4 | 0.0 | 0.0 | 0.7 |
| 13984.7 | PcUBI | YDA-1 | 0.91 | 33.4 | 3.5 | 0.0 | 0.7 | 0.1 | 0.0 | 0.2 | 0.5 | 0.0 | 0.0 | 2.8 | 57.9 | 14.2 | 0.0 | 0.0 | 0.0 | 0.9 | 0.7 |
| 13983.8 | TMM | YDA-1 | 0.98 | 8.8 | 1.2 | 0.0 | 0.1 | 0.0 | 0.0 | 0.0 | 0.1 | 0.0 | 0.4 | 0.0 | 89.3 | 5.4 | 0.0 | 0.0 | 0.0 | 0.0 | 0.3 |
| 13983.9 | TMM | YDA-1 | 0.99 | 1.8 | 0.8 | 0.0 | 0.0 | 0.0 | 0.0 | 0.0 | 0.2 | 0.0 | 0.0 | 0.0 | 97.0 | 0.0 | 0.2 | 0.0 | 0.0 | 0.0 | 0.2 |
| 13983.10 | TMM | YDA-1 | 0.99 | 2.4 | 0.6 | 0.0 | 0.0 | 0.0 | 0.0 | 0.0 | 0.0 | 0.0 | 0.6 | 0.0 | 96.9 | 0.0 | 0.3 | 0.0 | 0.0 | 0.2 | 0.3 |
| 13983.11 | TMM | YDA-1 | 0.98 | 7.1 | 0.4 | 0.0 | 0.0 | 0.0 | 0.0 | 0.0 | 0.9 | 0.2 | 1.0 | 0.2 | 91.3 | 3.8 | 0.3 | 0.0 | 0.0 | 0.0 | 0.3 |
| 13983.12 | TMM | YDA-1 | 0.99 | 1.7 | 0.3 | 0.0 | 0.0 | 0.0 | 0.0 | 0.0 | 0.2 | 0.0 | 0.2 | 0.0 | 97.3 | 0.0 | 0.0 | 0.0 | 0.0 | 0.0 | 0.3 |
| 13983.13 | TMM | YDA-1 | 0.98 | 4.0 | 1.0 | 0.0 | 0.0 | 0.0 | 0.0 | 0.0 | 0.5 | 0.0 | 0.0 | 0.0 | 94.2 | 0.6 | 0.0 | 0.0 | 0.0 | 0.0 | 0.7 |
| 13983.14 | TMM | YDA-1 | 0.98 | 8.9 | 0.8 | 0.0 | 0.6 | 0.0 | 0.1 | 0.3 | 0.7 | 0.0 | 1.0 | 0.0 | 89.4 | 0.0 | 0.2 | 0.0 | 0.0 | 0.0 | 0.6 |
| 13983.15 | TMM | YDA-1 | 0.94 | 11.7 | 2.0 | 0.9 | 0.1 | 0.0 | 0.0 | 0.7 | 0.3 | 0.5 | 0.0 | 0.0 | 82.7 | 3.9 | 0.7 | 0.0 | 0.0 | 0.7 | 0.0 |
| NLS-GFP | | | 0.99 | 1.8 | 0.5 | 0.0 | 0.0 | 0.0 | 0.0 | 0.0 | 0.0 | 0.0 | 0.0 | 0.0 | 97.3 | 0.0 | 0.0 | 0.0 | 0.0 | 0.0 | 0.4 |
| NLS-GFP | | | 0.99 | 2.0 | 0.6 | 0.0 | 0.1 | 0.0 | 0.0 | 0.0 | 0.3 | 0.0 | 0.1 | 0.0 | 97.2 | 0.0 | 0.0 | 0.0 | 0.0 | 0.1 | 0.2 |

## B

| Sample | Promoter | Target | $R^2$ | TIDE Score (%) | -10 | -9 | -8 | -7 | -6 | -5 | -4 | -3 | -2 | -1 | 0 | 1 | 2 |
|---|---|---|---|---|---|---|---|---|---|---|---|---|---|---|---|---|---|
| 13984.1 | PcUBI | YDA-2 | 0.92 | 18.3 | 0.1 | 1.5 | 0.4 | 0.8 | 0.1 | 2.6 | 0.0 | 2.6 | 0.0 | 4.7 | 74.1 | 4.9 | 0.0 |
| 13984.2 | PcUBI | YDA-2 | 0.96 | 2.9 | 0.0 | 0.0 | 0.0 | 1.2 | 0.0 | 0.0 | 0.0 | 0.0 | 0.0 | 0.0 | 92.8 | 0.0 | 0.0 |
| 13984.3 | PcUBI | YDA-2 | 0.89 | 11.4 | 0.0 | 0.0 | 0.4 | 0.0 | 0.0 | 0.1 | 1.3 | 1.3 | 0.0 | 4.6 | 77.3 | 3.0 | 0.0 |
| 13984.4 | PcUBI | YDA-2 | 0.92 | 25.4 | 0.0 | 0.0 | 0.2 | 2.1 | 0.0 | 2.2 | 2.8 | 0.0 | 0.2 | 1.7 | 66.8 | 15.5 | 0.0 |
| 13984.5 | PcUBI | YDA-2 | 0.91 | 50.4 | 0.0 | 0.0 | 1.4 | 0.8 | 0.0 | 1.7 | 5.5 | 1.0 | 0.8 | 9.4 | 40.2 | 29.6 | 0.0 |
| 13984.6 | PcUBI | YDA-2 | 0.94 | 19.2 | 3.2 | 0.0 | 0.4 | 0.2 | 1.4 | 0.1 | 3.5 | 0.8 | 0.0 | 4.9 | 74.6 | 4.6 | 0.0 |
| 13984.7 | PcUBI | YDA-2 | 0.93 | 13.0 | 0.0 | 0.0 | 0.4 | 0.1 | 0.0 | 0.0 | 0.0 | 0.9 | 0.0 | 6.8 | 79.6 | 4.5 | 0.0 |
| 13983.8 | TMM | YDA-2 | 0.93 | 6.5 | 0.0 | 0.0 | 0.4 | 0.0 | 0.0 | 0.0 | 0.0 | 0.4 | 0.0 | 3.9 | 86.9 | 1.8 | 0.0 |
| 13983.9 | TMM | YDA-2 | 0.94 | 3.4 | 0.0 | 0.0 | 0.4 | 0.0 | 0.0 | 0.0 | 0.0 | 0.0 | 0.0 | 2.1 | 90.7 | 0.4 | 0.0 |
| 13983.10 | TMM | YDA-2 | 0.95 | 2.8 | 0.0 | 0.0 | 0.0 | 0.7 | 0.0 | 0.0 | 0.0 | 1.2 | 0.0 | 0.0 | 92.7 | 0.0 | 0.0 |
| 13983.11 | TMM | YDA-2 | 0.94 | 5.2 | 0.0 | 0.0 | 0.0 | 0.2 | 0.0 | 0.4 | 0.1 | 1.5 | 0.0 | 1.4 | 89.0 | 0.4 | 0.0 |
| 13983.12 | TMM | YDA-2 | 0.96 | 3.7 | 0.0 | 0.0 | 0.0 | 0.0 | 0.0 | 0.0 | 0.0 | 0.0 | 0.0 | 2.8 | 92.0 | 0.4 | 0.0 |
| 13983.13 | TMM | YDA-2 | 0.94 | 4.0 | 0.0 | 0.0 | 0.0 | 0.0 | 0.0 | 0.0 | 0.0 | 0.6 | 0.0 | 3.3 | 90.4 | 0.0 | 0.0 |
| 13983.14 | TMM | YDA-2 | 0.96 | 3.8 | 0.0 | 0.0 | 0.0 | 0.5 | 0.0 | 0.0 | 0.0 | 0.8 | 0.0 | 1.4 | 92.5 | 0.0 | 0.0 |
| 13983.15 | TMM | YDA-2 | 0.87 | 8.5 | 0.0 | 0.0 | 0.0 | 1.7 | 0.0 | 0.0 | 0.0 | 0.7 | 0.0 | 4.2 | 78.7 | 1.6 | 0.0 |
| NLS-GFP | | | 0.95 | 2.8 | 0.0 | 0.0 | 0.0 | 0.0 | 0.0 | 0.0 | 0.0 | 0.0 | 0.0 | 2.7 | 92.5 | 0.1 | 0.0 |
| NLS-GFP | | | 0.95 | 3.4 | 0.0 | 0.0 | 0.0 | 0.0 | 0.0 | 0.0 | 0.0 | 0.0 | 0.0 | 2.7 | 91.8 | 0.7 | 0.0 |

**Supplementary File 13 | Indel profile of seedlings targeting *YDA* with *pPcUbi*:Cas9 or *pTMM*:Cas9.**

TIDE analysis of the upper band (1273 bp) from the YDA genotyping (**Supplementary File 15 B**) for individual *pPcUBI*:Cas9-mCherry;YDA-1,YDA-2 and *pTMM*:Cas9-mCherry;YDA-1,YDA-2 T1 lines. Parameters TIDE analysis: **A**, Target: *YDA-1* (expected cut at 163 bp), Alignment window: 50-143, Decomposition window: 188-400, Indel size: 20. **B**, Target: *YDA-2* (expected cut at 239 bp), Alignment window: 100-189, Decomposition window: 254-400, Indel size: 10. Significant values (p<0.001) are highlighted in green.

#### Supplementary File 14

| T1 Line | Promoter | Target | GFP phenotype in LR | $R^2$ | TIDE Score (%) | -10 | -9 | -8 | -7 | -6 | -5 | -4 | -3 | -2 | -1 | 0 | 1 | 2 | 3 | 4 | 5 | 6 | 7 | 8 | 9 | 10 |
|---|---|---|---|---|---|---|---|---|---|---|---|---|---|---|---|---|---|---|---|---|---|---|---|---|---|---|
| 13977.4 | GATA23 | GFP-1 | KO | 0.99 | 96.9 | 0.0 | 0.2 | 0.0 | 0.0 | 0.0 | 0.0 | 0.0 | 1.9 | 0.0 | 12.5 | 1.8 | 82.4 | 0.0 | 0.0 | 0.0 | 0.0 | 0.0 | 0.0 | 0.0 | 0.0 | 0.0 |
| 13977.5 | GATA23 | GFP-1 | KO | 0.99 | 96.4 | 0.0 | 0.0 | 0.0 | 0.0 | 0.1 | 0.0 | 1.2 | 3.4 | 1.8 | 9.6 | 2.1 | 78.9 | 0.0 | 0.7 | 0.0 | 0.0 | 0.0 | 0.7 | 0.0 | 0.1 | 0.0 |
| 13977.32 | GATA23 | GFP-1 | KO | 0.96 | 96.3 | 0.0 | 0.2 | 0.0 | 0.0 | 0.0 | 0.0 | 0.0 | 0.0 | 0.0 | 17.3 | 0.0 | 76.5 | 0.0 | 0.0 | 0.0 | 0.0 | 0.0 | 0.0 | 0.0 | 0.0 | 2.3 |
| 13977.33 | GATA23 | GFP-1 | KO | 0.99 | 95.3 | 0.0 | 0.2 | 0.0 | 0.0 | 0.0 | 0.1 | 0.0 | 0.0 | 0.0 | 0.0 | 3.7 | 94.7 | 0.0 | 0.1 | 0.2 | 0.0 | 0.0 | 0.0 | 0.0 | 0.0 | 0.0 |
| 13977.30 | GATA23 | GFP-1 | KO | 0.98 | 93.7 | 0.0 | 0.1 | 0.2 | 0.0 | 0.0 | 0.1 | 0.0 | 0.0 | 0.0 | 2.4 | 4.6 | 90.8 | 0.0 | 0.0 | 0.0 | 0.0 | 0.0 | 0.0 | 0.0 | 0.0 | 0.1 |
| 13977.34 | GATA23 | GFP-1 | KO | 0.98 | 93.0 | 1.9 | 0.0 | 0.0 | 0.0 | 0.0 | 0.2 | 0.0 | 0.0 | 0.0 | 4.6 | 4.5 | 86.3 | 0.0 | 0.0 | 0.0 | 0.0 | 0.0 | 0.0 | 0.0 | 0.0 | 0.0 |
| 13977.6 | GATA23 | GFP-1 | Chimeric | 0.93 | 29.4 | 0.4 | 0.0 | 0.8 | 0.0 | 0.2 | 0.1 | 0.0 | 1.2 | 0.0 | 2.9 | 63.7 | 23.2 | 0.3 | 0.0 | 0.0 | 0.0 | 0.0 | 0.1 | 0.0 | 0.0 | 0.2 |
| 13977.7 | GATA23 | GFP-1 | WT | 0.99 | 3.2 | 0.0 | 0.0 | 0.0 | 0.0 | 0.2 | 0.0 | 0.0 | 0.2 | 0.0 | 0.0 | 96.0 | 1.1 | 0.3 | 0.3 | 0.2 | 0.2 | 0.0 | 0.2 | 0.0 | 0.5 | 0.0 |
| 13977.31 | GATA23 | GFP-1 | WT | 0.99 | 1.7 | 0.0 | 0.0 | 0.0 | 0.0 | 0.0 | 0.0 | 0.0 | 0.6 | 0.0 | 0.0 | 97.4 | 0.5 | 0.6 | 0.0 | 0.0 | 0.0 | 0.0 | 0.0 | 0.0 | 0.0 | 0.0 |

**Supplementary File 14 | Indel profile of individual *pGATA23*:Cas9-mCherry;GFP-1 T1 lines**

Parameters TIDE analysis: Target: *GFP-1* (expected cut at 179 bp), Alignment window: 100-169, Decomposition window: 194-400, Indel size: 10. Significant values ($p<0.001$) are highlighted in green.

#### Supplementary File 15

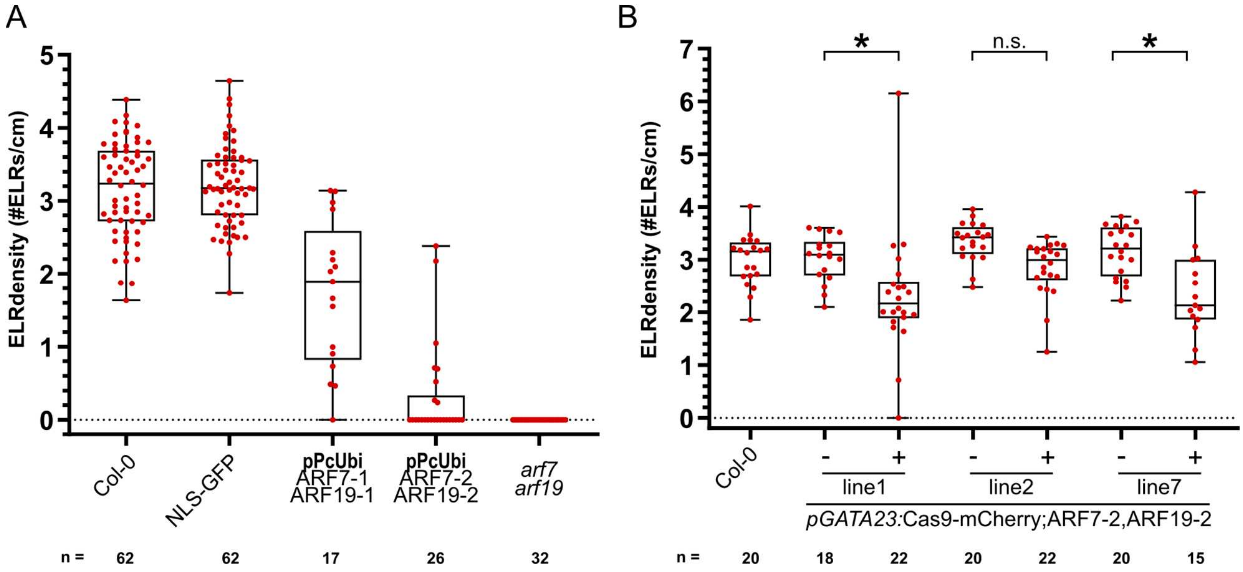

#### Supplementary File 15 | Targeting of *ARF7* and *ARF19*

**A**, Quantification of the emerged lateral root (ELR) density for Col-0, NLS-GFP, T1 seedlings of ubiquitous *pPcUBI*:Cas9 targeting *ARF7* and *ARF19*, and *arf7arf19* knockout line. **B**, Quantification of the emerged lateral root (ELR) density for Col-0 and FAST-positive (+) and –negative (-) T2 seedlings of *pGATA23*-CRISPR-TSKO lines targeting *ARF7* and *ARF19* simultaneously. ELR density was compared between FAST positive and negative seedlings within each line via Poisson regression analyses. n.s. indicates not significant with an $\alpha=0.05$. * indicates *p* values smaller than $2x10^{-3}$.

#### Supplementary File 16

### A

| Sample | Promoter | Target | mCherry Expression | $R^2$ | TIDE Score (%) | -22 | -17 | -10 | -9 | -8 | -7 | -6 | -5 | -4 | -3 | -2 | -1 | 0 | 1 | 2 |
|---|---|---|---|---|---|---|---|---|---|---|---|---|---|---|---|---|---|---|---|---|
| 13981.8 | PcUBI | ARF7-1 | Positive | 0.90 | 65.3 | 6.2 | 5.0 | 0.0 | 6.6 | 0.0 | 8.7 | 0.0 | 0.0 | 0.1 | 6.4 | 3.3 | 5.3 | 24.7 | 23.3 | 0.0 |
| 13981.9 | PcUBI | ARF7-1 | Positive | 0.99 | 24.8 | 0.8 | 0.0 | 0.0 | 0.0 | 0.0 | 0.0 | 6.1 | 0.1 | 0.4 | 1.2 | 0.3 | 5.1 | 73.9 | 10.7 | 0.0 |
| 13981.10 | PcUBI | ARF7-1 | Positive | 0.98 | 21.8 | 2.0 | 1.0 | 0.0 | 0.0 | 0.0 | 0.0 | 0.0 | 0.1 | 1.6 | 2.4 | 2.4 | 3.5 | 76.5 | 7.6 | 0.0 |
| 13981.11 | PcUBI | ARF7-1 | Positive | 0.92 | 63.8 | 4.7 | 0.0 | 0.0 | 0.0 | 0.0 | 0.0 | 0.0 | 2.6 | 0.5 | 2.6 | 11.9 | 1.8 | 28.6 | 38.2 | 1.4 |
| NLS-GFP | | | | 0.98 | 4.7 | 0.0 | 0.0 | 0.0 | 0.0 | 0.0 | 0.7 | 0.0 | 0.0 | 0.0 | 0.0 | 0.2 | 1.5 | 93.2 | 0.6 | 0.7 |

### B

| Sample | Promoter | Target | mCherry Expression | $R^2$ | TIDE Score (%) | -13 | -12 | -11 | -10 | -9 | -8 | -7 | -6 | -5 | -4 | -3 | -2 | -1 | 0 | 1 | 2 |
|---|---|---|---|---|---|---|---|---|---|---|---|---|---|---|---|---|---|---|---|---|---|
| 13981.8 | PcUBI | ARF19-1 | Positive | 0.99 | 9.6 | 1.3 | 0.1 | 0.0 | 0.1 | 0.1 | 0.0 | 0.5 | 0.0 | 0.0 | 0.0 | 0.0 | 0.0 | 1.6 | 89.0 | 5.3 | 0.0 |
| 13981.9 | PcUBI | ARF19-1 | Positive | 0.99 | 3.3 | 0.0 | 0.0 | 0.2 | 0.0 | 0.0 | 0.0 | 0.0 | 0.0 | 0.0 | 0.0 | 0.4 | 0.1 | 0.5 | 95.9 | 1.2 | 0.0 |
| 13981.10 | PcUBI | ARF19-1 | Positive | 0.99 | 3.1 | 0.0 | 0.0 | 0.1 | 0.0 | 0.0 | 0.0 | 0.0 | 0.0 | 0.1 | 0.0 | 0.4 | 0.0 | 0.5 | 96.2 | 1.6 | 0.0 |
| 13981.11 | PcUBI | ARF19-1 | Positive | 0.99 | 13.3 | 0.0 | 0.0 | 0.0 | 0.0 | 0.0 | 0.0 | 0.0 | 0.0 | 0.0 | 0.0 | 6.8 | 0.0 | 0.7 | 85.6 | 5.8 | 0.0 |
| NLS-GFP | | | | 0.99 | 1.4 | 0.0 | 0.0 | 0.0 | 0.0 | 0.0 | 0.0 | 0.0 | 0.0 | 0.0 | 0.0 | 0.0 | 0.0 | 0.0 | 97.9 | 0.0 | 0.0 |

### C

| Sample | Promoter | Target | mCherry Expression | $R^2$ | TIDE Score (%) | -22 | -21 | -20 | -10 | -9 | -8 | -7 | -6 | -5 | -4 | -3 | -2 | -1 | 0 | 1 | 2 |
|---|---|---|---|---|---|---|---|---|---|---|---|---|---|---|---|---|---|---|---|---|---|
| 13982.12 | PcUBI | ARF7-2 | Positive | 0.96 | 96.2 | 4.3 | 1.6 | 0.0 | 0.0 | 0.0 | 0.0 | 0.0 | 0.0 | 0.2 | 2.3 | 23.6 | 0.9 | 6.6 | 0.0 | 56.8 | 0.0 |
| 13982.13 | PcUBI | ARF7-2 | Positive | 0.95 | 95.0 | 2.5 | 0.6 | 0.0 | 2.1 | 0.0 | 1.8 | 0.0 | 0.0 | 7.3 | 1.1 | 0.0 | 24.7 | 5.9 | 0.0 | 49.2 | 0.0 |
| 13982.14 | PcUBI | ARF7-2 | Positive | 0.93 | 74.7 | 5.9 | 0.4 | 2.5 | 0.0 | 0.0 | 0.0 | 0.0 | 0.0 | 0.0 | 36.9 | 0.0 | 0.7 | 0.0 | 18.3 | 28.2 | 0.0 |
| 113982.5 | PcUBI | ARF7-2 | Negative | 0.99 | 0.5 | 0.0 | 0.0 | 0.0 | 0.0 | 0.0 | 0.0 | 0.0 | 0.0 | 0.0 | 0.0 | 0.0 | 0.0 | 0.0 | 98.8 | 0.0 | 0.0 |
| NLS-GFP | | | | 0.99 | 2.5 | 0.0 | 0.0 | 0.0 | 0.0 | 0.0 | 0.0 | 0.0 | 0.0 | 0.0 | 0.0 | 0.0 | 0.0 | 1.0 | 96.2 | 1.5 | 0.0 |

### D

| Sample | Promoter | Target | mCherry Expression | $R^2$ | TIDE Score (%) | -22 | -12 | -11 | -10 | -9 | -8 | -7 | -6 | -5 | -4 | -3 | -2 | -1 | 0 | 1 | 2 | 8 |
|---|---|---|---|---|---|---|---|---|---|---|---|---|---|---|---|---|---|---|---|---|---|---|
| 13982.12 | PcUBI | ARF19-2 | Positive | 0.99 | 98.5 | 3.9 | 0.0 | 0.0 | 0.1 | 0.0 | 1.0 | 0.0 | 0.8 | 0.3 | 0.5 | 0.0 | 0.0 | 0.7 | 0.7 | 87.6 | 0.0 | 2.0 |
| 13982.13 | PcUBI | ARF19-2 | Positive | 0.95 | 94.5 | 1.4 | 2.0 | 0.1 | 0.0 | 0.0 | 0.0 | 0.4 | 0.7 | 0.9 | 0.1 | 2.3 | 0.9 | 0.0 | 0.0 | 73.3 | 0.0 | 0.3 |
| 13982.14 | PcUBI | ARF19-2 | Positive | 0.99 | 98.9 | 0.0 | 0.0 | 0.0 | 0.0 | 0.0 | 0.0 | 0.0 | 23.4 | 0.0 | 0.0 | 0.1 | 0.0 | 0.0 | 0.0 | 73.7 | 0.0 | 0.0 |
| 13982.15 | PcUBI | ARF19-2 | Negative | 0.99 | 1.6 | 0.0 | 0.0 | 0.0 | 0.0 | 0.0 | 0.0 | 0.0 | 0.3 | 0.0 | 0.0 | 0.0 | 0.0 | 0.0 | 97.7 | 0.1 | 0.0 | 0.0 |
| NLS-GFP | | | | 0.99 | 1.2 | 0.0 | 0.0 | 0.0 | 0.0 | 0.0 | 0.0 | 0.0 | 0.1 | 0.0 | 0.0 | 0.0 | 0.0 | 0.0 | 98.1 | 0.0 | 0.0 | 0.0 |

**Supplementary File 16 | Genotyping of individual T1 lines of *pPcUBI*:Cas9-mCherry;ARF7-1,ARF19-1 and *pPcUBI*:Cas-mCherry;ARF7-2,ARF19-2**

Parameters TIDE analysis: **A**, Target: *ARF7-1* (expected cut at 213 bp), Alignment window: 100-188, Decomposition window: 243-550, Indel size: 25. **B**, Target: *ARF19-1* (expected cut at 195 bp), Alignment window: 100-170, Decomposition window: 248-500, Indel size: 25. **C**, Target: *ARF7-2* (expected cut at 218 bp), Alignment window: 100-193, Decomposition window: 248-670, Indel size: 25. **D**, Target: *ARF19-2* (expected cut at 169 bp), Alignment window: 100-144, Decomposition window: 199-500, Indel size: 25. Significant values (p<0.001) are highlighted in green.

### Supplementary File 17

**A**

| Sample | Promoter | Target | FAST Expression | $R^2$ | TIDE Score (%) | -22 | -21 | -20 | -16 | -10 | -9 | -8 | -7 | -6 | -5 | -4 | -3 | -2 | -1 | 0 | 1 | 2 | 3 | 4 | 5 | 17 |
|---|---|---|---|---|---|---|---|---|---|---|---|---|---|---|---|---|---|---|---|---|---|---|---|---|---|---|
| 13982.1.1 | GATA23 | ARF7-2 | Positive | 0.92 | 89.5 | 12.9 | 0.0 | 6.8 | 0.0 | 0.4 | 0.8 | 0.0 | 0.0 | 0.2 | 3.6 | 0.0 | 1.1 | 0.0 | 0.7 | 2.3 | 50.2 | 5.5 | 0.6 | 0.0 | 0.0 | 0.0 |
| 13982.1.4 | GATA23 | ARF7-2 | Positive | 0.92 | 46.3 | 11.7 | 0.0 | 0.0 | 0.0 | 0.0 | 0.0 | 0.5 | 0.0 | 0.0 | 0.0 | 0.3 | 0.0 | 0.0 | 0.0 | 46.0 | 32.5 | 0.0 | 0.0 | 0.0 | 0.2 | 0.0 |
| 13982.1.6 | GATA23 | ARF7-2 | Positive | 0.96 | 88.0 | 0.0 | 0.0 | 0.0 | 0.0 | 0.0 | 1.9 | 0.0 | 4.5 | 0.0 | 3.7 | 0.0 | 0.0 | 0.0 | 0.0 | 7.8 | 67.9 | 3.6 | 0.0 | 4.2 | 1.2 | 0.0 |
| 13982.1.9 | GATA23 | ARF7-2 | Negative | 0.99 | 0.0 | 0.0 | 0.0 | 0.0 | 0.0 | 0.0 | 0.0 | 0.0 | 0.0 | 0.0 | 0.0 | 0.0 | 0.0 | 0.0 | 0.0 | 98.9 | 0.0 | 0.0 | 0.0 | 0.0 | 0.0 | 0.0 |
| 13982.1.10 | GATA23 | ARF7-2 | Negative | 1.00 | 0.5 | 0.0 | 0.0 | 0.0 | 0.0 | 0.0 | 0.0 | 0.0 | 0.0 | 0.0 | 0.0 | 0.0 | 0.0 | 0.0 | 0.0 | 99.1 | 0.1 | 0.0 | 0.0 | 0.0 | 0.0 | 0.0 |
| 13982.2.15 | GATA23 | ARF7-2 | Positive | 0.95 | 70.9 | 2.9 | 0.0 | 0.0 | 3.9 | 0.0 | 0.0 | 0.0 | 0.0 | 0.0 | 0.0 | 0.2 | 15.4 | 0.0 | 0.0 | 23.7 | 47.4 | 0.0 | 0.0 | 0.0 | 0.0 | 0.0 |
| 13982.2.16 | GATA23 | ARF7-2 | Positive | 0.99 | 98.7 | 0.0 | 0.0 | 0.1 | 0.0 | 0.0 | 0.0 | 0.0 | 0.0 | 0.3 | 0.0 | 0.0 | 0.0 | 0.0 | 3.8 | 0.0 | 93.3 | 0.0 | 0.0 | 0.0 | 0.9 | 0.0 |
| 13982.2.17 | GATA23 | ARF7-2 | Positive | 0.90 | 85.6 | 11.7 | 0.0 | 6.0 | 0.0 | 0.0 | 0.0 | 0.0 | 0.0 | 0.0 | 0.0 | 0.0 | 0.0 | 0.0 | 4.2 | 4.3 | 54.3 | 0.4 | 0.0 | 0.1 | 0.0 | 4.8 |
| 13982.2.21 | GATA23 | ARF7-2 | Negative | 0.99 | 0.0 | 0.0 | 0.0 | 0.0 | 0.0 | 0.0 | 0.0 | 0.0 | 0.0 | 0.0 | 0.0 | 0.0 | 0.0 | 0.0 | 0.0 | 98.9 | 0.0 | 0.0 | 0.0 | 0.0 | 0.0 | 0.0 |
| 13982.2.22 | GATA23 | ARF7-2 | Negative | 0.99 | 2.1 | 0.0 | 0.2 | 0.0 | 0.2 | 0.0 | 0.0 | 0.0 | 0.0 | 0.0 | 0.0 | 0.0 | 0.0 | 0.0 | 0.5 | 97.3 | 0.0 | 0.0 | 0.0 | 0.0 | 0.5 | 0.0 |
| 13982.7.28 | GATA23 | ARF7-2 | Positive | 0.91 | 88.1 | 24.6 | 0.0 | 0.0 | 0.0 | 0.0 | 0.0 | 0.0 | 0.7 | 0.0 | 0.0 | 0.0 | 0.0 | 0.0 | 36.6 | 2.8 | 23.5 | 0.0 | 0.0 | 0.0 | 0.0 | 0.0 |
| 13982.7.29 | GATA23 | ARF7-2 | Positive | 0.95 | 94.6 | 4.7 | 0.0 | 2.2 | 0.0 | 1.1 | 0.0 | 0.5 | 0.0 | 0.0 | 0.0 | 7.6 | 1.3 | 2.6 | 10.4 | 0.0 | 61.8 | 0.0 | 0.0 | 0.0 | 0.0 | 0.0 |
| 13982.7.30 | GATA23 | ARF7-2 | Positive | 0.93 | 83.6 | 9.7 | 0.0 | 1.2 | 0.6 | 0.0 | 0.0 | 0.0 | 0.0 | 7.1 | 0.0 | 3.8 | 0.0 | 0.0 | 0.0 | 9.6 | 60.4 | 0.0 | 0.0 | 0.0 | 0.0 | 0.5 |
| 13982.7.32 | GATA23 | ARF7-2 | Positive | 0.90 | 84.2 | 22.2 | 0.0 | 0.2 | 0.0 | 0.0 | 4.5 | 0.0 | 0.0 | 0.0 | 0.0 | 0.0 | 0.0 | 0.0 | 1.6 | 6.3 | 48.7 | 0.0 | 0.0 | 0.0 | 0.6 | 0.5 |
| 13982.7.33 | GATA23 | ARF7-2 | Negative | 1.00 | 1.3 | 0.0 | 0.0 | 0.0 | 0.0 | 0.0 | 0.0 | 0.4 | 0.0 | 0.0 | 0.0 | 0.0 | 0.0 | 0.0 | 0.0 | 98.3 | 0.0 | 0.0 | 0.0 | 0.0 | 0.0 | 0.0 |

**B**

| Sample | Promoter | Target | FAST Expression | $R^2$ | TIDE Score (%) | -36 | -34 | -29 | -26 | -22 | -21 | -12 | -9 | -8 | -7 | -6 | -5 | -4 | -3 | -2 | -1 |
|---|---|---|---|---|---|---|---|---|---|---|---|---|---|---|---|---|---|---|---|---|---|
| 13982.1.1 | GATA23 | ARF19-2 | Positive | 0.96 | 96.2 | 0.0 | 7.5 | 0 | 0.0 | 0 | 0.0 | 0.0 | 0.0 | 0.0 | 0.0 | 0.0 | 0 | 0.0 | 5.8 | 4.5 | 5.7 |
| 13982.1.4 | GATA23 | ARF19-2 | Positive | 0.94 | 66.7 | 0.5 | 1.4 | 0.0 | 0.0 | 1 | 0.0 | 0.0 | 0.0 | 0.0 | 0.0 | 3.3 | 10.6 | 0.0 | 4.3 | 0.0 | 5.0 |
| 13982.1.6 | GATA23 | ARF19-2 | Positive | 0.94 | 93.9 | 0.0 | 0.0 | 7.8 | 2.8 | 0.0 | 0.0 | 0.0 | 0.0 | 0.0 | 2.4 | 15.3 | 1.9 | 0.0 | 0.0 | 0.0 | 4.0 |
| 13982.1.9 | GATA23 | ARF19-2 | Negative | 1.00 | 0.1 | 0.0 | 0.0 | 0.0 | 0.0 | 0.0 | 0.0 | 0.0 | 0.0 | 0.0 | 0.0 | 0.0 | 0.0 | 0.0 | 0.0 | 0.0 | 0.0 |
| 13982.1.10 | GATA23 | ARF19-2 | Negative | 1.00 | 0.3 | 0.0 | 0.0 | 0.0 | 0.0 | 0.2 | 0.0 | 0.0 | 0.0 | 0.0 | 0.0 | 0.0 | 0.0 | 0.0 | 0.0 | 0.0 | 0.0 |
| 13982.2.15 | GATA23 | ARF19-2 | Positive | 0.96 | 96.4 | 0.0 | 0.0 | 0.0 | 0.0 | 0.0 | 0.0 | 0.0 | 0.0 | 0.0 | 0.0 | 5.6 | 0.0 | 0.0 | 16.7 | 7.4 | 0.0 |
| 13982.2.16 | GATA23 | ARF19-2 | Positive | 0.94 | 94.0 | 0.2 | 0.0 | 0.0 | 0.0 | 0.0 | 0.0 | 0.0 | 0.0 | 0.0 | 0.0 | 28.8 | 0.0 | 0.0 | 0.0 | 0.0 | 0.0 |
| 13982.2.17 | GATA23 | ARF19-2 | Positive | 0.97 | 97.5 | 8.3 | 0.0 | 0.0 | 3.1 | 5.8 | 0.0 | 0.0 | 0.0 | 0.0 | 0.0 | 0.0 | 0.6 | 0.0 | 5.7 | 0.0 | 0.0 |
| 13982.2.21 | GATA23 | ARF19-2 | Negative | 1.00 | 0.0 | 0.0 | 0.0 | 0.0 | 0.0 | 0.0 | 0.0 | 0.0 | 0.0 | 0.0 | 0.0 | 0.0 | 0.0 | 0.0 | 0.0 | 0.0 | 0.0 |
| 13982.2.22 | GATA23 | ARF19-2 | Negative | 1.00 | 0.9 | 0.0 | 0.0 | 0.0 | 0.0 | 0.0 | 0.0 | 0.0 | 0.4 | 0.0 | 0.0 | 0.0 | 0.0 | 0.3 | 0.0 | 0.2 | 0.0 |
| 13982.7.28 | GATA23 | ARF19-2 | Positive | 0.93 | 93.2 | 0.0 | 0.0 | 0.0 | 0.0 | 0.0 | 0.0 | 0.0 | 0.0 | 0.0 | 10.5 | 12.1 | 0.0 | 2.2 | 45.2 | 0.0 | 0.0 |
| 13982.7.29 | GATA23 | ARF19-2 | Positive | 0.99 | 98.6 | 0.0 | 0.0 | 0.0 | 0.0 | 0.0 | 3.1 | 0.0 | 2.7 | 0.0 | 0.0 | 0.5 | 0.0 | 0.0 | 9.0 | 4.3 | 0.0 |
| 13982.7.30 | GATA23 | ARF19-2 | Positive | 0.93 | 92.8 | 10.3 | 0.0 | 0.0 | 0.0 | 0.0 | 0.0 | 16.9 | 0.0 | 0.0 | 0.0 | 12.2 | 0.0 | 1.2 | 1.2 | 0.0 | 0.0 |
| 13982.7.32 | GATA23 | ARF19-2 | Positive | 1.00 | 99.5 | 0.0 | 0.0 | 0.0 | 0.0 | 3.4 | 0.0 | 0.0 | 0.0 | 0.0 | 2.2 | 0.1 | 0.0 | 0.4 | 1.0 | 0.0 | 3.1 |
| 13982.7.33 | GATA23 | ARF19-2 | Negative | 1.00 | 0.0 | 0.0 | 0.0 | 0.0 | 0.0 | 0.0 | 0.0 | 0.0 | 0.0 | 0.0 | 0.0 | 0.0 | 0.0 | 0.0 | 0.0 | 0.0 | 0.0 |

| Sample | 0 | 1 | 2 | 3 | 7 | 8 | 9 | 10 | 11 | 12 | 13 | 14 | 15 | 25 |
|---|---|---|---|---|---|---|---|---|---|---|---|---|---|---|
| 13982.1.1 | 0.0 | 62.6 | 5.3 | 0.0 | 0.0 | 0.0 | 0.0 | 0.0 | 0.0 | 0.0 | 0.0 | 0.0 | 0.8 | 1.3 |
| 13982.1.4 | 27.7 | 39.7 | 0.0 | 0.0 | 0.0 | 0.0 | 0.0 | 0.0 | 0.0 | 0.0 | 0.0 | 0.0 | 0.0 | 0.0 |
| 13982.1.6 | 0 | 46.5 | 0 | 0.0 | 7.8 | 0.0 | 0.0 | 0.0 | 0.0 | 0.0 | 0.0 | 0.0 | 4.6 | 0.0 |
| 13982.1.9 | 99.6 | 0.0 | 0.0 | 0.0 | 0.0 | 0.0 | 0.0 | 0.0 | 0.0 | 0.0 | 0.0 | 0.0 | 0.0 | 0.0 |
| 13982.1.10 | 99.4 | 0.0 | 0.0 | 0.0 | 0.0 | 0.0 | 0.0 | 0.0 | 0.0 | 0.0 | 0.0 | 0.0 | 0.0 | 0.0 |
| 13982.2.15 | 0.0 | 56.8 | 0.0 | 0.0 | 0.0 | 9.9 | 0.0 | 0.0 | 0.0 | 0.0 | 0.0 | 0.0 | 0.0 | 0.0 |
| 13982.2.16 | 0.0 | 57.8 | 0.0 | 0.0 | 0.0 | 0.0 | 0.0 | 0.3 | 0.0 | 0.0 | 0.7 | 0.2 | 0.0 | 0.0 |
| 13982.2.17 | 0.0 | 64.4 | 2.7 | 2.4 | 0.0 | 0.0 | 0.6 | 2.5 | 0.0 | 1.4 | 0.0 | 0.0 | 0.0 | 0.0 |
| 13982.2.21 | 99.6 | 0.0 | 0.0 | 0.0 | 0.0 | 0.0 | 0.0 | 0.0 | 0.0 | 0.0 | 0.0 | 0.0 | 0.0 | 0.0 |
| 13982.2.22 | 98.8 | 0.0 | 0.0 | 0.0 | 0.0 | 0.0 | 0.0 | 0.0 | 0.0 | 0.0 | 0.0 | 0.0 | 0.0 | 0.0 |
| 13982.7.28 | 0.0 | 21.5 | 1.7 | 0.0 | 0.0 | 0.0 | 0.0 | 0.0 | 0.0 | 0.0 | 0.0 | 0.0 | 0.0 | 0.0 |
| 13982.7.29 | 0.0 | 78.5 | 0.0 | 0.0 | 0.0 | 0.0 | 0.0 | 0.0 | 0.0 | 0.0 | 0.0 | 0.0 | 0.0 | 0.0 |
| 13982.7.30 | 0.0 | 44.9 | 2.3 | 0.0 | 0.0 | 0.0 | 0.0 | 1.8 | 0.0 | 0.0 | 1.6 | 0.0 | 0.0 | 0.0 |
| 13982.7.32 | 0.0 | 88.8 | 0.0 | 0.0 | 0.0 | 0.0 | 0.6 | 0.0 | 0.0 | 0.0 | 0.0 | 0.0 | 0.0 | 0.0 |
| 13982.7.33 | 99.7 | 0.0 | 0.0 | 0.0 | 0.0 | 0.0 | 0.0 | 0.0 | 0.0 | 0.0 | 0.0 | 0.0 | 0.0 | 0.0 |

**Supplementary File 17 | Genotyping of individual T2 lines of *pGATA23*:Cas9-mCherry;ARF7-2,ARF19-2**

Parameters TIDE analysis: **A**, Target: *ARF7-2* (expected cut at 216 bp), Alignment window: 100-191, Decomposition window: 246-330, Indel size: 25. **B**, Target: *ARF19-2* (expected cut at 171 bp), Alignment window: 90-131, Decomposition window: 216-600, Indel size: 40. Significant values ($p<0.001$) are highlighted in green.

#### Supplementary File 18

**A**

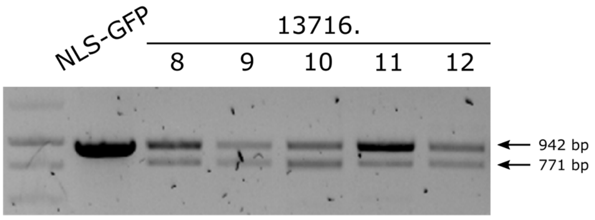

**B**

| Sample | Promoter | Target | $R^2$ | TIDE Score (%) | -10 | -9 | -8 | -7 | -6 | -5 | -4 | -3 | -2 | -1 | 0 | 1 | 2 | 3 | 4 | 5 | 6 | 7 | 8 | 9 | 10 |
|---|---|---|---|---|---|---|---|---|---|---|---|---|---|---|---|---|---|---|---|---|---|---|---|---|---|
| 13716.8 | PcUBI | CDKA;1-1 | 0.96 | 93.6 | 7.4 | 0.0 | 0.0 | 0.0 | 6.4 | 2.0 | 0.2 | 4.2 | 0.0 | 0.0 | 2.8 | 64.1 | 0.0 | 0.1 | 0.0 | 0.0 | 0.3 | 0.0 | 0.0 | 0.6 | 0.4 |
| 13716.9 | PcUBI | CDKA;1-1 | 0.99 | 98.6 | 0.0 | 0.0 | 0.0 | 0.0 | 0.0 | 0.0 | 0.0 | 16.9 | 0.0 | 0.0 | 0.0 | 79.8 | 0.0 | 0.3 | 0.0 | 0.0 | 0.0 | 0.7 | 0.0 | 0.0 | 0.4 |
| 13716.10 | PcUBI | CDKA;1-1 | 0.96 | 69.9 | 0.0 | 0.3 | 0.2 | 0.0 | 0.5 | 0.0 | 0.0 | 0.0 | 0.0 | 0.0 | 25.9 | 59.9 | 0.0 | 0.0 | 0.2 | 0.0 | 0.0 | 0.0 | 0.0 | 0.0 | 0.0 |
| 13716.11 | PcUBI | CDKA;1-1 | 0.99 | 77.5 | 0.4 | 0.0 | 0.0 | 1.0 | 0.0 | 0.1 | 0.0 | 0.0 | 0.0 | 0.6 | 21.5 | 72.4 | 0.0 | 0.0 | 0.0 | 0.0 | 0.0 | 0.0 | 0.0 | 0.0 | 0.1 |
| 13716.12 | PcUBI | CDKA;1-1 | 0.98 | 48.4 | 15.4 | 0.0 | 0.0 | 0.0 | 0.0 | 0.0 | 0.0 | 0.0 | 0.0 | 0.0 | 49.6 | 31.5 | 0.0 | 0.0 | 0.0 | 0.0 | 0.0 | 0.0 | 0.0 | 0.0 | 0.0 |

**C**

| Sample | Promoter | Target | $R^2$ | TIDE Score (%) | -23 | -22 | -21 | -20 | -10 | -9 | -8 | -7 | -6 | -5 | -4 | -3 | -2 | -1 | 0 | 1 | 2 | 3 | 4 | 5 | 6 | 7 | 8 | 9 | 10 |
|---|---|---|---|---|---|---|---|---|---|---|---|---|---|---|---|---|---|---|---|---|---|---|---|---|---|---|---|---|---|
| 13716.8 | PcUBI | CDKA;1-2 | 0.87 | 79.3 | 0.0 | 0.1 | 2.7 | 0.8 | 0.2 | 0.2 | 2.3 | 0.0 | 3.4 | 1.1 | 0.0 | 8.7 | 9.4 | 29.0 | 7.8 | 15.2 | 0.0 | 0.0 | 0.0 | 0.0 | 0.0 | 0.0 | 0.0 | 0.0 | 0.4 |
| 13716.9 | PcUBI | CDKA;1-2 | - | - | | | | | | | | | | | | | | | | | | | | | | | | | |
| 13716.10 | PcUBI | CDKA;1-2 | 0.92 | 34.1 | 0.0 | 5.6 | 5.9 | 0.0 | 0.0 | 0.0 | 0.3 | 0.0 | 0.0 | 0.0 | 0.0 | 4.6 | 0.0 | 5.3 | 58.1 | 5.3 | 0.0 | 3.1 | 0.0 | 0.0 | 0.0 | 0.0 | 0.0 | 0.1 | 0.0 |
| 13716.11 | PcUBI | CDKA;1-2 | 0.98 | 33.7 | 0.0 | 0.0 | 0.0 | 0.0 | 0.0 | 0.0 | 3.8 | 0.0 | 0.0 | 0.0 | 0.0 | 7.4 | 0.0 | 14.1 | 64.0 | 7.5 | 0.0 | 0.0 | 0.0 | 0.0 | 0.0 | 0.0 | 0.0 | 0.0 | 0.0 |
| 13716.12 | PcUBI | CDKA;1-2 | 0.99 | 16.0 | 0.0 | 0.0 | 0.0 | 0.0 | 7.1 | 0.0 | 0.0 | 0.0 | 0.0 | 0.0 | 0.0 | 0.0 | 0.0 | 4.4 | 82.8 | 1.2 | 0.0 | 0.0 | 0.0 | 0.0 | 0.0 | 0.0 | 0.0 | 0.0 | 0.0 |

**Supplementary File 18 | Genotyping and indel profile of *pPcUbi*:Cas9-mCherry;CDKA;1-1,CDKA;1-2**

**A**, Genotyping results for individual *pPcUBI*:Cas-mCherry;CDKA;1-1,CDKA;1-2 T1 lines. Lower band in gel is indicative of large deletion in *CDKA;1*. TIDE analysis was done on the upper band. Parameters TIDE analysis: **B**, Target: *CDKA;1-1* (expected cut at 421 bp), Alignment window: 100-391, Decomposition window: 456-560, Indel size: 30. **C**, Target: *CDKA;1-2* (expected cut at 293 bp), Alignment window: 100-263, Decomposition window: 328-432, Indel size: 30. Significant values (p<0.001) are highlighted in green.

#### Supplementary File 19

**A**

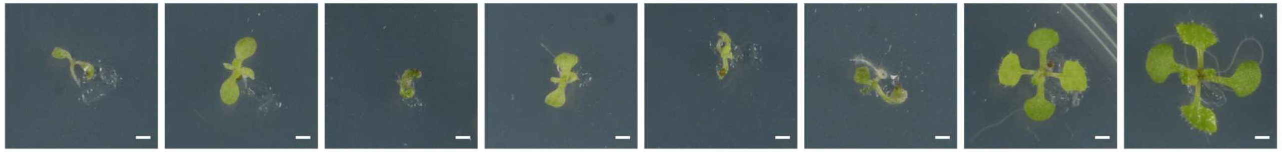

| | 14495.7 | 14495.8 | 14495.2 | 14495.15 | 14495.4 | 14495.17 | 14495.1 | NLS-GFP |
|---|---|---|---|---|---|---|---|---|
| CDKA;1-1 | 99.4 % | 98.9 % | 98.0 % | 97.3 % | 97.0 % | 94.7 % | 0.7 % | |
| CDKB1;1-1 | 97.5 % | 59.8 % | 98.0 % | 94.3 % | 95.5 % | 98.8 % | 0.0 % | |
| CDKB1;2-1 | 98.3 % | 66.1 % | 93.7 % | 61.2 % | 98.6 % | 92.0 % | 0.9 % | |

**B**

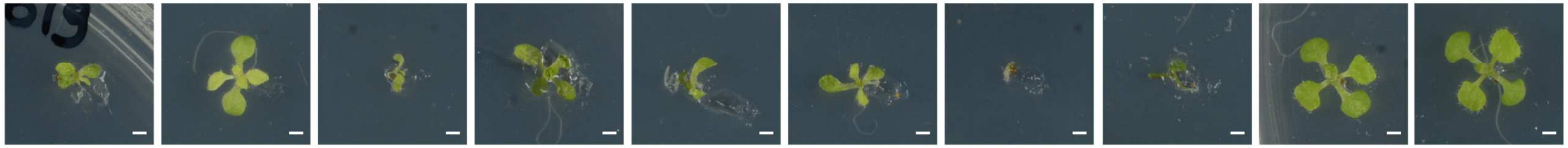

| | 14496.1 | 14496.17 | 14496.14 | 14496.6 | 14496.12 | 14496.7 | 14496.3 | 14496.18 | 14496.20 | NLS-GFP |
|---|---|---|---|---|---|---|---|---|---|---|
| CDKA;1-1 | 96.5 % | 81.6 % | 77.2 % | 71.5 % | 71.0 % | 69.4 % | 23.4 % | 1.0 % | 0.9 % | |
| CDKB1;1-2 | 65.0 % | 24.1 % | 6.1 % | 24.1 % | 68.1 % | 50.3 % | 42.1 % | 5.5 % | 4.3 % | |
| CDKB1;2-2 | 60.2 % | 22.1 % | 84.6 % | 25.4 % | 64.1 % | 47.4 % | 36.1 % | 1.2 % | 1.1 % | |

**Supplementary File 19 | Ubiquitous knockout of CDKA;1, CDKB1;1 and CDKB1;2**

Pictures of the individual T1 lines of *pPcUBI*:Cas9-mCherry;CDKA;1-1,CDKB1-1 (A) and *pPcUBI*:Cas9-mCherry;CDKA;1-1,CDKB1-2 (B) and the TIDE scores for the corresponding target sites (see **Supplemental File 21** for the genotyping). Scale bars represent 1 mm.

### Supplementary File 20

## A

| T1 Line | Promoter | Target | $R^2$ | TIDE Score (%) | -28 | -12 | -11 | -10 | -9 | -8 | -7 | -6 | -5 | -4 | -3 | -2 | -1 | 0 | 1 | 2 | 3 | 4 | 5 |
|---|---|---|---|---|---|---|---|---|---|---|---|---|---|---|---|---|---|---|---|---|---|---|---|
| 14495.7 | PcUBI | CDKA;1-1 | 0.99 | 99.4 | 0.0 | 0.0 | 0.0 | 0.0 | 0.0 | 0.0 | 0.0 | 0.0 | 0.0 | 0.0 | 0.0 | 0.0 | 0.0 | 0.0 | 98.5 | 0.0 | 0.7 | 0.1 | 0.0 |
| 14495.8 | PcUBI | CDKA;1-1 | 0.99 | 98.9 | 0.0 | 0.0 | 0.0 | 0.0 | 0.0 | 0.0 | 0.0 | 0.0 | 0.0 | 0.0 | 14.6 | 0.0 | 0.0 | 0.3 | 84.3 | 0.0 | 0.0 | 0.0 | 0.0 |
| 14495.2 | PcUBI | CDKA;1-1 | 0.98 | 98.0 | 0.0 | 0.0 | 0.0 | 0.0 | 0.0 | 30.7 | 0.0 | 0.0 | 0.0 | 0.0 | 0.0 | 0.0 | 0.0 | 0.0 | 67.4 | 0.0 | 0.0 | 0.0 | 0.0 |
| 14495.15 | PcUBI | CDKA;1-1 | 0.97 | 97.3 | 0.0 | 0.0 | 0.0 | 47.0 | 0.0 | 0.0 | 0.0 | 0.0 | 0.0 | 0.0 | 0.0 | 0.0 | 0.0 | 0.0 | 50.0 | 0.0 | 0.4 | 0.0 | 0.0 |
| 14495.4 | PcUBI | CDKA;1-1 | 0.97 | 97.0 | 8.0 | 0.0 | 3.0 | 0.0 | 0.0 | 0.0 | 0.0 | 0.0 | 0.0 | 0.0 | 34.5 | 0.0 | 0.0 | 0.0 | 50.2 | 0.0 | 1.3 | 0.0 | 0.0 |
| 14495.17 | PcUBI | CDKA;1-1 | 0.95 | 94.7 | 0.0 | 0.0 | 0.0 | 2.0 | 0.0 | 0.0 | 0.0 | 2.3 | 0.0 | 0.0 | 27.9 | 0.0 | 2.2 | 0.0 | 56.5 | 0.0 | 0.0 | 0.0 | 0.0 |
| 14495.1 | PcUBI | CDKA;1-1 | 1.00 | 0.7 | 0.0 | 0.0 | 0.0 | 0.1 | 0.0 | 0.0 | 0.0 | 0.0 | 0.0 | 0.0 | 0.0 | 0.0 | 0.0 | 98.8 | 0.3 | 0.0 | 0.0 | 0.0 | 0.0 |
| NLS-GFP | | CDKA;1-1 | 0.98 | 0.3 | 0.0 | 0.0 | 0.0 | 0.0 | 0.0 | 0.0 | 0.0 | 0.0 | 0.0 | 0.0 | 0.0 | 0.0 | 0.0 | 97.2 | 0.0 | 0.0 | 0.0 | 0.0 | 0.0 |

## B

| T1 Line | Promoter | Target | $R^2$ | TIDE Score (%) | -28 | -27 | -16 | -10 | -9 | -8 | -7 | -6 | -5 | -4 | -3 | -2 | -1 | 0 | 1 | 2 | 3 | 37 |
|---|---|---|---|---|---|---|---|---|---|---|---|---|---|---|---|---|---|---|---|---|---|---|
| 14495.7 | PcUBI | CDKB1;1-1 | 0.98 | 97.5 | 0.0 | 0.0 | 0.0 | 0.0 | 0.0 | 0.0 | 0.0 | 0.0 | 0.0 | 0.0 | 0.0 | 0.0 | 0.0 | 0.9 | 92.7 | 3.4 | 0.0 | 0.0 |
| 14495.8 | PcUBI | CDKB1;1-1 | 0.99 | 59.8 | 0.0 | 0.0 | 0.0 | 0.0 | 0.0 | 0.0 | 0.0 | 0.0 | 0.0 | 0.0 | 0.0 | 0.0 | 0.0 | 38.7 | 59.8 | 0.0 | 0.0 | 0.0 |
| 14495.2 | PcUBI | CDKB1;1-1 | 0.98 | 98.0 | 0.0 | 0.0 | 0.0 | 0.0 | 0.0 | 0.0 | 0.0 | 0.0 | 0.0 | 0.0 | 0.0 | 0.0 | 0.0 | 0.0 | 70.4 | 0.0 | 0.0 | 27.5 |
| 14495.15 | PcUBI | CDKB1;1-1 | 0.97 | 94.3 | 0.0 | 0.0 | 0.0 | 0.0 | 0.0 | 0.0 | 0.0 | 0.0 | 1.4 | 1.7 | 0.0 | 0.0 | 0.0 | 2.8 | 85.1 | 5.4 | 0.0 | 0.0 |
| 14495.4 | PcUBI | CDKB1;1-1 | 0.99 | 95.5 | 0.0 | 0.0 | 3.4 | 3.7 | 0.0 | 0.0 | 0.0 | 0.0 | 0.0 | 0.0 | 4.3 | 0.0 | 0.0 | 3.8 | 84.2 | 0.0 | 0.0 | 0.0 |
| 14495.17 | PcUBI | CDKB1;1-1 | 0.99 | 98.8 | 4.2 | 2.3 | 1.8 | 1.1 | 2.2 | 0.0 | 0.0 | 0.0 | 0.6 | 0.0 | 0.0 | 0.9 | 2.9 | 0.0 | 79.6 | 0.0 | 0.0 | 0.0 |
| 14495.1 | PcUBI | CDKB1;1-1 | 0.99 | 0.0 | 0.0 | 0.0 | 0.0 | 0.0 | 0.0 | 0.0 | 0.0 | 0.0 | 0.0 | 0.0 | 0.0 | 0.0 | 0.0 | 99.4 | 0.0 | 0.0 | 0.0 | 0.0 |
| NLS-GFP | | CDKB1;1-1 | 0.99 | 0.8 | 0.0 | 0.0 | 0.0 | 0.0 | 0.0 | 0.0 | 0.0 | 0.0 | 0.0 | 0.0 | 0.0 | 0.0 | 0.0 | 98.4 | 0.0 | 0.0 | 0.0 | 0.0 |

## C

| T1 Line | Promoter | Target | $R^2$ | TIDE Score (%) | -28 | -22 | -10 | -9 | -8 | -7 | -6 | -5 | -4 | -3 | -2 | -1 | 0 | 1 | 2 | 3 | 4 | 5 | 6 | 7 |
|---|---|---|---|---|---|---|---|---|---|---|---|---|---|---|---|---|---|---|---|---|---|---|---|---|
| 14495.7 | PcUBI | CDKB1;2-1 | 0.99 | 98.3 | 0.0 | 6.2 | 0.0 | 0.0 | 0.0 | 0.0 | 0.0 | 0.0 | 0.0 | 0.0 | 19.3 | 0.1 | 0.7 | 72.6 | 0.0 | 0.0 | 0.0 | 0.0 | 0.0 | 0.0 |
| 14495.8 | PcUBI | CDKB1;2-1 | 0.99 | 66.1 | 0.0 | 0.0 | 0.0 | 0.0 | 0.0 | 0.0 | 0.0 | 0.0 | 1.6 | 0.0 | 0.0 | 1.0 | 32.5 | 62.0 | 0.0 | 0.5 | 0.0 | 0.0 | 0.0 | 0.7 |
| 14495.2 | PcUBI | CDKB1;2-1 | 0.94 | 93.7 | 0.2 | 0.0 | 0.7 | 0.0 | 0.0 | 0.0 | 0.1 | 0.0 | 0.9 | 1.3 | 14.1 | 0.0 | 0.0 | 62.0 | 2.1 | 0.0 | 0.0 | 0.0 | 0.0 | 0.3 |
| 14495.15 | PcUBI | CDKB1;2-1 | 0.97 | 61.2 | 0.0 | 0.0 | 0.0 | 0.0 | 0.0 | 0.0 | 0.0 | 0.0 | 0.0 | 0.0 | 0.0 | 0.0 | 35.8 | 0.0 | 52.6 | 0.0 | 0.0 | 0.0 | 0.0 | 2.7 |
| 14495.4 | PcUBI | CDKB1;2-1 | 0.99 | 98.6 | 0.0 | 0.5 | 0.0 | 0.0 | 0.0 | 0.0 | 0.0 | 0.0 | 0.0 | 10.5 | 10.5 | 0.0 | 0.0 | 74.5 | 0.0 | 0.0 | 0.0 | 0.0 | 0.0 | 0.0 |
| 14495.17 | PcUBI | CDKB1;2-1 | 0.99 | 92.0 | 4.4 | 0.0 | 3.6 | 0.0 | 0.0 | 0.0 | 0.0 | 0.0 | 0.0 | 7.1 | 0.0 | 0.0 | 6.9 | 76.0 | 0.0 | 0.0 | 0.0 | 0.0 | 0.0 | 0.0 |
| 14495.1 | PcUBI | CDKB1;2-1 | 1.00 | 0.9 | 0.0 | 0.0 | 0.0 | 0.0 | 0.0 | 0.0 | 0.0 | 0.0 | 0.0 | 0.0 | 0.0 | 0.0 | 98.7 | 0.0 | 0.0 | 0.0 | 0.0 | 0.0 | 0.0 | 0.0 |
| NLS-GFP | | CDKB1;2-1 | 0.99 | 2.3 | 0.0 | 0.0 | 0.3 | 0.0 | 0.0 | 0.0 | 0.0 | 0.0 | 0.0 | 0.0 | 0.0 | 0.0 | 96.9 | 0.9 | 0.3 | 0.0 | 0.0 | 0.0 | 0.0 | 0.0 |

## D

| T1 Line | Promoter | Target | $R^2$ | TIDE Score (%) | -28 | -25 | -10 | -9 | -8 | -7 | -6 | -5 | -4 | -3 | -2 | -1 | 0 | 1 | 2 | 3 | 4 | 5 | 12 |
|---|---|---|---|---|---|---|---|---|---|---|---|---|---|---|---|---|---|---|---|---|---|---|---|
| 14496.1 | PcUBI | CDKA;1-1 | 0.97 | 96.5 | 17.8 | 0.0 | 17.7 | 0.0 | 0.0 | 0.0 | 8.1 | 0.0 | 0.0 | 0.0 | 0.0 | 0.0 | 0.0 | 53.0 | 0.0 | 0.0 | 0.0 | 0.0 | 0.0 |
| 14496.17 | PcUBI | CDKA;1-1 | 1.00 | 81.6 | 0.5 | 0.0 | 0.0 | 0.0 | 0.0 | 0.0 | 0.0 | 0.0 | 0.0 | 2.5 | 0.0 | 0.0 | 17.9 | 78.2 | 0.0 | 0.0 | 0.0 | 0.0 | 0.0 |
| 14496.14 | PcUBI | CDKA;1-1 | 0.96 | 77.2 | 0.0 | 0.0 | 0.0 | 0.0 | 0.0 | 0.0 | 42.5 | 0.0 | 0.0 | 0.0 | 0.0 | 0.0 | 18.6 | 34.7 | 0.0 | 0.0 | 0.0 | 0.0 | 0.0 |
| 14496.6 | PcUBI | CDKA;1-1 | 0.97 | 71.5 | 0.0 | 0.0 | 0.0 | 0.0 | 0.0 | 3.5 | 0.0 | 0.0 | 3.6 | 4.0 | 0.7 | 3.3 | 25.7 | 56.4 | 0.0 | 0.0 | 0.0 | 0.0 | 0.0 |
| 14496.12 | PcUBI | CDKA;1-1 | 0.96 | 71.0 | 0.0 | 0.0 | 0.0 | 6.2 | 0.0 | 0.0 | 0.0 | 0.0 | 0.0 | 13.1 | 0.0 | 0.0 | 24.6 | 46.1 | 0.0 | 0.0 | 5.6 | 0.0 | 0.0 |
| 14496.7 | PcUBI | CDKA;1-1 | 0.97 | 69.4 | 10.2 | 3.1 | 4.1 | 0.0 | 0.0 | 0.0 | 3.5 | 0.0 | 0.0 | 0.0 | 0.0 | 0.0 | 27.3 | 48.4 | 0.0 | 0.0 | 0.0 | 0.0 | 0.0 |
| 14496.3 | PcUBI | CDKA;1-1 | 0.99 | 23.4 | 0.0 | 0.0 | 0.0 | 0.0 | 0.0 | 0.0 | 0.0 | 0.0 | 0.0 | 0.0 | 0.0 | 0.0 | 75.6 | 0.0 | 0.0 | 0.0 | 0.0 | 0.0 | 23.4 |
| 14496.18 | PcUBI | CDKA;1-1 | 0.99 | 1.0 | 0.0 | 0.0 | 0.0 | 0.0 | 0.0 | 0.0 | 0.0 | 0.0 | 0.0 | 0.0 | 0.0 | 0.0 | 98.5 | 0.4 | 0.0 | 0.0 | 0.0 | 0.0 | 0.2 |
| 14496.20 | PcUBI | CDKA;1-1 | 1.00 | 0.9 | 0.0 | 0.0 | 0.0 | 0.0 | 0.0 | 0.0 | 0.0 | 0.0 | 0.0 | 0.0 | 0.0 | 0.0 | 98.9 | 0.1 | 0.0 | 0.1 | 0.1 | 0.2 | 0.0 |
| NLS-GFP | | CDKA;1-1 | 0.98 | 0.3 | 0.0 | 0.0 | 0.0 | 0.0 | 0.0 | 0.0 | 0.0 | 0.0 | 0.0 | 0.0 | 0.0 | 0.0 | 97.2 | 0.0 | 0.0 | 0.0 | 0.0 | 0.0 | 0.0 |

## E

| T1 Line | Promoter | Target | $R^2$ | TIDE Score (%) | -13 | -12 | -11 | -10 | -9 | -8 | -7 | -6 | -5 | -4 | -3 | -2 | -1 | 0 | 1 | 2 | 3 | 4 | 5 |
|---|---|---|---|---|---|---|---|---|---|---|---|---|---|---|---|---|---|---|---|---|---|---|---|
| 14496.1 | PcUBI | CDKB1;1-2 | 0.97 | 65.0 | 0.0 | 0.0 | 3.4 | 0.0 | 0.0 | 0.0 | 0.0 | 0.0 | 0.0 | 0.0 | 0.0 | 0.0 | 0.6 | 31.7 | 57.9 | 1.2 | 0.0 | 0.0 | 0.0 |
| 14496.17 | PcUBI | CDKB1;1-2 | 0.98 | 24.1 | 0.0 | 0.0 | 0.0 | 0.0 | 0.0 | 0.0 | 0.0 | 0.0 | 0.0 | 0.0 | 0.0 | 0.0 | 0.1 | 74.0 | 24.0 | 0.0 | 0.0 | 0.0 | 0.0 |
| 14496.14 | PcUBI | CDKB1;1-2 | 0.99 | 6.1 | 0.0 | 0.0 | 0.0 | 0.0 | 0.0 | 0.0 | 0.0 | 0.0 | 0.0 | 0.0 | 0.0 | 0.0 | 1.2 | 92.4 | 4.9 | 0.0 | 0.0 | 0.0 | 0.0 |
| 14496.6 | PcUBI | CDKB1;1-2 | 0.98 | 24.1 | 0.0 | 0.0 | 0.0 | 0.0 | 0.0 | 0.0 | 0.0 | 0.0 | 0.0 | 0.0 | 0.0 | 0.0 | 0.5 | 74.2 | 23.6 | 0.0 | 0.0 | 0.0 | 0.0 |
| 14496.12 | PcUBI | CDKB1;1-2 | 0.98 | 68.1 | 0.0 | 0.0 | 0.0 | 0.0 | 0.0 | 0.0 | 0.0 | 0.0 | 0.0 | 0.0 | 0.0 | 0.0 | 0.0 | 29.8 | 66.9 | 1.2 | 0.0 | 0.0 | 0.0 |
| 14496.7 | PcUBI | CDKB1;1-2 | 0.97 | 50.3 | 0.0 | 0.0 | 0.0 | 0.0 | 0.0 | 0.0 | 0.0 | 0.0 | 0.0 | 0.0 | 0.0 | 0.0 | 1.5 | 46.9 | 47.5 | 0.9 | 0.0 | 0.0 | 0.0 |
| 14496.3 | PcUBI | CDKB1;1-2 | 0.96 | 42.1 | 0.0 | 0.0 | 0.0 | 0.0 | 7.1 | 0.0 | 0.0 | 0.0 | 0.0 | 0.0 | 0.0 | 0.0 | 0.0 | 54.0 | 34.9 | 0.0 | 0.0 | 0.0 | 0.0 |
| 14496.18 | PcUBI | CDKB1;1-2 | 0.98 | 5.5 | 0.0 | 0.0 | 0.0 | 0.0 | 0.0 | 0.0 | 0.0 | 0.0 | 0.0 | 0.0 | 0.0 | 0.0 | 1.1 | 92.1 | 4.3 | 0.0 | 0.0 | 0.0 | 0.0 |
| 14496.20 | PcUBI | CDKB1;1-2 | 0.99 | 4.3 | 0.6 | 0.3 | 0.1 | 0.0 | 0.0 | 0.0 | 0.0 | 0.0 | 0.0 | 0.2 | 0.0 | 0.0 | 0.1 | 95.1 | 0.5 | 0.0 | 0.0 | 0.0 | 0.0 |
| NLS-GFP | | CDKB1;1-2 | 0.98 | 0.0 | 0.0 | 0.0 | 0.0 | 0.0 | 0.0 | 0.0 | 0.0 | 0.0 | 0.0 | 0.0 | 0.0 | 0.0 | 0.0 | 98.2 | 0.0 | 0.0 | 0.0 | 0.0 | 0.0 |

## F

| T1 Line | Promoter | Target | $R^2$ | TIDE Score (%) | -10 | -9 | -8 | -7 | -6 | -5 | -4 | -3 | -2 | -1 | 0 | 1 | 2 | 3 | 4 | 5 |
|---|---|---|---|---|---|---|---|---|---|---|---|---|---|---|---|---|---|---|---|---|
| 14496.1 | PcUBI | CDKB1;2-2 | 0.98 | 60.2 | 0.0 | 0.0 | 0.0 | 0.0 | 0.0 | 0.4 | 0.0 | 0.0 | 0.0 | 0.0 | 38.1 | 59.5 | 0.0 | 0.0 | 0.0 | 0.2 |
| 14496.17 | PcUBI | CDKB1;2-2 | 0.99 | 22.1 | 0.0 | 0.0 | 0.0 | 0.0 | 0.0 | 0.0 | 0.0 | 0.0 | 0.0 | 0.6 | 76.8 | 21.1 | 0.0 | 0.0 | 0.0 | 0.1 |
| 14496.14 | PcUBI | CDKB1;2-2 | 0.98 | 84.6 | 0.0 | 0.0 | 0.0 | 0.0 | 0.0 | 0.0 | 0.0 | 0.0 | 13.2 | 9.9 | 13.9 | 61.4 | 0.0 | 0.0 | 0.0 | 0.0 |
| 14496.6 | PcUBI | CDKB1;2-2 | 0.99 | 25.4 | 0.0 | 0.0 | 0.0 | 0.2 | 0.0 | 0.0 | 0.0 | 0.0 | 0.0 | 0.0 | 73.1 | 24.6 | 0.0 | 0.0 | 0.0 | 0.5 |
| 14496.12 | PcUBI | CDKB1;2-2 | 0.98 | 64.1 | 0.0 | 0.0 | 2.1 | 0.0 | 0.0 | 0.0 | 0.0 | 0.0 | 0.0 | 0.0 | 34.3 | 60.9 | 0.0 | 0.0 | 0.0 | 0.0 |
| 14496.7 | PcUBI | CDKB1;2-2 | 0.98 | 47.4 | 0.0 | 0.0 | 0.0 | 0.0 | 0.0 | 0.0 | 0.0 | 0.0 | 0.0 | 0.0 | 50.6 | 46.6 | 0.0 | 0.0 | 0.0 | 0.0 |
| 14496.3 | PcUBI | CDKB1;2-2 | 0.98 | 36.1 | 0.0 | 0.0 | 0.0 | 0.0 | 0.0 | 0.0 | 0.0 | 0.0 | 0.0 | 0.0 | 61.6 | 36.1 | 0.0 | 0.0 | 0.0 | 0.0 |
| 14496.18 | PcUBI | CDKB1;2-2 | 0.99 | 1.2 | 0.0 | 0.0 | 0.0 | 0.0 | 0.0 | 0.0 | 0.0 | 0.0 | 0.0 | 0.0 | 97.8 | 0.0 | 0.0 | 0.0 | 0.0 | 0.0 |
| 14496.20 | PcUBI | CDKB1;2-2 | 0.99 | 1.1 | 0.0 | 0.0 | 0.0 | 0.0 | 0.0 | 0.0 | 0.0 | 0.0 | 0.0 | 0.0 | 97.6 | 0.0 | 0.0 | 0.0 | 0.0 | 0.2 |
| NLS-GFP | | CDKB1;2-2 | 0.99 | 1.8 | 0.0 | 0.0 | 0.0 | 0.0 | 0.0 | 0.0 | 0.0 | 0.0 | 0.0 | 0.0 | 97.1 | 0.0 | 0.0 | 0.0 | 0.0 | 0.0 |

**Supplementary File 20 | Genotyping of individual T1 lines of *pPcUbi*:Cas9-mCherry;CDKA;1-1,CDKB1-1 and *pPcUbi*:Cas9-mCherry;CDKA;1-1,CDKB1-1**

Lines are ranked according to TIDE score of *CDKA;1-1* target. Parameters TIDE analysis: **A**, Target: *CDKA;1-1* (expected cut at 465 bp), Alignment window: 250-435, Decomposition window: 500-665, Indel size: 30. **B**, Target: *CDKB1;1-1* (expected cut at 449 bp), Alignment window: 225-399, Decomposition window: 504-630, Indel size: 50. **C**,Target: *CDKB1;2-1* (expected cut at 402 bp), Alignment window: 100-372, Decomposition window: 437-537, Indel size: 30. **D**, Target: *CDKA;1-1* (expected cut at 465 bp), Alignment window: 250-435, Decomposition window: 500-665, Indel size: 30. **E**, Target: *CDKB1;1-2* (expected cut at 486 bp), Alignment window: 250-436, Decomposition window: 521-665, Indel size: 30. **F**, Target: *CDKB1;2-2* (expected cut at 187 bp), Alignment window: 100-157, Decomposition window: 222-387, Indel size: 30. Significant values ($p<0.001$) are highlighted in green.

**Supplementary File 21**

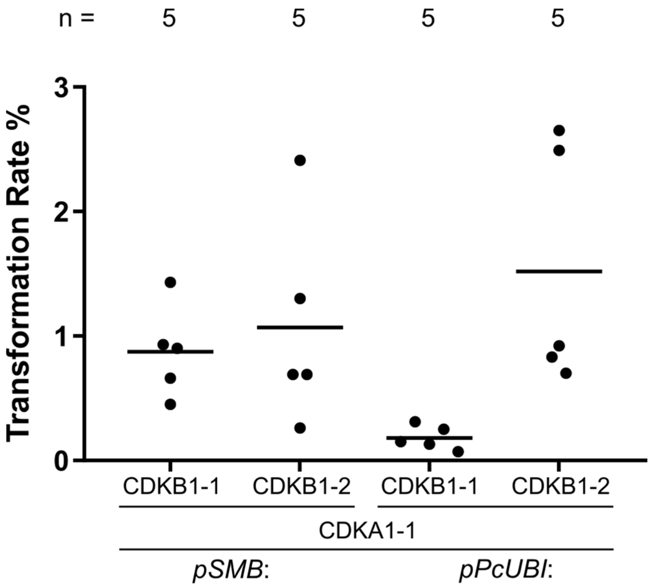

**Supplementary File 21 | Transformation rate of constructs targeting *CDKA;1*, *CDKB1;1* and *CDKB1;2***

The transformation rate was calculated as the number of FAST positive seeds in the total number of T1 seeds from individual T0 plants. The line indicates the mean. Both gRNAs CDKB1-1 and CDKB1-2 are able to simultaneously target *CDKB1;1* and *CDKB1;2*. Constructs driven by *pSMB* (pSMB:Cas9-P2A-mTagBFP2;CDKA1-1,CDKB1-1 and *pSMB*:Cas9-P2A-mTagBFP2;CDKA1-1,CDKB1-2) are used as controls for *pPcUBI* constructs (*pPcUBI*:Cas9-P2A-mCherry;CDKA1-1,CDKB1-1 and *pPcUBI*:Cas9-P2A-mCherry;CDKA1-1,CDKB1-2).

**Supplementary Table3. Oligos**

| Oligo Number | Name | Sequence | Description | Template | Reference | Method |
|---|---|---|---|---|---|---|
| **Cloning** | | | | | | |
| 45 | FWstart | TTTTGGTCTCGGGCTCGATGGATAAGAAGTACTCTATCGGACT | Cloning pGG-C-Cas9PTA-D | pGGK7_AtCas9 | unpublished | BsaI digestion - Ligation in pGGC000 |
| 431 | Cas9_nostop_REV | TTTTGGTCTCACTGAAACCTTCCTCTTCTTCTTAGGATCA | Cloning pGG-C-Cas9PTA-D | pGGK7_AtCas9 | unpublished | BsaI digestion - Ligation in pGGC000 |
| 1426 | RB54 | TTTTGGTCTCAGGCTCCATGGTGTCTAAGGGCGAAGAGCTG | Cloning pGG-C-mRuby3-D | pNCS-mRuby3 | Bajar et al. (2016) | BsaI digestion - Ligation in pGGC000 |
| 1427 | RB55 | TTTTGGTCTCACTGACTTGTACAGCTCGTCCATGCCAC | Cloning pGG-C-mRuby3-D | pNCS-mRuby3 | Bajar et al. (2016) | BsaI digestion - Ligation in pGGC000 |
| 435 | P2A_mCherry-F | TTTTGGTCTCATCAGCTGGATCTGGAGCTACTAATTTTTCTCTTCTTAAGCAAGCTGGAGATGTTGAAGAAAATCCTGGACCCATGGTGAGCAAGGGCGAGG | Cloning pGG-D-P2A-mCherry-NLS-E | pXK7-AtCas9-mCherry-NiPDS-2 | unpublished | BsaI digestion - Ligation in pGGD000 |
| 436 | mRuby3-R | TTTTGGTCTCAGCAGTTACTCTTCTTCTTGATCAGCTTCTG | Cloning pGG-D-P2A-mCherry-NLS-E | pXK7-AtCas9-mCherry-NiPDS-2 | unpublished | BsaI digestion - Ligation in pGGD000 |
| 535 | Gibson_mTagBFP2_FW | TTCTTAAGCAAGCTGGAGATGTTGAAGAAAATCCTGGACCCATGTCATCTAAGGGTGAAG | Cloning pGG-D-P2A-mTagBFP2-NLS-E | pXK7-AtCas9-mTagBFP2-NiPDS-2 | unpublished | NotI/NcoI digestion pGG-D-P2A-mCherry-NLS-E - Gibson assembly |
| 536 | Gibson_mTagBFP2_REV | TAGCTTTCCTTGCTTGCTCTTCACGCTTGAATTCAGATGCGGCCGCTGATCCAGATCCAC | Cloning pGG-D-P2A-mTagBFP2-NLS-E | pXK7-AtCas9-mTagBFP2-NiPDS-2 | unpublished | NotI/NcoI digestion pGG-D-P2A-mCherry-NLS-E - Gibson assembly |
| 581 | Gibson_GFP_GW | TTCTTAAGCAAGCTGGAGATGTTGAAGAAAATCCTGGACCCATGGTGAGCAAGGGC | Cloning pGG-D-P2A-GFP-NLS-E | pXK7-AtCas9-GFP-NiPDS-2 | unpublished | NotI/NcoI digestion pGG-D-P2A-mCherry-NLS-E - Gibson assembly |
| 536 | Gibson_mTagBFP2_REV | TAGCTTTCCTTGCTTGCTCTTCACGCTTGAATTCAGATGCGGCCGCTGATCCAGATCCAC | Cloning pGG-D-P2A-GFP-NLS-E | pXK7-AtCas9-GFP-NiPDS-2 | unpublished | NotI/NcoI digestion pGG-D-P2A-mCherry-NLS-E - Gibson assembly |
| 298 | HindIII-A-F | TCTGATCCAAGCTCAAGCTAAGCTTACCTTGAGACCGTCGAC | Cloning pFASTR-AG | pEN-L1-AG-L2 | Houbaert et al. (2018) | HindIII digestion pFASTR - Gibson assembly |
| 430 | Gibson_pFASTR_REV | GTGTTGATCTAAGTAAGGTAAGCTTATACTGAGACCGCGGCC | Cloning pFASTR-AG | pEN-L1-AG-L2 | Houbaert et al. (2018) | HindIII digestion pFASTR - Gibson assembly |
| 1428 | OLEP-AF | TTTTGGTCTCAACCTTACTTAGATCAACACATAAAAGTT | Cloning pGG-A-pOLE1-B | pFAST-G01 | Shimada et al. (2010) | BsaI digestion - Ligation in pGGA000 |
| 1429 | OLEP-BR | TTTTGGTCTCATGTTTTTTTTGTTCTTGTTTACTAGAGAG | Cloning pGG-A-pOLE1-B | pFAST-G01 | Shimada et al. (2010) | BsaI digestion - Ligation in pGGA000 |
| 1430 | SMBP-AF | TTTTGGTCTCAACCTTTGGTATTCATCCTCAATATGCAA | Cloning pGG-A-pSMB-B | pEN-L4-pSMB-XVE-R1 | Huysmans et al. (2018) | BsaI digestion - Ligation in pGGA000 |
| 1431 | SMBP-BR | TTTTGGTCTCATGTTTATCCTTACTCTTCTTTAAGCAAAC | Cloning pGG-A-pSMB-B | pEN-L4-pSMB-XVE-R1 | Huysmans et al. (2018) | BsaI digestion - Ligation in pGGA000 |
| 423 | pFAMAF | AGAAGTGAAGCTTGGTCTCAACCTATCACTAAGTGGTTTTACTAGTG | Cloning pGG-A-pFAMA-B | Genomic DNA | | BsaI digestion - Ligation in pGGA000 |
| 424 | pFAMAR | AGGGCGAGAATTCGGTCTCATGTTTGCTATTCGTGGTAGTTGATTATAAACTGC | Cloning pGG-A-pFAMA-B | Genomic DNA | | BsaI digestion - Ligation in pGGA000 |
| 425 | pTMMF | AGAAGTGAAGCTTGGTCTCAACCTGTTACTAGTAAGATGACACGTATTAC | Cloning pGG-A-pTMM-B | Genomic DNA | | BsaI digestion - Ligation in pGGA000 |
| 426 | pTMMR | AGGGCGAGAATTCGGTCTCATGTTTTCTTAGTTGTTGTTGTTGTGTGAATG | Cloning pGG-A-pTMM-B | Genomic DNA | | BsaI digestion - Ligation in pGGA000 |
| 582 | GATA23P_FW | TTTTGGTCTCAACCTCATAACTTTTCAATAATGGATCTCGG | Cloning pGG-A-pGATA23-B | GATA23 GW entry | | BsaI digestion - Ligation in pGGA000 |
| 583 | GATA23P_REV | TTTTGGTCTCATGTTCAAATAAAAAAAAACAATCTTAGTTCTTAGAG | Cloning pGG-A-pGATA23-B | GATA23 GW entry | | BsaI digestion - Ligation in pGGA000 |
| 1432 | OLEG-BF | TTTTGGTCTCAAACAATGGCGGATACAGCTAGAGGAACCCATCACGATATCATCGGCAGAGATCAGTACCCGATGATGGGCCGAGATCGAGATCAGTA | Cloning pGG-B-OLE1-C | pFAST-G01 | Shimada et al. (2010) | BsaI digestion - Ligation in pGGB000 |
| 1433 | OLEG-CR | TTTTGGTCTCAAGCCAGTAGTGTGCTGGCCACCAC | Cloning pGG-B-OLE1-C | pFAST-G01 | Shimada et al. (2010) | BsaI digestion - Ligation in pGGB000 |
| 1434 | NOST-EF | TTTTGGTCTCACTGCCTAGAGTCAAGCAGATCGTTC | Cloning pGG-E-NOST-F | PEN-R2-6-L3 | Karimi et al. (2007) | BsaI digestion - Ligation in pGGE000 |
| 1435 | NOST-FR | TTTTGGTCTCATAGTTCGATCTAGTAACATAGATGAC | Cloning pGG-E-NOST-F | PEN-R2-6-L3 | Karimi et al. (2007) | BsaI digestion - Ligation in pGGE000 |
| 345 | AtU6-26-AarI | ATTGGGATGCAGGTGGACACCTGCAGCT | Cloning pGG-F-AtU6-26-AarI-AarI-G | | | Annealed oligo's in pGG-F-AtU6-26-BbsI-BbsI-G |
| 346 | Scaffold-AarI | AAACAGCTGCAGGTGTCCACCTGCATCC | Cloning pGG-F-AtU6-26-AarI-AarI-G | | | Annealed oligo's in pGG-F-AtU6-26-BbsI-BbsI-G |
| 361 | AarIF-GlinkerIIF | ACTATAAGCTAGGTAGTAACTAGTACCTGGATGCAGGTGGACACCTGCAGCTGTATCTAGAATCGCGTATGCGAGT | Cloning pGG-F-AarI-AarI-G | | | Annealed oligo's in pGGF000 |
| 362 | AarIF-GlinkerIIR | ATACACTCGCATACGCGATTCTAGATACAGCTGCAGGTGTCCACCTGCATCCAGGTACTAGTTACTACCTAGCTTA | Cloning pGG-F-AarI-AarI-G | | | Annealed oligo's in pGGF000 |
| 195 | XbaIOLE1PF | GGCATGCGACGTCGGGCCCTCTAGTACTTAGATCAACACATAAAAGTTAG | Cloning pFASTRK24GW and pFASTGK24GW | Golden Gate mixture | | KpnI and XbaI digestion GW destination - Gibson assembly |
| 196 | KpnINOSTR | TGATCGATAATTCGCGGTACTCGATCTAGTAACATAGATGACACC | Cloning pFASTRK24GW and pFASTGK24GW | Golden Gate mixture | | KpnI and XbaI digestion GW destination - Gibson assembly |
| 298 | HindIII-A-F | TCTGATCCAAGCTCAAGCTAAGCTTACCTTGAGACCGTCGAC | Cloning Golden Gate destinations | pEN-L1-AG-L2 | Houbaert et al. (2018) | HindIII and PstI digestion GW destination - Gibson assembly |
| 313 | PstIccdBR | CCCGACGTCGCATGCCTGCAGATACTGAGACCGCGGCC | Cloning Golden Gate destinations | pEN-L1-AG-L2 | Houbaert et al. (2018) | HindIII and PstI digestion GW destination - Gibson assembly |
| 120 | SP6-univer.pr | ATTTAGGTGACACTATAG | Cloning unarmed gRNA modules | pGG-A-AtU6-PTA-B | Houbaert et al. (2018) | BsaI digestion - Gibson assembly in Empty entry vector |
| 283 | B-scaffoldR | TTTTTGGTCTCATGTTAAAAAAAGCACCGACTCGGTG | Cloning pGG-A-AtU6-26-BbsI-BbsI-B | pGG-A-AtU6-PTA-B | Houbaert et al. (2018) | BsaI digestion - Gibson assembly in pGGA000 |
| 284 | C-scaffoldR | TTTTTGGTCTCAAGCCAAAAAAAGCACCGACTCGGTG | Cloning pGG-B-AtU6-26-BbsI-BbsI-C | pGG-A-AtU6-PTA-B | Houbaert et al. (2018) | BsaI digestion - Gibson assembly in pGGB000 |
| 230 | D-scaffoldR | TTTTTGGTCTCACTGAAAAAAAAGCACCGACTCGGTG | Cloning pGG-C-AtU6-26-BbsI-BbsI-D | pGG-A-AtU6-PTA-B | Houbaert et al. (2018) | BsaI digestion - Gibson assembly in pGGC000 |
| 231 | E-scaffoldR | TTTTTGGTCTCAGCAGAAAAAAAGCACCGACTCGGTG | Cloning pGG-D-AtU6-26-BbsI-BbsI-E | pGG-A-AtU6-PTA-B | Houbaert et al. (2018) | BsaI digestion - Gibson assembly in pGGD000 |
| 232 | F-scaffoldR | TTTTTGGTCTCATAGTAAAAAAAGCACCGACTCGGTG | Cloning pGG-E-AtU6-26-BbsI-BbsI-F | pGG-A-AtU6-PTA-B | Houbaert et al. (2018) | BsaI digestion - Gibson assembly in pGGE000 |
| 233 | G-scaffoldR | TTTTTGGTCTCAATACAAAAAAAGCACCGACTCGGTG | Cloning pGG-F-AtU6-26-BbsI-BbsI-G | pGG-A-AtU6-PTA-B | Houbaert et al. (2018) | BsaI digestion - Gibson assembly in pGGF000 |
| 391 | Scaffold-BbsI-ccdBR | ACTTGCTATTTCTAGCTCTAAAACTTGTCTTCCTTATATTCCCCAGAACATCAGG | Cloning unarmed gRNA modules (*ccdB*$^{+}$) | pEN-L1-AG-L2 | Houbaert et al. (2018) | BbsI digestion of unarmed gRNA modules - Gibson assembly |
| 392 | AtU6-BbsI-ccdBF | CAGCTAGAGTCGAAGTAGTGATTGGGGTCTTCAAGCCAGATAACAGTATGCGTATTTGC | Cloning unarmed gRNA modules (*ccdB*$^{+}$) | pEN-L1-AG-L2 | Houbaert et al. (2018) | BbsI digestion of unarmed gRNA modules - Gibson assembly |
| 1436 | BsaI_ccdBCmR_F | TTTTATTGTGAGACCGCGGCCGCATTAG | PCR amplification *ccdB* and *CmR (* for replacement in vector compatible with 1 or 2 gRNA's) | pEN-L4-AG-R1 | Houbaert et al. (2018) | BsaI digestion - Ligation |
| 1437 | BsaI_ccdBCmR_R | TTTTAAACTGAGACCGTCGACTTATATTCCC | PCR amplification *ccdB* and *CmR (* for replacement in vector compatible with 1 or 2 gRNA's) | pEN-L4-AG-R1 | Houbaert et al. (2018) | BsaI digestion - Ligation |
| 23 | M13-FW | GTAAAACGACGGCCAG | PCR amplification *ccdB* and *CmR (* for replacement in vector compatible with multiple gRNA's) | pEN-L4-AG-R1 | Houbaert et al. (2018) | BsaI digestion - Ligation |
| 24 | M13-RV | CAGGAAACAGCTATGAC | PCR amplification *ccdB* and *CmR (* for replacement in vector compatible with multiple gRNA's) | pEN-L4-AG-R1 | Houbaert et al. (2018) | BsaI digestion - Ligation |
| 1879 | Gibson_A_FW_ccdB_cmR | TCCAAGCTCAAGCTAAGCTTACCTTGAGACCGCGGCCGCATTAG | PCR amplification *ccdB* and *CmR (* for replacement in expression vector with an empty promoter module) | pEN-L4-AG-R1 | Houbaert et al. (2018) | Gibson assembly |
| 1880 | Gibson_B_REV_ccdB_cmR | GGTACCAGTCGACTGAATACTGTTTGAGACCGTCGACTTATATTCCC | PCR amplification *ccdB* and *CmR (* for replacement in expression vector with an empty promoter module) | pEN-L4-AG-R1 | Houbaert et al. (2018) | Gibson assembly |
| 138 | GFP-94F | ATTGGCTGAAGCACTGCACGCCGT | Cloning gRNA target sequence *GFP-1* | | | |
| 139 | GFP-94R | AAACACGGCGTGCAGTGCTTCAGC | Cloning gRNA target sequence *GFP-1* | | | |
| 134 | GFP-35F | ATTGCAAGCAGAAGAACGGCATCA | Cloning gRNA target sequence *GFP-2* | | | |
| 135 | GFP-35R | AAACTGATGCCGTTCTTCTGCTTG | Cloning gRNA target sequence *GFP-2* | | | |
| 432 | Paired GFP-94F | TTTTGGTCTCAATTGGCTGAAGCACTGCACGCCGTGTTTTAGAGCTAGAAATAGC | Cloning gRNA target sequence *GFP-1* in position 1 dual gRNA | pEN-2xAtU6 template | | |
| 433 | Paired SMB-1R | TTTTGGTCTCAAAACGGTTCGTTCGGGTCCCAGTCCAATCACTACTTCGACTC | Cloning gRNA target sequence *SMB-1* in position 2 dual gRNA | pEN-2xAtU6 template | | |
| 434 | Paired SMB-4R | TTTTGGTCTCAAAACTGAGGAGCACGACCAGTGTACAATCACTACTTCGACTC | Cloning gRNA target sequence *SMB-2* in position 2 dual gRNA | pEN-2xAtU6 template | | |
| 437 | Paired_AtPDS-3R | TTTTGGTCTCAAAACGACCGCCGACCATGGCTGGCCAATCACTACTTCGACTC | Cloning gRNA target sequence *PDS* in position 2 dual gRNA | pEN-2xAtU6 template | | |
| 594 | ARF7-1_FW | TTTTGGTCTCAATTGGCGCTGCACAACAATCTTGGGTTTTAGAGCTAGAAATAGC | Cloning gRNA target sequence *ARF7-1* in position 1 dual gRNA | pEN-2xAtU6 template | | |
| 595 | ARF7-2_FW | TTTTGGTCTCAATTGGCGAGTGACACAAGTACTCAGTTTTAGAGCTAGAAATAGC | Cloning gRNA target sequence *ARF7-2* in position 1 dual gRNA | pEN-2xAtU6 template | | |
| 596 | ARF19-1_REV | TTTTGGTCTCAAAACTATGGATACTCGGCTTGGCCCAATCACTACTTCGACTC | Cloning gRNA target sequence *ARF19-1* in position 2 dual gRNA | pEN-2xAtU6 template | | |
| 597 | ARF19-2_REV | TTTTGGTCTCAAAACAACATCAGCTGTGATTTCTCCAATCACTACTTCGACTC | Cloning gRNA target sequence *ARF19-2* in position 2 dual gRNA | pEN-2xAtU6 template | | |
| 598 | YDA-1_FW | TTTTGGTCTCAATTGCTGTCCCCCGAAGTCCGGCAGTTTTAGAGCTAGAAATAGC | Cloning gRNA target sequence *YDA-1* in position 1 dual gRNA | pEN-2xAtU6 template | | |
| 599 | YDA-2_REV | TTTTGGTCTCAAAACCAGAACTTCCACTAGACACACAATCACTACTTCGACTC | Cloning gRNA target sequence *YDA-2* in position 2 dual gRNA | pEN-2xAtU6 template | | |
| 446 | CDKA1-FW | TTTTGGTCTCAATTGGAAGATCAGGCTAGAGCAGGGTTTTAGAGCTAGAAATAGC | Cloning gRNA target sequence *CDKA1-1* in position 1 dual gRNA | pEN-2xAtU6 template | | |
| 447 | CDKA1-REV | TTTTGGTCTCAAAACCATCCTGCAATCTGTGTAGCCAATCACTACTTCGACTC | Cloning gRNA target sequence *CDKA1-2* in position 2 dual gRNA | pEN-2xAtU6 template | | |
| 913 | CDKB_1R | TTTTGGTCTCAAAACTGGTGATTCTGAGTTTCAGCCAATCACTACTTCGACTC | Cloning gRNA target sequence *CDKB1-1* in position 2 dual gRNA | pEN-2xAtU6 template | | |
| 914 | CDKB_2R | TTTTGGTCTCAAAACTTGCTGCTCAGTTGGTGTTCCAATCACTACTTCGACTC | Cloning gRNA target sequence *CDKB1-2* in position 2 dual gRNA | pEN-2xAtU6 template | | |
| 1589 | pGGA-F | CTAGCATGGATCTCGGGCCCGGTCTCAACCTGGATGCAGGTGCACATATACC | Cloning pGG-A-AarI-SacB-AarI-B | | | |
| 1590 | pGGB-F | CTAGCATGGATCTCGGGCCCGGTCTCAAACAGGATGCAGGTGCACATATACC | Cloning pGG-B-AarI-SacB-AarI-C | | | |
| 1591 | pGGC-F | CTAGCATGGATCTCGGGCCCGGTCTCAGGCTGGATGCAGGTGCACATATACC | Cloning pGG-C-AarI-SacB-AarI-D | | | |
| 1592 | pGGD-F | CTAGCATGGATCTCGGGCCCGGTCTCATCAGGGATGCAGGTGCACATATACC | Cloning pGG-D-AarI-SacB-AarI-E | | | |
| 1593 | pGGE-F | CTAGCATGGATCTCGGGCCCGGTCTCACTGCGGATGCAGGTGCACATATACC | Cloning pGG-E-AarI-SacB-AarI-F | | | |
| 1594 | pGGF-F | CTAGCATGGATCTCGGGCCCGGTCTCAACTAGGATGCAGGTGCACATATACC | Cloning pGG-F-AarI-SacB-AarI-G | | | |
| 1595 | pGGA-R | GACACGGGCCAGAGGAGCTCGGTCTCATGTTAGCTGCAGGTGTTATTTG | Cloning pGG-A-AarI-SacB-AarI-B | | | |
| 1596 | pGGB-R | GACACGGGCCAGAGGAGCTCGGTCTCAAGCCAGCTGCAGGTGTTATTTG | Cloning pGG-B-AarI-SacB-AarI-C | | | |
| 1597 | pGGC-R | GACACGGGCCAGAGGAGCTCGGTCTCACTGAAGCTGCAGGTGTTATTTG | Cloning pGG-C-AarI-SacB-AarI-D | | | |
| 1598 | pGGD-R | GACACGGGCCAGAGGAGCTCGGTCTCAGCAGAGCTGCAGGTGTTATTTG | Cloning pGG-D-AarI-SacB-AarI-E | | | |
| 1599 | pGGE-R | GACACGGGCCAGAGGAGCTCGGTCTCATAGTAGCTGCAGGTGTTATTTG | Cloning pGG-E-AarI-SacB-AarI-F | | | |
| 1600 | pGGF-R | GACACGGGCCAGAGGAGCTCGGTCTCAATACAGCTGCAGGTGTTATTTG | Cloning pGG-F-AarI-SacB-AarI-G | | | |
| | ApaI_A_LinkerIII_E_F | CGGTCTCAACCTTAAGCTAGGTAGTAACTAGTGCGACTGAAGCTTATCGCGTATGCGAGTGCTGCTGAGACCGAGCT | Cloning pGG-A-LinkerIII-E | | | |
| | SacI_E_LinkerIII_A_R | CGGTCTCAGCAGCACTCGCATACGCGATAAGCTTCAGTCGCACTAGTTACTACCTAGCTTAAGGTTGAGACCGGGCC | Cloning pGG-A-LinkerIII-E | | | |
| | ApaI_A_LinkerIII_F_F | CGGTCTCAACCTTAAGCTAGGTAGTAACTAGTGCGACTGAAGCTTATCGCGTATGCGAGTGACTATGAGACCGAGCT | Cloning pGG-A-LinkerIII-F | | | |
| | SacI_F_LinkerIII_A_R | CGGTCTCATAGTCACTCGCATACGCGATAAGCTTCAGTCGCACTAGTTACTACCTAGCTTAAGGTTGAGACCGGGCC | Cloning pGG-A-LinkerIII-F | | | |
| | ApaI_A_LinkerIII_D_F | CGGTCTCAACCTTAAGCTAGGTAGTAACTAGTGCGACTGAAGCTTATCGCGTATGCGAGTGTCAGTGAGACCGAGCT | Cloning pGG-A-LinkerIII-D | | | |
| | SacI_D_LinkerIII_A_R | CGGTCTCACTGACACTCGCATACGCGATAAGCTTCAGTCGCACTAGTTACTACCTAGCTTAAGGTTGAGACCGGGCC | Cloning pGG-A-LinkerIII-D | | | |
| | ApaI_A_LinkerIII_B_F | CGGTCTCAACCTTAAGCTAGGTAGTAACTAGTGCGACTGAAGCTTATCGCGTATGCGAGTGAACATGAGACCGAGCT | Cloning pGG-A-LinkerIII-B | | | |
| | SacI_B_LinkerIII_A_R | CGGTCTCATGTTCACTCGCATACGCGATAAGCTTCAGTCGCACTAGTTACTACCTAGCTTAAGGTTGAGACCGGGCC | Cloning pGG-A-LinkerIII-B | | | |
| | ApaI_B_LinkerII_G_F | CGGTCTCAAACATAAGCTAGGTAGTAACTAGTGCGACTGTCTAGAATCGCGTATGCGAGTGGTATTGAGACCGAGCT | Cloning pGG-B-linkerII-G | | | |
| | SacI_G_LinkerII_B_R | CGGTCTCAATACCACTCGCATACGCGATTCTAGACAGTCGCACTAGTTACTACCTAGCTTATGTTTGAGACCGGGCC | Cloning pGG-B-linkerII-G | | | |
| | ApaI_D_LinkerII_G_F | CGGTCTCATCAGTAAGCTAGGTAGTAACTAGTGCGACTGTCTAGAATCGCGTATGCGAGTGGTATTGAGACCGAGCT | Cloning pGG-D-linkerII-G | | | |
| | SacI_G_LinkerII_D_R | CGGTCTCAATACCACTCGCATACGCGATTCTAGACAGTCGCACTAGTTACTACCTAGCTTACTGATGAGACCGGGCC | Cloning pGG-D-linkerII-G | | | |
| | ApaI_F_LinkerII_G_F | CGGTCTCAACTATAAGCTAGGTAGTAACTAGTGCGACTGTCTAGAATCGCGTATGCGAGTGGTATTGAGACCGAGCT | Cloning pGG-F-linkerII-G | | | |
| | SacI_G_LinkerII_F_R | CGGTCTCAATACCACTCGCATACGCGATTCTAGACAGTCGCACTAGTTACTACCTAGCTTATAGTTGAGACCGGGCC | Cloning pGG-F-linkerII-G | | | |
| | RB42 | TTTTGGTCTCAGGCTCCATGTCATCTAAGGGTGAAGAGCTTA | Cloning pGG-C-mTagBFP2-D | | | |
| | RB43 | TTTTGGTCTCACTGATCCACCTCCCAGTCCTGCGTAT | Cloning pGG-C-mTagBFP2-D | | | |

**Supplementary Table3. Oligos**

| Oligo Number | Name | Sequence | Description | Template | Reference | Method |
|---|---|---|---|---|---|---|
| **Vector validation** | | | | | | |
| 61 | GG_seq_fw | GTGAGCGGATAACAATTTCACA | Sequencing Golden Gate Entry | | | |
| 62 | GG_seq_rev | CGACGGCCAGGTAATACGACT | Sequencing Golden Gate Entry | | | |
| 81 | Cas9PTA_int1F | AACTTCAAGTCTAACTTCGATCTC | Sequencing Cas9 | | | |
| 82 | Cas9PTA_int2F | AAAGTCTGAGGAAACCATCACC | Sequencing Cas9 | | | |
| 83 | Cas9PTA_int3F | TCTCTCACCTTTAAAGAGGATATC | Sequencing Cas9 | | | |
| 84 | Cas9PTA_int4F | CCTCGATTCTAGGATGAACACC | Sequencing Cas9 | | | |
| 85 | Cas9PTA_int5F | TATCATGGAAAGGTCATCTTTCGA | Sequencing Cas9 | | | |
| 169 | pGG3838 | GGCATTTCAGTCAGTTGCTCAATG | Sequencing pFASTR A-G | | | |
| 170 | pGG4468 | GGACTTGTGCGACATGTCGTTTTC | Sequencing pFASTR A-G and pFASTR-BsaI-CmR-ccdB-BsaI-Cas9-P2A-mCherry-G7T-AtU6-GFP-1 | | | |
| 342 | Ole1P-204R | GAGCTAGTTAGAATCTCAGGCCTTG | Sequencing pFASTR A-G | | | |
| 427 | pFAMA505F | CTCCTATGGATCCGCGACTC | Sequencing pGG-A-pFAMA-B | | | |
| 428 | pFAMA1102F | TAAGCTAGGGAGTTGCTCTAGATCC | Sequencing pGG-A-pFAMA-B | | | |
| 429 | pFAMA1815F | GGGTCACGAATGTGTACCATTCAC | Sequencing pGG-A-pFAMA-B | | | |
| 638 | SacB seq | GTTCCTGAGTTCGATTCGTCCAC | Sequencing pGG-F-A-AarI-SacB-AarI-G-G | | | |
| 330 | G7TF | CGCACTCAGTCTTTCATCTACGG | Sequencing of Expression plasmids | | | |
| 342 | Ole1P-204R | GAGCTAGTTAGAATCTCAGGCCTTG | Sequencing of Expression plasmids | | | |
| 527 | Cas9 REV | CCTTGAACTTCTTAGATGGCACC | Sequencing pFASTR-BsaI-CmR-ccdB-BsaI-Cas9-P2A-mCherry-G7T-AtU6-GFP-1 | | | |
| 20 | SS42 | TCCCAGGATTAGAATGATTAGG | Colony PCR | | | |
| 139 | GFP-94R | AAACACGGCGTGCAGTGCTTCAGC | Colony PCR | | | |
| 1658 | gentF2 | CGGCGGATTTGTCCTACTCA | | | | |
| **Genotyping** | | | | | | |
| 1210 | GFP F | TTGCCGTCCTCCTTGAAGTC | Genotyping *GFP* | | | |
| 1211 | NLS R | GCTCCATGCAGCCTTCTCTT | Genotyping *GFP* | | | |
| 101 | AtPDS FW2241 | GACGTCAGGAAGAACATGGTC | Genotyping *PDS* | | | |
| 102 | AtPDS REV2875 | AACACCTCGTCGGTCACGC | Genotyping *PDS* | | | |
| 1085 | SMB 1,256 F | CTTATACGTGTGTGTGCGCG | Genotyping *SMB* | | | |
| 1086 | SMB 2,234 R | GCAGCTGAGTCAGGGCTAAA | Genotyping *SMB* | | | |
| 1108 | YDA 337 F | CCTCGTCCTATTGTCCGTCC | Genotyping *YDA* | | | |
| 1109 | YDA_1,609 R | ACTGAGATTTGCCCCTGAGA | Genotyping *YDA* | | | |
| 1104 | ARF7_2,873 F | GGAATTTCCGGTGCTCTTGC | Genotyping *ARF7-1* | | | |
| 1105 | ARF7_3,509 R | GCTGTGTTCTTAGCTGCTGC | Genotyping *ARF7-1* | | | |
| 1079 | ARF7_1,388 F | GGCTGATCCTGAGACGGATG | Genotyping *ARF7-2* | | | |
| 1080 | ARF7_2,143 R | GCACGTCTTATCCCCAACAGAA | Genotyping *ARF7-2* | | | |
| 1106 | ARF19_1,562 F | TCTGATCTCAGGCCAACCAA | Genotyping *ARF19* | | | |
| 1107 | ARF19_2,451 R | TCCCATCCAAGGCATTGCTC | Genotyping *ARF19* | | | |
| 866 | CDKA_F | GCTCTGGATTCAGCTGGCTT | Genotyping *CDKA1* | | | |
| 867 | CDKA_R | CTTCAGTGAGTTTGTGCGGC | Genotyping *CDKA1* | | | |
| 897 | CDKB1;1_F | TGTTCCTCTTAAGTCTTATACG | Genotyping *CDKB1;1* | | | |
| 898 | CDKB1;1_R | CCAAGCAATTCAGCTTAGGAAA | Genotyping *CDKB1;1* | | | |
| 899 | CDKB1;2_F | TGTGCCTCTTAAGGCTTATACA | Genotyping *CDKB1;2* | | | |
| 900 | CDKB1;2_R | AGGAACGAACCCCAACAGAA | Genotyping *CDKB1;2* | | | |

**Supplementary Table3. Oligos**

| Oligo Number | Name | Sequence | Description | Template | Reference | Method |
|---|---|---|---|---|---|---|
| **Vector validation** | | | | | | |
| 61 | GG_seq_fw | GTGAGCGGATAACAATTTCACA | Sequencing Golden Gate Entry | | | |
| 62 | GG_seq_rev | CGACGGCCAGGTAATACGACT | Sequencing Golden Gate Entry | | | |
| 81 | Cas9PTA_int1F | AACTTCAAGTCTAACTTCGATCTC | Sequencing Cas9 | | | |
| 82 | Cas9PTA_int2F | AAAGTCTGAGGAAACCATCACC | Sequencing Cas9 | | | |
| 83 | Cas9PTA_int3F | TCTCTCACCTTTAAAGAGGATATC | Sequencing Cas9 | | | |
| 84 | Cas9PTA_int4F | CCTCGATTCTAGGATGAACACC | Sequencing Cas9 | | | |
| 85 | Cas9PTA_int5F | TATCATGGAAAGGTCATCTTTCGA | Sequencing Cas9 | | | |
| 169 | pGG3838 | GGCATTTCAGTCAGTTGCTCAATG | Sequencing pFASTR A-G | | | |
| 170 | pGG4468 | GGACTTGTGCGACATGTCGTTTTC | Sequencing pFASTR A-G and pFASTR-BsaI-CmR-ccdB-BsaI-Cas9-P2A-mCherry-G7T-AtU6-GFP-1 | | | |
| 342 | Ole1P-204R | GAGCTAGTTAGAATCTCAGGCCTTG | Sequencing pFASTR A-G | | | |
| 427 | pFAMA505F | CTCCTATGGATCCGCGACTC | Sequencing pGG-A-pFAMA-B | | | |
| 428 | pFAMA1102F | TAAGCTAGGGAGTTGCTCTAGATCC | Sequencing pGG-A-pFAMA-B | | | |
| 429 | pFAMA1815F | GGGTCACGAATGTGTACCATTCAC | Sequencing pGG-A-pFAMA-B | | | |
| 638 | SacB_seq | GTTCCTGAGTTCGATTCGTCCAC | Sequencing pGG-F-A-AarI-SacB-AarI-G-G | | | |
| 330 | G7TF | CGCACTCAGTCTTTCATCTACGG | Sequencing of Expression plasmids | | | |
| 342 | Ole1P-204R | GAGCTAGTTAGAATCTCAGGCCTTG | Sequencing of Expression plasmids | | | |
| 527 | Cas9_REV | CCTTGAACTTCTTAGATGGCACC | Sequencing pFASTR-BsaI-CmR-ccdB-BsaI-Cas9-P2A-mCherry-G7T-AtU6-GFP-1 | | | |
| 20 | SS42 | TCCCAGGATTAGAATGATTAGG | Colony PCR | | | |
| 139 | GFP-94R | AAACACGGCGTGCAGTGCTTCAGC | Colony PCR | | | |
| 1658 | gentF2 | CGGCGGATTTGTCCTACTCA | | | | |
| **Genotyping** | | | | | | |
| 1210 | GFP_F | TTGCCGTCCTCCTTGAAGTC | Genotyping *GFP* | | | |
| 1211 | NLS_R | GCTCCATGCAGCCTTCTCTT | Genotyping *GFP* | | | |
| 101 | AtPDS_FW2241 | GACGTCAGGAAGAACATGGTC | Genotyping *PDS* | | | |
| 102 | AtPDS_REV2875 | AACACCTCGTCGGTCACGC | Genotyping *PDS* | | | |
| 1085 | SMB_1,256 F | CTTATACGTGTGTGTGCGCG | Genotyping *SMB* | | | |
| 1086 | SMB_2,234 R | GCAGCTGAGTCAGGGCTAAA | Genotyping *SMB* | | | |
| 1108 | YDA_337 F | CCTCGTCCTATTGTCCGTCC | Genotyping *YDA* | | | |
| 1109 | YDA_1,609 R | ACTGAGATTTGCCCCTGAGA | Genotyping *YDA* | | | |
| 1104 | ARF7_2,873 F | GGAATTTCCGGTGCTCTTGC | Genotyping *ARF7-1* | | | |
| 1105 | ARF7_3,509 R | GCTGTGTTCTTAGCTGCTGC | Genotyping *ARF7-1* | | | |
| 1079 | ARF7_1,388 F | GGCTGATCCTGAGACGGATG | Genotyping *ARF7-2* | | | |
| 1080 | ARF7_2,143 R | GCACGTCTTATCCCCAACAGAA | Genotyping *ARF7-2* | | | |
| 1106 | ARF19_1,562 F | TCTGATCTCAGGCCAACCAA | Genotyping *ARF19* | | | |
| 1107 | ARF19_2,451 R | TCCCATCCAAGGCATTGCTC | Genotyping *ARF19* | | | |
| 866 | CDKA_F | GCTCTGGATTCAGCTGGCTT | Genotyping *CDKA1* | | | |
| 867 | CDKA_R | CTTCAGTGAGTTTGTGCGGC | Genotyping *CDKA1* | | | |
| 897 | CDKB1;1_F | TGTTCCTCTTAAGTCTTATACG | Genotyping *CDKB1;1* | | | |
| 898 | CDKB1;1_R | CCAAGCAATTCAGCTTAGGAAA | Genotyping *CDKB1;1* | | | |
| 899 | CDKB1;2_F | TGTGCCTCTTAAGGCTTATACA | Genotyping *CDKB1;2* | | | |
| 900 | CDKB1;2_R | AGGAACGAACCCCAACAGAA | Genotyping *CDKB1;2* | | | |

#### Supplementary File 23

| Target Gene | Target Sequence |
|---|---|
| *GFP-1* | GCTGAAGCACTGCACGCCGT**AGG** |
| *GFP-2* | CAAGCAGAAGAACGGCATCA**AGG** |
| *PDS3* | GCCAGCCATGGTCGGCGGTC**AGG** |
| *SMB-1* | GACTGGGACCCGAACGAACC**GGG** |
| *SMB-2* | TACACTGGTCGTGCTCCTCA**TGG** |
| *YDA-1* | TGTGTCTAGTGGAAGTTCTG**TGG** |
| *YDA-2* | CTGTCCCCCGAAGTCCGGCA**AGG** |
| *ARF7-1* | GCGCTGCACAACAATCTTGG**CGG** |
| *ARF7-2* | GCGAGTGACACAAGTACTCA**CGG** |
| *ARF19-1* | GGCCAAGCCGAGTATCCATA**TGG** |
| *ARF19-2* | GAGAAATCACAGCTGATGTT**GGG** |
| *CDKA;1-1* | GAAGATCAGGCTAGAGCAGG**AGG** |
| *CDKA;1-2* | GCTACACAGATTGCAGGATG**TGG** |
| *CDKB1;1-1* and *CDKB1;2-1* | GCTGAAACTCAGAATCACCA**GGG** |
| *CDKB1;1-2* and *CDKB1;2-2* | GAACACCAACTGAGCAGCAA**TGG** |

**Supplementary File 23 | gRNA target sequences**

The 23-nt target sequences are given for each gRNA. PAM sequences are in bold.

#### Supplementary File 24

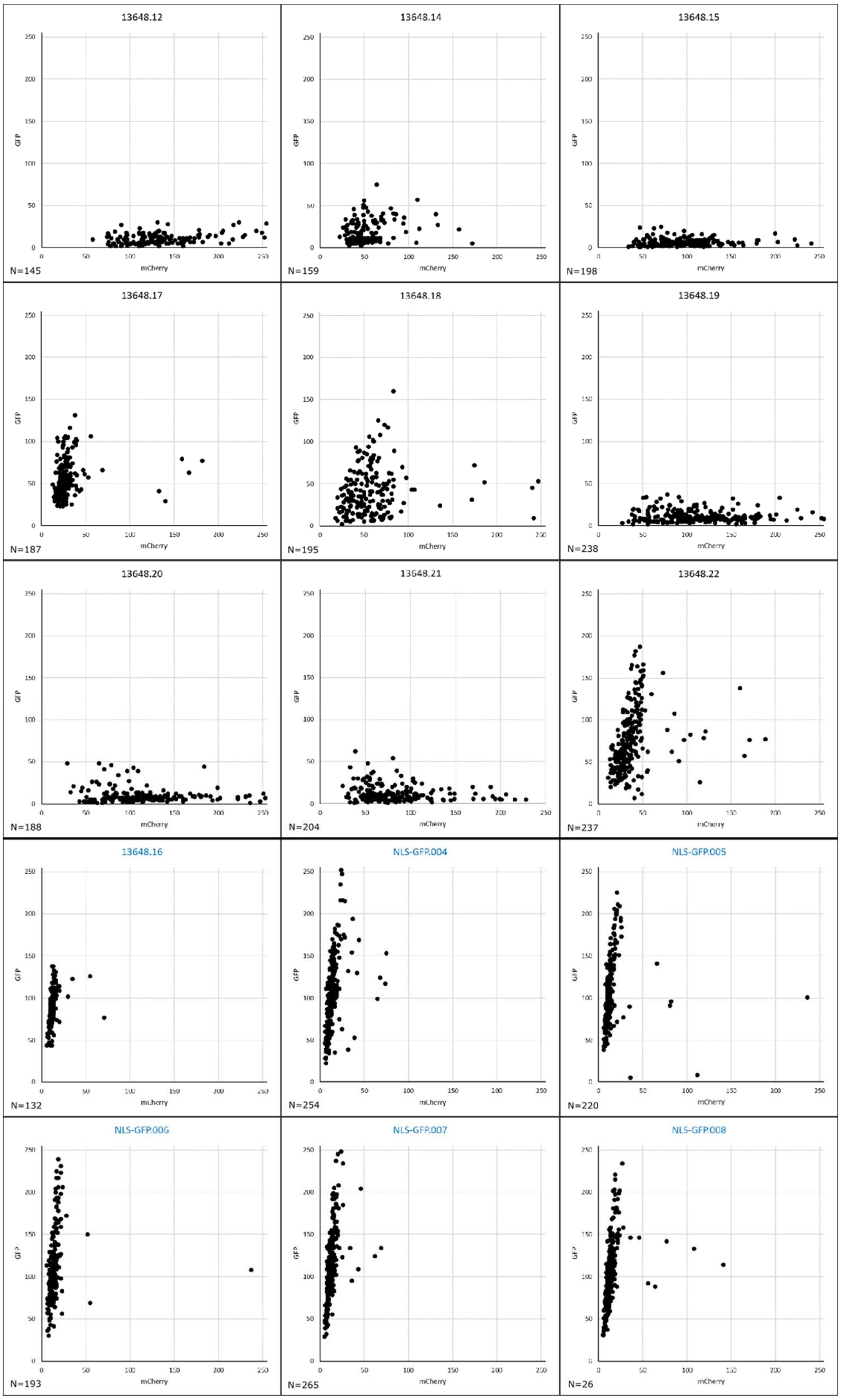

**Supplementary File 24 | Signal intensity values for GFP and mCherry in root cap nuclei**

Values for individual root cap nuclei are plotted for *pSMB:*Cas9-mCherry;GFP1 T1 and NLS-GFP seedlings. Plant lines in blue are used as reference for no GFP mutation and were not used for correlation analysis in Figure 2C.
