## Supplementary Methods for "CRISPR-TSKO facilitates efficient cell type-, tissue-, or organ-specific mutagenesis in Arabidopsis"

**Materials**

| **Product** | **Concentration** | **Provider** |
|---|---|---|
| CutSmart Buffer | | New England Biolabs |
| T4 DNA ligase | 400,000 U/mL | New England Biolabs |
| BsaI-HF®v2 | 20,000 U/mL | New England Biolabs |
| BbsI-HF | 20,000 U/mL | New England Biolabs |
| ATP | 10 mM | ThermoFisher Scientific |
| NheI | 10 U/mL | Promega Corporation |
| Buffer B | 10X | Promega Corporation |
| Acetylated BSA | 10 µg/µL | Promega Corporation |
| AarI | 2 U/mL | ThermoFisher Scientific |
| Buffer AarI | 10X | ThermoFisher Scientific |
| Oligonucleotides | 50X (0.025 mM) | ThermoFisher Scientific |
| Q5® High-Fidelity DNA Polymerase | 2,000 U/mL | New England Biolabs |
| Q5® Reaction Buffer | 5X | New England Biolabs |
| 10 mM dNTP's | 10 mM | New England Biolabs |
| One Shot™ ccdB Survival™ 2 T1R Competent Cells | | ThermoFisher Scientific |
| ZymocleanTM Gel DNA Recovery Kit | | Zymo Research |

**Primers**

| | |
|---|---|
| Oligo 23 | GTAAAACGACGGCCAG |
| Oligo 24 | CAGGAAACAGCTATGAC |
| Oligo 1436 | TTTTATTGTGAGACCGCGGCCGCATTAG |
| Oligo 1437 | TTTTAAACTGAGACCGTCGACTTATATTCCC |

### I. Golden Gate Assembly of One-step Cloning vectors

#### A. *Golden Gate*

The following components are mixed in a PCR tube:

| **Component** | | **Amount** | **Volume (µL)** |
|---|---|---|---|
| A-B entry vector | (Tissue-specific) Promotor | 100 ng | 1 |
| B-C entry vector | N-tag or Linker | 100 ng | 1 |
| C-D entry vector | Nuclease | 100 ng | 1 |
| D-E entry vector | C-tag or Linker | 100 ng | 1 |
| E-F entry vector | Terminator | 100 ng | 1 |
| F-G entry vector | **Variable*** | 100 ng | 1 |
| Destination vector | | 100 ng | 1 |
| 10X CutSmart Buffer | | 1X | 1.5 |
| 10 mM ATP | | 1 mM | 1.5 |
| T4 DNA ligase | | 200 U | 0.5 |
| BsaI-HF®v2 | | 10 U | 0.5 |
| MQ | | | 4 |
| | | | 15 |

* The F-G entry vector used depends on your final goal (cloning one, two, or multiple gRNAs). For cloning a one-step cloning vector compatible with 1 or 2 gRNAs, make use of the unarmed gRNA module pGG-F-AtU6-26-AarI-AarI-G. For cloning a vector compatible with multiple gRNAs, make use of the linker pGG-F-AarI-SacB-AarI-G.

Golden Gate reaction conditions:

| | | |
|---|---|---|
| 37°C | 3 min | 30 x |
| 16°C | 3 min | |
| 50°C | 5 min | |
| 80°C | 5 min | |
| 16°C | ∞ | |

Five µL of the reaction mixture are transformed into 50 µL *ccdB*-sensitive DH5α *E. coli* cells via heat shock. The transformed cells are plated on LB medium containing 100 µg/mL spectinomycin. The vector can be validated using colony PCR and restriction digest.

Note: The one-step cloning vectors compatible with multiple gRNAs contain the *SacB* gene. This gene encodes an enzyme that converts sucrose to levan, which accumulates in the periplasm and is toxic to E. coli. This allows for counter selection in subsequent cloning on LB medium containing 10% sucrose.

B. *AarI restriction digest*

AarI digestion is performed according to the manufacturer's recommendations.

| Component | Amount | Volume |
|---|---|---|
| Plasmid DNA (100 ng/µL) | 1 µg | 10.0 µL |
| MQ | | 7.1 µL |
| Buffer AarI | 1x | 2.0 µL |
| Oligonucleotides | 0.5 µM | 0.4 µL |
| AarI | 1 U | 0.5 µL |
| | | 20.0 µL |

Mix gently and spin down for a few seconds, before incubating at 37°C for 1-16 hours. Heat inactivate the AarI enzyme for 20 min at 65°C, or directly run on a 0.8% agarose gel. Include the undigested vector as a control.

Expected bands:

- The one-step cloning vectors compatible with cloning one or two gRNAs generates two fragments. The smallest fragment (32 bp) is too small to be visualized. The largest fragment is shifted upwards compared to the undigested vector.
- The one step cloning vectors compatible with cloning multiple gRNAs generates two fragments. The fragment containing the *SacB* selectable marker has a size of 1898 base pairs. The largest fragment has a size of > 10 kb.

Gel purify the largest fragment

C. *Replacement of AarI restriction sites for BsaI restriction sites*

A fragment containing the *ccdB* and *CmR* selectable markers flanked with BsaI sites, is PCR amplified with Q5® High-Fidelity DNA Polymerase from the vector pEN-L4-A-G-R1. The primers for this reaction depend on the vector that you are cloning.

- For cloning a one-step vector compatible with 1 or 2 gRNAs, the PCR reaction is performed with oligos 1436 & 1437 (resulting in a fragment of 1451 bp). Following BsaI digestion, this fragment will have the ATTG and AAAC overlaps corresponding to the AtU6-26 promoter and SpCas9 scaffold sequences, respectively.
- For cloning a vector compatible with multiple gRNAs, the PCR reaction is performed with oligo 23 & 24 (resulting in a fragment of 1834 bp). Following BsaI digestion, this fragment will have the ACCT and ATAC overhangs corresponding to the A and G sites in the GreenGate cloning system.

Mix the following components in a PCR tube:

| **Component** | **Amount** | **Volume (µL)** |
|---|---|---|
| pEN-L4-A-G-R1 (100 ng/µL) | 50 ng | 0.5 |
| FW primer (10 µM) | 0.5 µM | 2.5 |
| REV primer (10 µM) | 0.5 µM | 2.5 |
| dNTP's | 200 µM | 1 |
| Q5® Reaction Buffer | 1X | 10 |
| Q5® High-Fidelity DNA Polymerase | 1U | 0.5 |
| MQ | | 33 |
| | | 50* |

Reaction conditions:

| | | |
|---|---|---|
| 98°C | 30 sec | |
| 98°C | 10 sec | 34 x |
| $T_a$** | 30 sec | |
| 72°C | 1 min | |
| 72°C | 2 min | |
| 16°C | ∞ | |

* The manufacturer's recommended final volume is 50 µL. This can however be lowered to 10 µL to reduce costs.

** 1436 + 1437 ($T_a$ = 67°C) and 23 + 24 ($T_a$= 57°C), as calculated by the NEB Tm Calculator (http://tmcalculator.neb.com/#!/main)

Purify from gel and digest with BsaI

| Component | Amount | Volume (µL) |
|---|---|---|
| PCR product | 1 µg | |
| 10X CutSmart buffer | 1X | 5 |
| BsaI-HF®v2 | 10 U | 0.5 |
| MQ | | |
| | | 50 |

Mix gently and spin down for a few seconds. Incubate at 37°C for at least 1 hour, and heat inactivate the BsaI enzyme for 20 min at 80°C.

The following components are mixed in a microcentrifuge tube:

| Component | Amount | Volume (µL) |
|---|---|---|
| AarI digested vector | | 1 |
| BsaI digested PCR product | | 1 |
| T4 DNA Ligase | 400 U | 1 |
| 10X T4 DNA Ligase buffer | 1X | 1 |
| MQ | | 6 |
| | | 10 |

Gently mix the reaction and microfuge briefly. According to the manufacturer's recommendation incubate at 16°C overnight or at room temperature for 10 minutes. We however generally incubate at least for one hour (at room temperature). Heat inactivate at 65°C for 10 minutes. Chill on ice and transform 1-5 µL of the reaction mixture into 50 µL One Shot™ *ccd*B Survival™ 2 T1R Competent Cells via heat shock. Transformed cells are selected on LB medium containing 25 µg/µL chloramphenicol or on LB medium containing 10% sucrose. The inserted fragment is verified via Sanger sequencing, and the vector can be validated by colony PCR and restriction digest (NheI is recommended).

### II. Vectors compatible with cloning one or two gRNAs

Loading the vectors can be done with either annealed oligos for a single gRNA or with a PCR product for two gRNAs.

A. Single gRNA: Annealed Oligos

- FW: 5'-ATTG-$N_{20}$ with $N_{20}$ being the protospacer of the gRNA
- REV: 5'-AAAC-$N_{20}$ with $N_{20}$ being the reverse complement of the protospacer

```
5'- ATTGNNNNNNNNNNNNNNNNNNNN
        NNNNNNNNNNNNNNNNNNNNCAAA -5'
```

Protocol:

- Add 1 µL of each 100 µM oligo to 48 µL of MQ water
- Incubate with a slow-cooling program on the thermal cycler e.g.: 5 minutes at 95°C; 95-85°C, -2°C/second; 85-25°C, -0.1°C/second
- The annealed oligos can be cloned in the destination vector via a Golden Gate reaction

| Component | Amount | Volume (µL) |
|---|---|---|
| Annealed oligos | 100 ng | 1 |
| Destination vector | 100 ng | 1 |
| 10X CutSmart Buffer | 1X | 1.5 |
| 10 mM ATP | 1 mM | 1.5 |
| T4 DNA ligase | 200 U | 0.5 |
| BsaI-HF®v2 | 5 U | 0.5 |
| MQ | | 9 |
| | | 15 |

| | | |
|---|---|---|
| 37°C | 3 min | 30 x |
| 16°C | 3 min | |
| 50°C | 5 min | |
| 80°C | 5 min | |
| 16°C | ∞ | |

Transform 5 µL of the reaction mixture via heat shock into 50 µL *ccdB*-sensitive DH5α *E. coli* cells, and plate on LB medium containing 100 µg/mL Spectinomycin. The vectors can be validated by Sanger sequencing and restriction digest with NheI.

B. Two gRNAs: PCR product

For cloning two gRNAs in the destination vector we used the following approach as described by (Xing et al., 2014) with some modifications. **BsaI sites** in bold.

- FW: 5'-TTTT**GGTCTC**AATTG-$N_{20}$-GTTTTAGAGCTAGAAATAGC with $N_{20}$ being the protospacer of the first gRNA
- REV: 5'-TTTT**GGTCTC**AAAAC-$N_{20}$-CAATCACTACTTCGACTC with $N_{20}$ being the reverse complement of the protospacer of the second gRNA

| **Component** | **Amount** | **Volume (µL)** |
|---|---|---|
| pEN-2xAtU6 template | 50 ng* | 0.5 |
| 10 µM FW primer | 0.5 µM | 2.5 |
| 10 µM REV primer | 0.5 µM | 2.5 |
| 10 mM dNTP's | 200 µM | 1 |
| 5X Q5® Reaction Buffer | 1X | 10 |
| Q5® High-Fidelity DNA Polymerase | 0.02 U/µl | 0.5 |
| MQ | | 33 |
| | | 50** |

| | | |
|---|---|---|
| 98°C | 30 sec | |
| 98°C | 10 sec | 34 x |
| 55°C*** | 15 sec | |
| 72°C | 15 sec | |
| 72°C | 2 min | |
| 16°C | ∞ | |

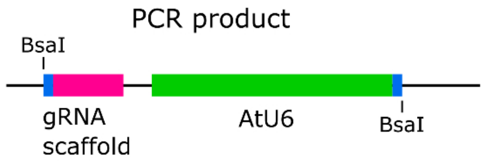


* This is the amount of plasmid used for this report. We routinely use less (≤1ng).

** The manufacturer's recommended final volume is 50 µL. This can however be lowered to reduce the cost.

*** As calculated by the NEB Tm Calculator (http://tmcalculator.neb.com/#!/main) using the invariable 3' sequence of the primer.

5-10 µL of PCR products are checked on an 0,8% agarose gel and should give rise to a band of 575 bp. The remainder of PCR products are column purified and cloned in the destination vector via a Golden Gate reaction. If primer dimers are observed in the gel, gel purification is recommended.

The following components are mixed in a PCR tube:

| **Component** | **Amount** | **Volume (µL)** |
|---|---|---|
| PCR product | ~10 ng | 1 |
| Destination vector | 100 ng | 1 |
| 10X CutSmart Buffer | 1X | 1.5 |
| 10 mM ATP | 1 mM | 1.5 |
| T4 DNA ligase | 200 U | 0.5 |
| BsaI-HF®v2 | 5 U | 0.5 |
| MQ | | 9 |
| | | 15 |

| | | |
|---|---|---|
| 37°C | 3 min | 30 x |
| 16°C | 3 min | |
| 50°C | 5 min | |
| 80°C | 5 min | |
| 16°C | ∞ | |

Five µL of the reaction mixture is transformed via heat shock into 50 µL *ccdB*-sensitive DH5α *E. coli* cells. The cells are plated on LB medium containing 100 µg/mL spectinomycin. Plasmids are validated with a restriction digest and the inserted fragment verified via Sanger sequencing.

### III. Multiple gRNA vectors

The unarmed gRNA modules (*ccdB*$^{+}$) can be loaded with either one or two gRNAs.

The loading of a single gRNA can be done as described in **II.** A., with some modifications. The Golden Gate assembly of the annealed oligos is done with BbsI-HF, transformed into *ccdB*-sensitive DH5α *E. coli* cells and plated on LB medium containing 100 µg/mL Carbenicillin.

For cloning two gRNAs in the unarmed gRNA modules, the loading of two gRNAs can be done essentially as described in **II.**B, with some modifications. Use the following primer sequences. **BbsI sites** in bold.

- FW: 5'- TTTT**GAAGAC**ATATTG-$N_{20}$-GTTTTAGAGCTAGAAATAGC with $N_{20}$ being the protospacer of the first gRNA
- REV: 5'- TTTT**GAAGAC**TTAAAC-$N_{20}$- CAATCACTACTTCGACTC with $N_{20}$ being the reverse complement of the protospacer of the second gRNA

PCR products are checked on an 0,8% agarose gel and should give rise to a band of 577 bp. The remainder of the PCR product are column purified. Golden Gate assembly is done with BbsI-HF, transformed into *ccdB*-sensitive DH5α *E. coli* cells and plated on LB medium containing 100 µg/mL Carbenicillin.

Confirm gRNA clones by Sanger sequencing.

These armed gRNA entry modules can be combined in a destination vector via a Golden Gate reaction. One or more linkers (**Supplemental Table 1**) can be used, to span the unused modules. The following components are mixed in a PCR tube:

| **Component** | **Amount** | **Volume (µL)** |
|---|---|---|
| A-B armed gRNA module | 100 ng | 1 |
| B-C armed gRNA module | 100 ng | 1 |
| C-D armed gRNA module | 100 ng | 1 |
| D-E armed gRNA module | 100 ng | 1 |
| E-F armed gRNA module | 100 ng | 1 |
| F-G armed gRNA module | 100 ng | 1 |
| Destination vector | 100 ng | 1 |
| 10X CutSmart Buffer | 1x | 1.5 |
| 10 mM ATP | 1 mM | 1.5 |
| T4 DNA ligase | 200 U | 0.5 |
| BsaI-HF®v2 | 10 U | 0.5 |
| MQ | | 4 |
| | | 15 |

| | | |
|---|---|---|
| 37°C | 3 min | 30 x |
| 16°C | 3 min | |
| 50°C | 5 min | |
| 80°C | 5 min | |
| 16°C | ∞ | |

Five µL of the reaction mixture are transformed into 50 µL *ccdB*-sensitive DH5α *E. coli* cells via heat shock. The transformed cells are plated on LB medium containing 100 µg/mL spectinomycin. The vector can be validated using colony PCR and restriction digest. Sanger sequencing is not recommended as the arrays of gRNAs usually cause the reactions to fail.
